# Modeling the Controls on Microbial Iron and Manganese Reduction in Methanic Sediments

**DOI:** 10.1101/2025.03.30.646147

**Authors:** Racheli Neumann Wallheimer, Itay Halevy, Orit Sivan

## Abstract

Microbial iron and manganese respiration processes have been observed in deep methanic sediments of lacustrine and marine environments, challenging the “classical” model of microbial respiration in aquatic systems. Nonetheless, assessments of the type and relative role of these respiration processes in the methanic zone are lacking. Here, we quantify both the thermodynamic and the kinetic controls of potential iron and manganese respiration processes in the diffusive controlled steady state methanic sediments of lacustrine and marine sites – Lake Kinneret (LK) and the Southeastern Mediterranean Sea (MedS). We consider the substrates (electron donors) and iron and manganese oxides (electron acceptors) at concentrations that have been measured at these sites. Using theoretical bioenergetic methods, we develop a nominal model to calculate catabolic rates, considering both kinetic and thermodynamic parameters. Then, we estimate the biomass growth rates from the catabolic rates, the energy generated in each reduction-oxidation (redox) reaction, the biomass yield from a given amount of energy, the number of cells participating in each reaction, and the energetic needs of the cells. Lastly, we estimate the microbial community sizes of expected iron and manganese reducers. Additionally, we perform a Monte Carlo simulation to account for variations in uncertain parameter values, along with a sensitivity analysis. Together, these calculations enable estimation of the expected total reaction rates of the various metabolic processes.

Our results indicate that the type of iron or manganese oxide, which determines its thermodynamic and kinetic properties, is more significant in influencing bioreaction rates than its concentration. Thus, bioreactions with amorphous manganese oxides are more favorable than those with highly reactive iron oxides. Among the iron oxides, the reduction of amorphous iron oxyhydroxide and ferrihydrite are the only reactions capable of generating biomass in the methanic sediments at both sites. In both environments, manganese oxide reduction by ammonium and methane oxidation are expected to be significant, while manganese oxide reduction by hydrogen and acetate oxidation are expected to be considerable only in LK. The most probable iron oxide reduction process in LK is hydrogen oxidation, followed by methane oxidation. In the MedS iron oxide reduction is most probably coupled to the oxidation of ammonium (Feammox) to molecular nitrogen (N_2_), and in a few cases may be coupled to methane oxidation. The Monte Carlo simulation agrees with the nominal model results for manganese reduction, and additionally predicts that iron reduction may be possible with some combinations of parameter values. These findings improve our understanding of the thermodynamic and kinetic controls on the composition of microbial communities and their effect on the geochemistry of methanic sediments.

## 1. Introduction

Iron and manganese oxide reduction are among the main microbial dissimilatory respiration processes in anaerobic sediments. During these processes, ferric iron (Fe(III)) or tetra-valent manganese (Mn(IV)) in amorphous or crystalline oxides are reduced to aqueous ferrous iron (Fe(II)) or divalent manganese (Mn(II)). As iron and manganese oxides are poorly soluble, while other electron acceptors (EA) are dissolved, their reduction reactions are relatively difficult to perform, and require unique direct and indirect electron transfer mechanisms. The known direct electron transfer mechanisms (DIET) for iron reduction require contact of the cell with the oxide and include (1) electron transfer via a protein on the cell’s surface, and (2) electron transfer by a conductive nanowire. The indirect electron transfer mechanisms do not require contact with the oxide and include (3) electron transfer via metal chelators (siderophores) that solubilize the oxides and reduce them on/in the cell, and (4) electron transfer by electron shuttling compounds (Dong et al., 2021; Gralnick and Newman, 2007).

The reactivity and bioavailability of the various oxides are dictated by different variables, for example, the particle size and surface area, their concentration and solubility. In general, amorphous oxides, such as amorphous iron oxyhydroxide (FeOOH), and poorly-crystalline oxides, such as ferrihydrite (Fe(OH)_3_), are considered more bioavailable than highly-crystalline oxides, such as hematite (Fe_2_O_3_) and magnetite (Fe_3_O_4_) (Bonneville et al., 2004; Larsen and Postma, 2001; Munch and Ottow, 1983). Manganese oxides, on the other hand, are mostly present in sediments as an amorphous solid (Burdige 1993). Amorphous manganese oxides (MnO_2_) were found to be more bioavailable to a pure culture than amorphous iron oxides (Lovley and Phillips, 1988a).

The coupled electron donors (ED) in the reduction of manganese and iron oxides are often organic compounds, which can also serve as a carbon source, including methane (CH_4_) or acetate (CH_3_COO^−^) (Jørgensen, 2000). However, inorganic compounds (for instance, hydrogen (H_2_), ammonium (NH_4_^+^) or sulfide (H_2_S)) can serve as electron donors as well, along with the utilization of carbon dioxide (CO_2_) as a carbon source (Jørgensen, 2000).

According to the “classical” theory of microbial respiration in sediments, the order of respiration reactions reflects a cascade of decreasing Gibbs free energies (Δ*G*_*r*_) of the net reduction-oxidation (redox) reactions (Jørgensen, 2000). In this model, as long as there are iron and manganese oxides, their thermodynamically more favorable reduction is expected to occur before (and at the expense of) sulfate reduction. For the same reason, sulfate reduction is expected to occur before methanogenesis, with the transition between these metabolisms occurring in a sulfate-methane transition zone (SMTZ). In the zone of iron and manganese reduction, the dissolved ferrous iron (Fe^2+^) tends to precipitate as siderite (FeCO_3_) (Ram et al., 1998; Sivan et al., 2011) and vivianite (Fe_3_(PO_4_)_2_) (März et al., 2008; Slomp et al., 2013; Hsu et al., 2014; Egger et al., 2015a), and the divalent manganese precipitates as rhodochrosite (MnCO_3_) (Kuleshov and Shterenberg, 1988). In the zone of sulfate reduction, the produced sulfide reacts with both dissolved Fe^2+^ and residual particulate Fe(III) minerals to form iron sulfide minerals, notably FeS and pyrite (FeS_2_) (Berner, 1984; Figueroa, 2023). The methanic zone is characterized by high dissolved methane concentrations and negligible sulfate and sulfide concentrations, as the excess of Fe(II) over dissolved sulfide leads to the precipitation of FeS and pyrite in both the sulfate reduction zone and the upper methanic zone (Conrad et al., 1986). In sediments underlying water columns with appreciable sulfate concentrations, such as marine sediments, anaerobic oxidation of methane (AOM), mainly coupled to sulfate reduction at the SMTZ, prevents the release to the water column and atmosphere of up to 90% of the methane produced in the methanic zone (Valentine, 2002). In freshwater environments, where sulfate concentrations are typically low, AOM is coupled to the reduction of other electron acceptors, such as nitrate, iron or manganese oxides (Crowe et al., 2011; Sivan et al., 2011; Bar-Or er al., 2017; Lenstra et al., 2023). It was found in enriched culture experiments that an archaea of the order *Methanosarcinales* can perform both Fe-AOM and Mn-AOM independently (Ettwig et al., 2016). In addition, methane can also be oxidized under hypoxia by aerobic bacteria (Ettwig et al., 2010; Bar-Or et al., 2015; 2017; Gafni et al., 2024), probably as part of a cryptic cycle of oxygen (Ettwig et al., 2010).

Iron and manganese oxides also play a critical role in the recycling of both organic and inorganic compounds through microbial processes. Specifically, anaerobic ammonium oxidation coupled to reduction of iron (Feammox) (Clement et al., 2005) or manganese (Mnammox) (Chen et al., 2020; Samperio-Ramos et al., 2024; Dai et al., 2024) is fundamental in the nitrogen and iron cycles. Both these reactions can result in the production of either molecular nitrogen, nitrite (NO_2_^−^) or nitrate (NO_3_^−^) (Yang et al., 2012; Chen et al., 2020). The production of nitrate and nitrite is particularly important, as it can participate in additional oxidation reactions, facilitating the formation of iron oxides (Sørensen and Thorling, 1991; Straub et al., 1996; Hauck et al., 2001; Kappler et al., 2005; Melton et al., 2014; Schaedler et al., 2018).

While the classical model has been a robust foundation for understanding microbial metabolism in natural aqueous environments, and many aspects of it still hold – specifically, the correct prediction of the cascade of respiration reactions – it does not account for the reduction of iron and manganese in the methanogenic zone. However, in many lacustrine and marine sediments, a significant increase in Fe^2+^ (Sivan et al., 2011; Riedinger et al., 2014; Treude et al., 2014; Egger et al., 2017) and dissolved divalent manganese (Mn^2+^) (Bar-Or et al., 2015; 2017) have been observed at the depth of methanogenesis, sometimes accompanied by a deep methane sink. A study on microbial cultures has revealed that microorganisms from the Black Sea are more effective at reducing manganese oxides compared to ferrihydrite (Thamdrup et al., 2000). Slow kinetics of microbial iron and manganese reduction and of iron oxide reaction with aqueous sulfide can partially explain how the oxides persist through the sulfidic zone, preserved below their “classical” reduction zone (Sivan et al., 2007; 2011), to result in another zone of iron and manganese reduction in the deep methanic zone. The presence of methane in this zone suggests that iron or manganese reduction may be coupled to AOM (Fe-AOM or Mn-AOM, respectively). Such iron-driven AOM was indeed evident in methanic lake sediments (e.g., Bar-Or et al., 2015, 2017; Sivan et al., 2011) and in nonsteady-state depositional marine system (Beal et al., 2009; Sivan et al., 2014; Egger et al., 2015b; Lenstra et al., 2023), but not in deep marine sediments in quasi-steady state (Aromokeye et al., 2020; Liang et al., 2022; Yorshansky et al., 2022). Thermodynamic and kinetic theory can be used to understand this difference between lacustrine and marine sediments, as is done here.

The local reaction rates of the various respiration processes determine their dominance in sediments, and these rates are not controlled by thermodynamics alone. Catabolic reaction rates (i.e., the rates at which cellular reactions break down molecules to release energy, typically measured per cell or tissue) and community sizes of different microbial functional groups dictate the total reaction rates, according to Eq. 1:

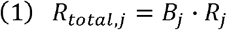

where *R*_*total,j*_ is the total reaction rate of the *j*th reaction (mol cm^−3^ day^−1^), *R*_*j*_ is the cell-specific catabolic rate (mol cell^−1^ day^−1^) and *B*_*j*_ is the number of cells of the functional group which performs the *j*th reaction in a given volume of water or sediment (cell cm^−3^).

Catabolism transforms the substrates into products in a net energy-releasing reaction. A large portion of the energy is stored in adenosine triphosphate (ATP), which is synthesized by the phosphorylation of adenosine diphosphate (ADP) as part of the electron transport chain, according to Eq. 2:

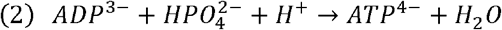

However, the synthesis of ATP itself requires energy, and the amount of energy available for this process affects the rate of catabolism, as reflected in the thermodynamic factor (see Eq. 8 in the Methods). The more energy-yielding the net catabolic reaction is, the more energy is available for the phosphorylation reaction, and the catabolism accelerates, until it is no longer limited by the thermodynamic driving force (Bethke et al., 2011; Jin, 2012; Jin and Bethke, 2003). The phosphorylation energy depends on the electron donor and acceptor, and for iron reduction it is approximately 45 kJ/mol (for iron reduction reactions coupled to H_2_ and CH_3_COO^−^ oxidation, the phosphorylation energies are 45 and 30-68 kJ mol ATP^−1^, respectively) (Jin, 2012).

The basic condition for the generation of one ATP molecule per one reaction turnover is |Δ*G*_*r*_| > Δ*G*_*p*_. However, it has been shown that it is possible to synthesize less than one ATP molecule per reaction turnover on average (i.e., more than one reaction turnover is required for the synthesis of a single ATP molecule), and the number of ATP molecules synthesized per one reaction turnover is referred to as the ATP yield (see Jin (2012) for a review of ATP yield values). The ATP yield is not fixed for any microbial functional group and can change according to environmental conditions. This means that in low-energy environments, such as most methanic sediments, the ATP yield of a certain microorganism will decrease, relatively to microorganisms of the same functional group in a high-energy environment (Bethke et al., 2011; Jin, 2012; Jin and Bethke, 2003). This allows the microorganism to increase the thermodynamic driving force and accelerate the catabolic process, even in energy-limiting environments. Thus, we would expect iron and manganese reducers in the methanogenic zone to generate less ATP per reaction turnover than the iron and manganese reducers in the “classical” iron and manganese reduction zones. Some of the ATP molecules generated during catabolism later provide energy for biomass synthesis during anabolism.

Although iron and manganese oxide respiration processes in deep lacustrine and marine sediments have been identified and discussed, the exact mechanisms, and the thermodynamic and kinetic controls of these processes are not well quantified. Recent research has provided insights into the rates of iron and manganese reduction when coupled to methane oxidation in the shallow northern Baltic Sea sediments (Lenstra et al., 2023). However, since iron and manganese reduction can be linked to multiple substrate oxidation reactions, and since functional genes for iron and manganese reduction and inhibitors for the iron and manganese cycles are poorly known, it is difficult to determine the rate of each specific reaction experimentally. Numerical modeling of the total iron reduction rate from pore-water Fe^2+^ concentration profiles is also problematic, as Fe^2+^ in anaerobic sediments is scavenged in various abiotic processes (Postma 1982; Egger et al. 2016), by complexation with organic compounds (Elderfield, 1981; Burdige, 1993), and potentially authigenic magnetite (Amiel et al., 2020).

Similarly, aqueous Mn^2+^ produced during manganese reduction can also be scavenged, mostly by complexation with organic compounds (Elderfield, 1981; Burdige, 1993). In anoxic sediments, where manganese and carbonate concentrations are high, some of the Mn^2+^ may also precipitate as a pure or mixed carbonate phase (Holdren et al., 1975; Grill, 1978; Suess, 1979; Aller 1980; Thomson et al., 1986; Middelburg et al., 1987; Burdige, 1993). Thus, quantification of both iron and manganese reduction in lacustrine and marine sediments remains only partly explored.

In the current study, we applied bioenergetic theory developed in previous studies (see Section 2) and quantified the potential dissimilatory iron and manganese reduction processes in diffusive controlled steady state methanic sediments. We considered the natural iron and manganese oxides found in both lake and marine sediments as electron acceptors and the existing electron donors. To model the controls on iron and manganese reduction rates in methanic sediments, we considered thermodynamic and kinetic parameters, besides the Gibbs free energy of the net catabolic reaction, which affect the total reaction rates. These factors were integrated into catabolic rate expressions, together with biomass growth rates and the expected community sizes of the different microbial functional groups that participate in these reactions.

## 2. Methods

### 2.1. Gibbs free energies of net redox reactions

The net standard-state Gibbs free energy (Δ*G*^*0*^_*r*_) of a reaction may be calculated as the difference between the Gibbs free energy of formation (Δ*G*^*0*^_*f*_) of the products and reactants participating in the reaction:

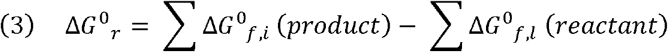

To calculate the Δ*G*^*0*^_*r*_ of a reaction at a different temperature and the same pressure, one may use the integral form of the Gibbs-Helmholtz equation:

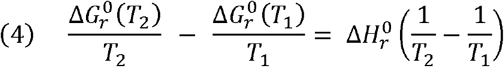

where T_i_ is the temperature, Δ*G*^*0*^_*r*_(T_i_) is the standard Gibbs free energy of the reaction at T=T_i_, and ΔH^0^_r_ is the enthalpy of the reaction, which is considered constant over small temperature ranges. The ΔH^0^_r_ of a reaction may be calculated using Hess’s law:

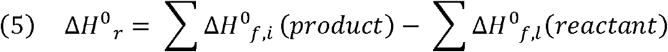

The Δ*G*^*0*^_*f*_ and Δ*H*^*0*^_*f*_ for 298 K and 1 atm of the relevant compounds were taken from multiple sources and are given in Table S1 in the supplementary data.

To determine the actual Gibbs free energy of a reaction (Δ*G*_*r*_), one must know the activities of the reactants and products.

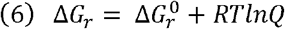

where *Q* is the reaction quotient, a function of the activities of the participating compounds and their stoichiometric coefficients. For example, for the reaction 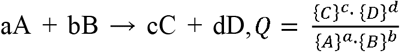 where capital letters in curly brackets denote the compounds’ activities and small letters are the compounds’ stochiometric coefficients.

The standard transformed Gibbs free energy of catabolism, 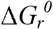^*’*^, under biochemical conditions of neutral pH, 298 K and 1 bar, is calculated using the equation:

where ν_H_^+^ is the stoichiometric coefficient of H^+^ in the reaction and *Kw* is the dissociation constant for water, which is equal to [H□O□]·[OH□] = 1.0·10□^14^ at 298 K. It should be noted that energetic calculations under these physical conditions are generally sufficient for modeling most metabolic processes (excluding thermophiles), with only minimal errors expected from small variations in these conditions (Amend and Shock, 2001).

### 2.2. Ca tabolic rate calculations

Microbially mediated reactions can be limited by the ability of transporters and enzymes to consume the substrate, leading to reaction rates that display a saturation behavior. This means that the rate of the reaction increases with increasing substrate concentration in the microbes’ growth environment, but only to a point at which the transporters and/or enzymes are saturated with respect to the substrate. Such saturation behavior is represented mathematically by Michaelis-Menten kinetics (Michaelis and Menten, 1913):

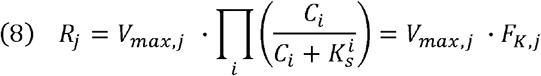

where *R*_*j*_ is the rate of the *j*th reaction, *V*_*max,j*_ is the maximum rate capacity of that reaction, and 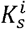 denotes the half-saturation constant of the *i*th species – the concentration at which the rate of the reaction is V_j_ = 0.5·V_max,j_. We denote the product of the Michaelis-Menten terms by the kinetic factor, *F*_*K, j*_.

Michaelis-Menten kinetics alone (Eq. 8) effectively predicts the maximum rates of biologically catalyzed reactions in high-energy environments. However, in energy-limited environments, such as methanic sediments, the substrate concentrations are usually low and the Δ*G*_*r*_ is a small negative number (i.e., the reaction is close to thermodynamic equilibrium). Catabolic rates are then even slower than is predicted from Michaelis-Menten kinetics (see Jin and Bethke (2007) for a review of studies substantiating this observation). To reproduce this behavior, a thermodynamic factor, *F*_*T*_, may be included in the rate expression:

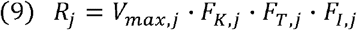

where *F*_*I,j*_ is the inhibition factor of certain inhibitor of the *j*th reaction. The kinetic, thermodynamic, and inhibition factors each range between zero and unity. A similar approach for calculating catabolic rates based on Michaelis-Menten kinetics was taken in previous biogeochemical models (Reed et al., 2014; Louca et al., 2016; Lenstra et al., 2023) LaRowe et al. (2012) suggested the following expression for the thermodynamic factor:

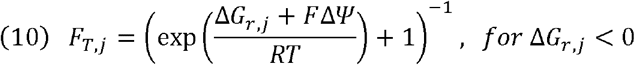

where *F* is the Faraday constant and ΔΨ is the electrochemical potential across the cell membrane, which must be maintained for the microorganism to exist. This potential may be too low to generate ATP in one reaction turnover, and in this case, ATP may be generated over several reaction turnovers. The electrochemical potential, ΔΨ, is often measured to be about 120-160 mV (Kadenbach, 2003; LaRowe et al., 2012). For Δ*G*_*r,j*_ of −23.51 kJ, *F*_*T,j*_ is equal to 0.99, which means that Δ*G*_*r,j*_ values more negative than this will have very little effect on the catabolic rate.

### 2.3. Biomass growth rate calculations

#### 2.3.1. Power supply and dissipated power

The approach adopted in this study to accomplish microbial growth rates is described by LaRowe and Amend (2015). This is done by comparing the power generated by catabolism to the dissipated power. The dissipated power is the power that is not invested in generation of new biomass or in the replacement of existing cellular components, but rather in a suite of other activities, including what is generally called maintenance (Pirt, 1965; Russell and Cook, 1995; Van Bodegom, 2007).

The power supply (*P*_*s,j*_) available to a certain microbial functional group from catalyzing the *j*th net redox reaction in a certain environment is given by:

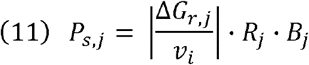

where *v*_*i*_ represents the stoichiometric coefficient of a particular reactant or product in the *j*th reaction. This parameter is used to normalize the Δ*G*_*r,j*_ to a specific reactant or product’s activity, so that the units of Δ*G*_*r,j*_ will be consistent with those of the rate (LaRowe and Amend, 2015). For Δ*G*_*r*_ in units of J mol^−1^ and *R*_*j*_ in mol cm^−3^ sec^−1^, *P*_*s*_ takes on units of volume-specific power, W cm^−3^.

Given a cell-specific maintenance power requirement (*P*_*cs*_), the catabolic power dissipated (*P*_*d,j*_) in a cm^3^ of sediment or fluid can by calculated by:

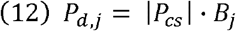

where *P*_*cs*_ for anaerobes is set between 3.6·10^−16^ and 8.6·10^−15^ W cell^−1^, based on pure culture experiments (Rinker and Kelly 2000).

According to this model, when *P*_*s,j*_ > *P*_*d,j*_, there is growth, or at least biomolecular replacement. Conversely, when *P*_*s,j*_ < *P*_*d,j*_, either the population catalyzing the *j*th net redox reaction shrinks, or the number of metabolically active cells decreases.

#### 2.3.2. Gibbs free energy of biomass synthesis

We define the energy invested in biomass synthesis of a functional group catalyzing the *j*th net redox reaction, Δ*G*^*0’*^ _*syn,j*_ (in kJ (mol e)^−1^ cell^−1^) considering biomass of the average composition C_5_H_7_O_2_N. The biomass synthesis energy may be calculated assuming that pyruvate (CH_3_COCOO^−^) is a representative cellular intermediate for biomass synthesis (external carbon source → pyruvate → biomass). Thus:

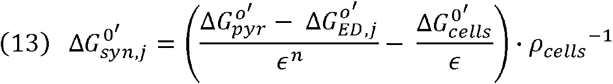

where Δ*G*^*0’*^_*pyr*_ and Δ*G*^*0’*^_*ED*,j_ are the standard Gibbs free energies of the half-reactions for reduction of pyruvate (35.09 kJ e-eq^−1^ at 298 K; Rittmann and McCarty, 2001) and oxidation of an electron donor (Table S2), respectively. The Δ*G*^*0’*^_*cells*_ is the Gibbs free energy required to synthesize biomass of the composition C_5_H_7_O_2_N from pyruvate at 298 K, which is equal to 18.8 kJ mol^−1^ per transfer of one electron. The efficiency of energy transfer, □, is equal to 0.6. The density of cells, ρ_cells_, is the number of cells per gram biomass. Assuming a cellular carbon content of 19 fg C cell^−1^, ρ_cells_ is equal to 2.794·10^13^ cells per gram biomass. The exponent *n* is an integer that equals to −1 when the conversion of the carbon source to pyruvate releases energy (for example, when using glucose as a carbon source), and equals +1 when it consumes energy (for example, when using acetate, methane or carbon dioxide as a carbon source).

Since an inorganic carbon source is used in autotrophic reactions, the electron donor does not produce pyruvate. Thus, for autotrophy, Rittmann and McCarty (2001) propose to set Δ*G*^*0’*^_*ED*_ equal to the value of the water-oxygen half-reaction in photosynthesis (−78.72 kJ e-eq^−1^ at 298 K), in which organic carbon is synthesized from carbon dioxide.

#### 2.3.3. The yield coefficient

The yield coefficient (*Y*_*j*_) describes the amount of biomass that can be produced from a given amount of energy generated in the respiration redox reaction. This value is not fixed, but rather a function of environmental variables.

We followed the methodology of Rittmann and McCarty (2001), who considered the fact that during microbial growth, electrons from the electron donor are proportioned into anabolism and catabolism. Given Δ*G*^*0’*^_*syn,j*_ per one moles of electrons (kJ (mol e^−^)^−1^), and Δ*G*^*0’*^ which is the standard Gibbs energy of (anabolic) cell synthesis (kJ (mol e^−^)^−1^), the energy and electron balances can be expressed as (VanBriesen, 2002):

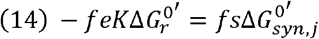

with

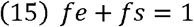

where *fe* is the fraction of electrons transferred from the electron donor to the terminal electron acceptor, and *fs* is the fraction of electrons invested in biomass production. The parameter *K* represents the efficiency of catabolic energy capture, which is assigned an average value of 0.6 (60%), while the remaining 40 percent of energy is dissipated as heat (Rittmann and McCarty, 2001; VanBriesen, 2002).

The yield coefficient (kJ cell^−1^) of each functional group catalyzing the *j*th reaction, which oxidizes a certain electron donor, is directly proportional to the values of 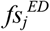 and inversely proportional to Δ*G*^*0’*^_*syn,j*_, as follows:

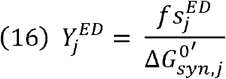

#### 2.3.4. The biomass yield

The approach described by LaRowe and Amend (2015) and adopted in this study for calculating biomass yields, combines the Δ*G*_*r*_ of biomolecule synthesis, catabolic power supply (*P*_*s,j*_) and the proportion of *P*_*s,j*_ that does not result in new biomass (*P*_*d,j*_). The change in the biomass of a certain functional group catalyzing the *j*th net redox reaction as a function of time, 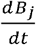, can be summarized as a combination of equations (13) and (14) and a yield coefficient, *Y*_*j*_ (cell J^−1^):

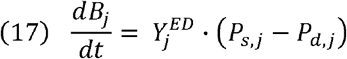

or in a more detailed form as:

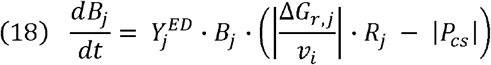

### 2.4. Monte Carlo parameter sampling and probability distributions

Monte Carlo experiments are computation algorithms that repeatedly select random samplings of uncertain parameter values. The uncertain parameters are selected from distributions (type and parameters, such as the mean and standard deviation for a normal distribution) that represent the state of knowledge and uncertainty in their values. Repeated calculation of the desired results for different uncertain parameter values within defined ranges provides an estimate of the distribution of the results, and their statistical properties. In this research, a Monte Carlo algorithm was used to obtain the distribution of catabolic rates and biomass growth rates of the selected microbial functional groups.

Types of distributions chosen for the Monte Carlo algorithm in this study are the normal and log-normal distributions. A log-normal distribution was selected in cases where the standard deviation in parameter values is exceptionally high, or to avoid sampling negative values of concentrations and kinetic parameters.

### 2.5. Flux calculations

To assess the availability of iron and sulfur species, fluxes of the relevant dissolved and solid compounds to the sediment were calculated. These calculations were not part of the bioenergetic model, but were used to evaluate the source of iron oxides in the methanic zone. Fluxes of dissolved chemical species (mol cm^−2^ sec^−1^) were calculated using Fick’s first law of diffusion:

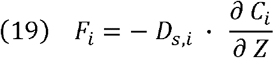

where *Z* is the depth within the sediment (cm), *C*_*i*_ is the concentration of dissolved species *i* at a certain depth (mol cm^−3^) and *D*_*S*_ is the diffusion coefficient of species *i* in pore water (cm^2^ sec^−1^). This coefficient may by calculated using the approximation D_s_ ~ D_0_ · □ ^2^ (after Lerman, 1979), where *D*_*0*_ is the diffusion coefficient of species *i* in water and □ is the sediment’s porosity, which is defined as the volume of pore water divided by the volume of the total sediment, assuming a saturated sediment.

Fluxes of solid species (mol cm^−2^ sec^−1^) were calculated for highly reactive iron oxides considering the sedimentation rate (ω, cm sec^−1^) and the concentration of a solid species *l*:

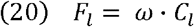

### 2.6. Study sites

Two study sites in this research are Lake Kinneret (LK) (Sea of Galilee) (Fig. S1), located in northern Israel, and the South Eastern Mediterranean Sea (MedS) (Fig. S2). In LK, a significant rise in the concentrations of dissolved Fe^2+^ and Mn^2+^ was observed in the methanic zone (up to 80 μM of Fe^2+^ and 82 μM of Mn^2+^; Fig. S3; Sivan et al., 2011; Bar-Or et al., 2015). This increase was accompanied by a decrease in methane concentrations (Fig. S3b), and previous studies have suggested the occurrence of Fe-AOM in this zone (Adler et al., 2011; Bar-Or et al., 2017; Sivan et al., 2011; Vigderovich et al., 2023). In this zone, Anaerobic Methanotrophs (ANME) type 1 were detected in low concentrations, as well as aerobic methanotrophs, iron reducing bacteria and possibly ammonium (NH_4_^+^) oxidizing bacteria (Bar-Or et al. 2015; Elul et al. 2021). The presence of possible ammonium oxidizing bacteria in the deep sediment of LK suggests that anaerobic ammonium oxidation can be coupled to reduction of iron (Feammox) or manganese (Mnammox) in this environment.

In the MedS, on the other hand, although a rise in Fe^2+^ concentrations was observed at the methanic zone (up to 64 μM; Fig. S4; Vigderovich et al., 2019), no Fe-AOM was detected (Yorshansky et al., 2022). The high Fe^2+^ concentrations and the presence of nitrite and nitrate suggest that Feammox and Mnammox may take place there (Vigderovich et al., 2019). Within the MedS methanic sediments, low concentrations of ANMEs type 1 and 2 were found as well as iron reducing bacteria and possibly anammox bacteria and magnetotactic bacteria (Vigderovich et al., 2019).

We used microbiological (Tables S3-S7) and geochemical (Tables S18-S20) data collected in previous studies from stations SG-1 and A at the MedS and LK sites, respectively. The data includes the porewater and the sedimentary geochemical (Figs. S3-S4) and microbial (Fig. S5) profiles. The abundant iron and manganese oxides in both sediments are: ferrihydrite (poorly crystalline Fe(III) hydroxide), hematite, goethite (α-FeOOH), lepidocrocite (γ-FeOOH), magnetite (Fe_3_O_4_), amorphous iron oxyhydroxide and amorphous manganese oxide. The abundant electron donors, which we included in the calculations, are: acetate, hydrogen, methane, ammonium and iron sulfide (FeS).

We assumed quasi-steady state in all sites based on the solid S and Fe/Mn oxide profiles, as well as porewater profiles from the Eastern Mediterranean (Sela-Adler et al., 2015; Wurgaft et al., 2019; Vigderovich et al., 2019; Amiel et al., 2020) and Lake Kinneret (Adler et al., 2011; Sivan et al., 2011; Bar-Or et al., 2017), along with our extensive profiling data. These profiles show natural variations due to factors such as drilling location changes and sampling but indicate steady conditions across the studied sites. This assumption was used only in the calculation of total iron to sulfate fluxes, supporting the persistence of iron oxides throughout the sulfidic zone. However, it was not part of the bioenergetic model development, and we have clarified this in Section 2.6 of the revised manuscript.

Contrasting a lacustrine environment (LK) with a marine environment (MedS) provides valuable insights into the similarities and differences in biogeochemical processes across distinct aquatic systems. In the MedS, we specifically chose two study sites: one with relatively high methane concentrations (SG-1) and another with lower methane concentrations (PC-3). This comparison allowed us to assess how methane influences microbial processes under different conditions.

## 3. Re sults

### 3.1. Gibbs free energy of net redox reactions

The standard Gibbs free energy and enthalpy (Δ*G*^*0*^_*r*_ and Δ*H*^*0*^_*r*_, respectively) of all net redox reactions (including magnetite reduction, for which no kinetic data were available), were calculated after normalization to the transfer of one electron (see balanced reactions, presented in Tables S9-S15; Eq. 3). Based on the activities of relevant aqueous species in the methanic zones of the MedS and LK (Tables S18-S20), the values of the actual Gibbs free energy were calculated for the methanic zone of both sites (Table S16; Fig. 1). The N_2_ concentrations were not measured, and we assumed that N_2_ was saturated (i.e., at equilibrium with atmospheric N_2_) at the sites’ temperature of 14°C. The resulting aqueous N_2_ concentration is 0.6 mM.

**Figure 1:**
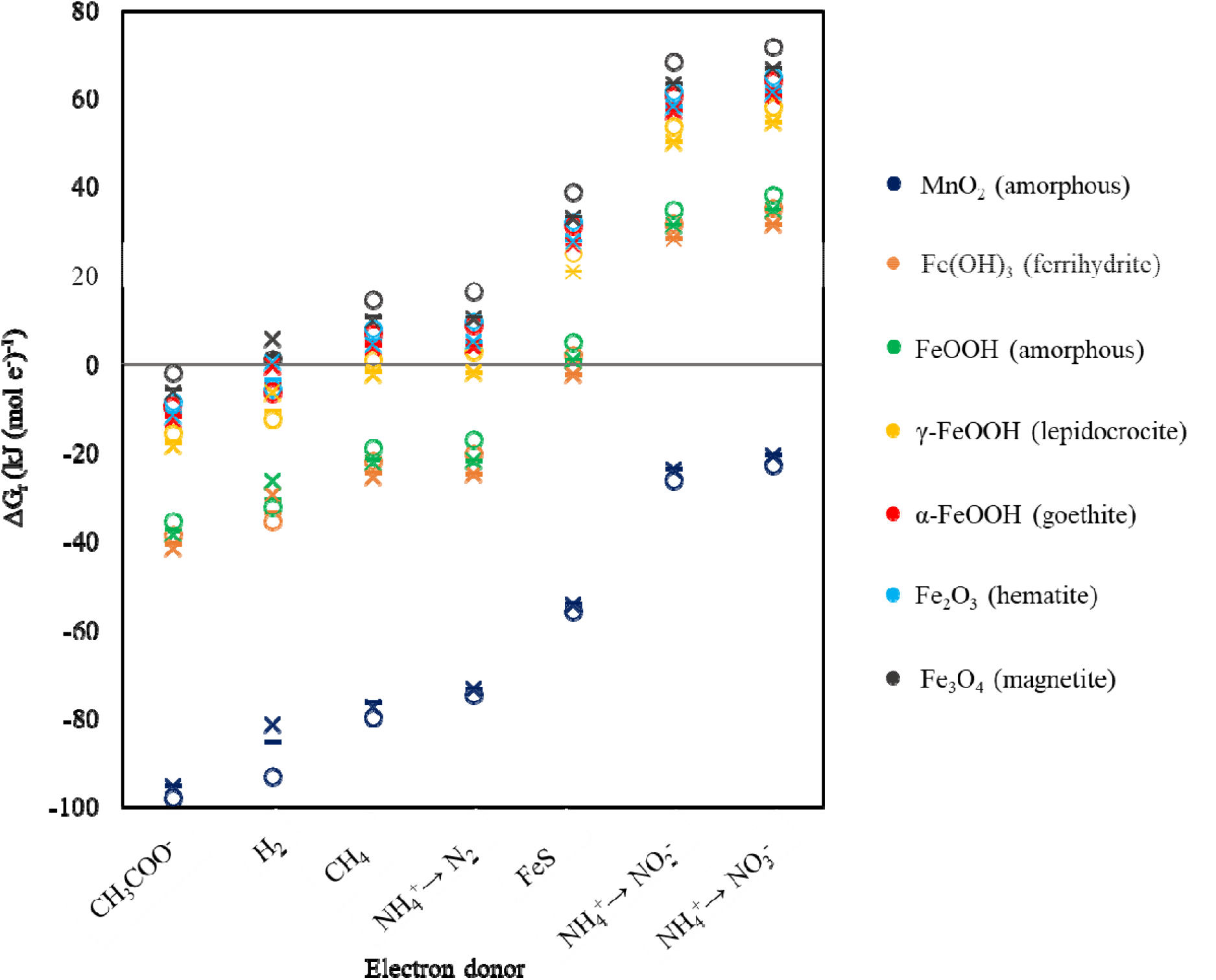
Actual Gibbs free energy of the net redox reactions. The symbol ○ denotes Lake Kinneret, wh ile X and – denote the marine sites SG-1 and PC-3, respectively.

**Figure 2:**
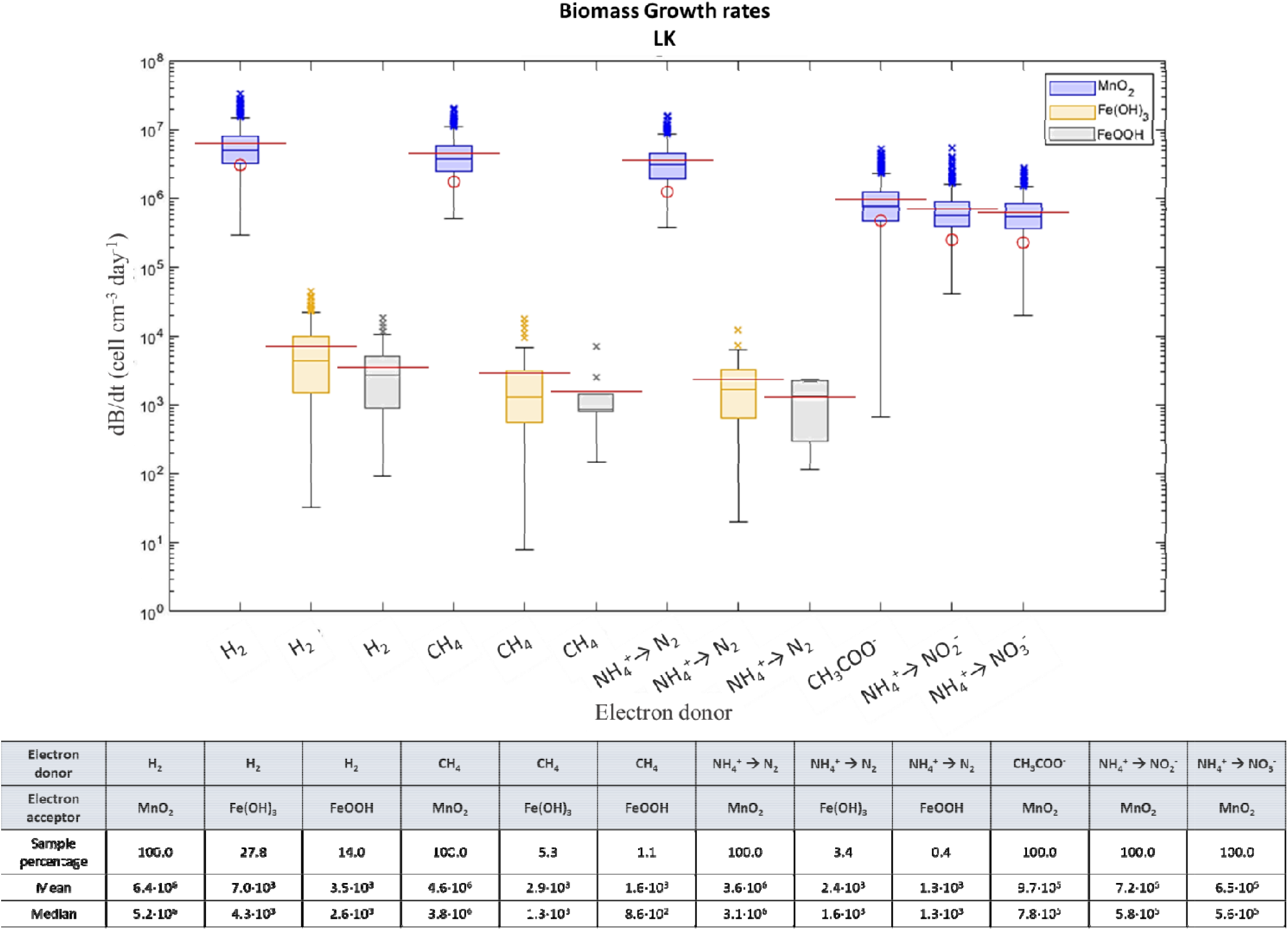
Monte Carlo simulation of biomass growth rates in LK. Whiskers cover 99.3% of all par ameter samplings, and the outliers (marked by ‘×’) represent the remaining 0.7% of the parameter sam plings. The boxes present the 25^th^ to the 75^th^ percentiles, and the bold lines in the box centers rep resent the median. Red lines represent the mean values, and red circles are the values calculated wit h the nominal model. The sample percentage in the table represents the proportion of times each out come occurred out of 1000 runs of the Monte Carlo algorithm.

According to the Δ*G*_*r*_ calculations, both sites are similar in terms of the order of favorability of redox reactions with the various electron donors and acceptors considered (Fig. 1). Acetate is the most energetically favorable electron donor, followed by H_2_, methane, ammonium and FeS. Ammonium oxidation to N_2_ generates more energy than ammonium oxidation to nitrite or nitrate, which are the least energetically favorable oxidation half-reactions.

Amorphous manganese oxide is the most favorable electron acceptor in all cases, followed by amorphous and poorly crystalline iron oxides (ferrihydrite), and lastly the crystalline iron oxides, with substrate oxidation by magnetite and hematite reduction producing near-equilibrium or positive Δ*G*_*r*_ values in all cases.

### 3.2. Catabolic rates in the nominal model

Nominal values (i.e., fixed, predetermined values with no associated uncertainty or error bars) of catabolic rates were calculated for all modeled reactions in the MedS and LK, except FeS oxidation, for which kinetic parameters are missing (see Table S17 for kinetic parameters; Tables 1-3; Figs. 3, 5 and 7). The nominal model (i.e., a model which considers nominal values) does not present possible variations in catabolic rates associated with uncertainty in parameter values, considering instead fixed mean values of kinetic parameters, fixed high concentrations of reactants and low concertation of products measured at the selected stations (Tables S18-S20). The sensitivity of the model results to all parameter values is examined in Section 3.4. The inhibition factor (*F*_*I,j*_) in all calculations was neglected, as the concentrations of O_2_ are negligible in the methanic zones, and manganese oxides and nitrate do not inhibit dissimilatory iron reduction, but rather oxidize the product Fe^2+^ in a secondary reaction (Lovley and Phillips, 1988b; Obuekwe et al., 1981).

**Table 1:** Nominal catabolic rates (R) (mol oxide cell^−1^ day^−1^) of biomass generating reactions when consuming different electron donors and electron acceptors (EA) in LK.

| EA | $\text{CH}_4$ | $\text{H}_2$ | $\text{NH}_4^+ \rightarrow \text{N}_2$ | $\text{NH}_4^+ \rightarrow \text{NO}_2^-$ | $\text{NH}_4^+ \rightarrow \text{NO}_3^-$ | $\text{CH}_3\text{COO}^-$ |
| --- | --- | --- | --- | --- | --- | --- |
| $\text{MnO}_2$ (amorphous) | $8.0 \cdot 10^{-14}$ | $6.9 \cdot 10^{-14}$ | $7.2 \cdot 10^{-14}$ | $7.2 \cdot 10^{-14}$ | $7.0 \cdot 10^{-14}$ | $6.2 \cdot 10^{-15}$ |
| $\text{Fe}(\text{OH})_3$ (ferrihydrite) | $1.2 \cdot 10^{-15}$ | $1.1 \cdot 10^{-15}$ | | | | |
| $\text{FeOOH}$ (amorphous) | | $8.5 \cdot 10^{-16}$ | | | | |

**Table 2:**
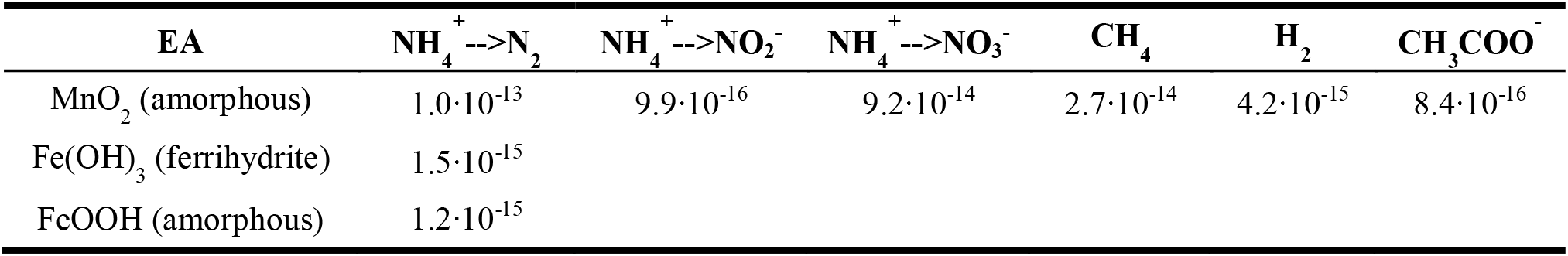
Nominal catabolic rates (R) (mol oxide cell^−1^ day^−1^) of biomass generating reactions when consuming different electron donors and electron acceptors (EA) in PC-3.

| EA | $\text{NH}_4^+ \rightarrow \text{N}_2$ | $\text{NH}_4^+ \rightarrow \text{NO}_2^-$ | $\text{NH}_4^+ \rightarrow \text{NO}_3^-$ | $\text{CH}_4$ | $\text{H}_2$ | $\text{CH}_3\text{COO}^-$ |
| --- | --- | --- | --- | --- | --- | --- |
| $\text{MnO}_2$ (amorphous) | $1.0 \cdot 10^{-13}$ | $9.9 \cdot 10^{-16}$ | $9.2 \cdot 10^{-14}$ | $2.7 \cdot 10^{-14}$ | $4.2 \cdot 10^{-15}$ | $8.4 \cdot 10^{-16}$ |
| $\text{Fe}(\text{OH})_3$ (ferrihydrite) | $1.5 \cdot 10^{-15}$ | | | | | |
| $\text{FeOOH}$ (amorphous) | $1.2 \cdot 10^{-15}$ | | | | | |

**Table 3:** Nominal catabolic rates (R) (mol oxide cell^−1^ day^−1^) of biomass generating reactions when consuming different electron donors and electron acceptors (EA) in SG-1.

| EA | $\text{NH}_4^+ \rightarrow \text{N}_2$ | $\text{NH}_4^+ \rightarrow \text{NO}_2^-$ | $\text{NH}_4^+ \rightarrow \text{NO}_3^-$ | $\text{CH}_4$ | $\text{CH}_3\text{COO}^-$ | $\text{H}_2$ |
| --- | --- | --- | --- | --- | --- | --- |
| $\text{MnO}_2$ (amorphous) | $1.0 \cdot 10^{-13}$ | $9.9 \cdot 10^{-14}$ | $9.2 \cdot 10^{-14}$ | $6.3 \cdot 10^{-14}$ | $8.4 \cdot 10^{-16}$ | $2.2 \cdot 10^{-16}$ |
| $\text{Fe}(\text{OH})_3$ (ferrihydrite) | $1.5 \cdot 10^{-15}$ | | | | | |
| $\text{FeOOH}$ (amorphous) | $1.2 \cdot 10^{-15}$ | | | | | |

**Figure 3:**
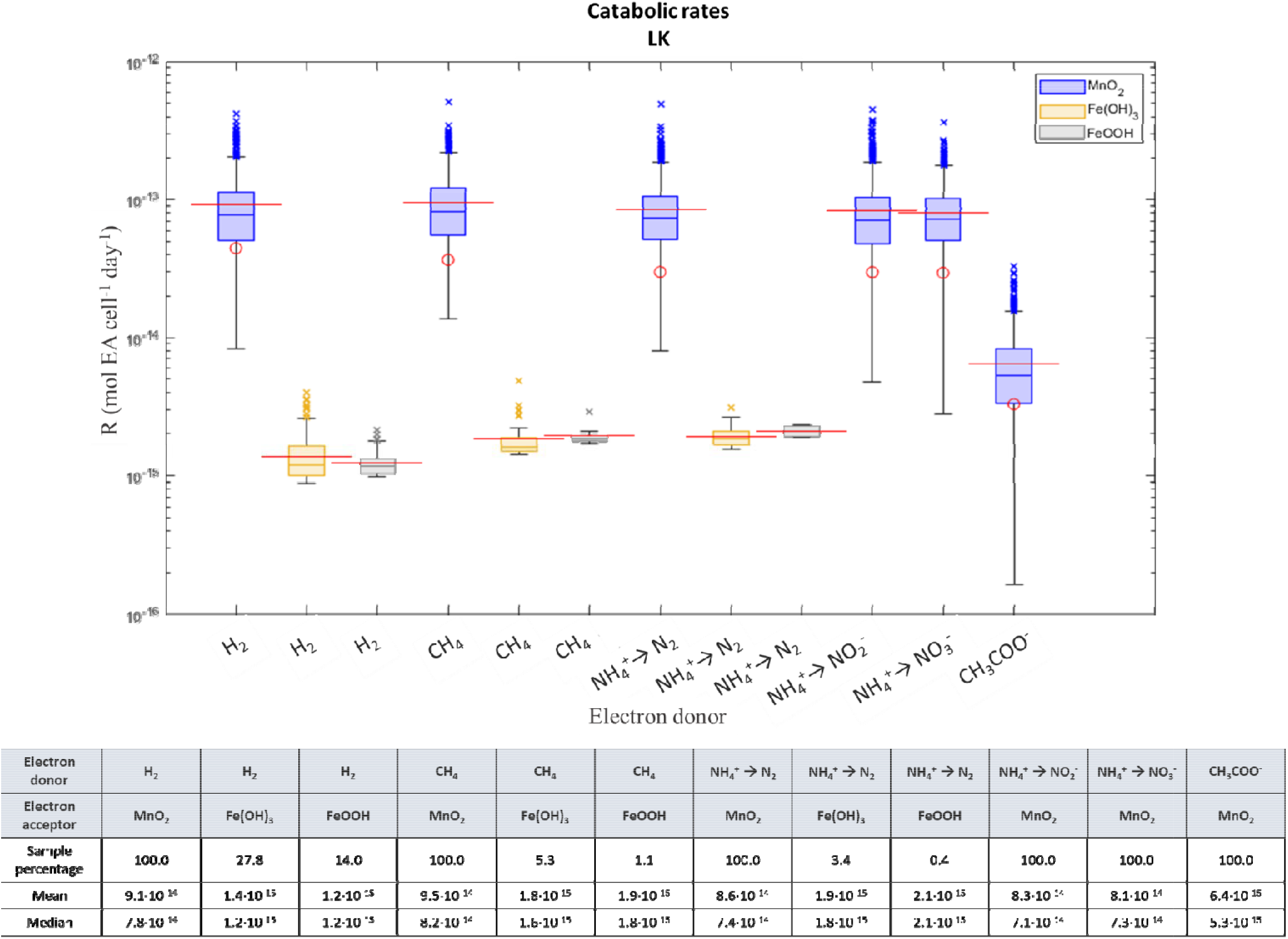
Monte Carlo simulation of catabolic rates in LK. Whiskers cover 99.3% of all parameter sam plings, and the outliers (marked by ‘×’) represent the remaining 0.7% of the parameter samplings. Th e boxes present the 25^th^ to the 75^th^ percentiles, and the bold lines in the box centers represent the me dian. Red lines represent the mean values, and red circles are the values calculated with the nom inal model. The sample percentage in the table represents the proportion of times each outcome occ urred out of 1000 runs of the Monte Carlo algorithm.

**Figure 4:**
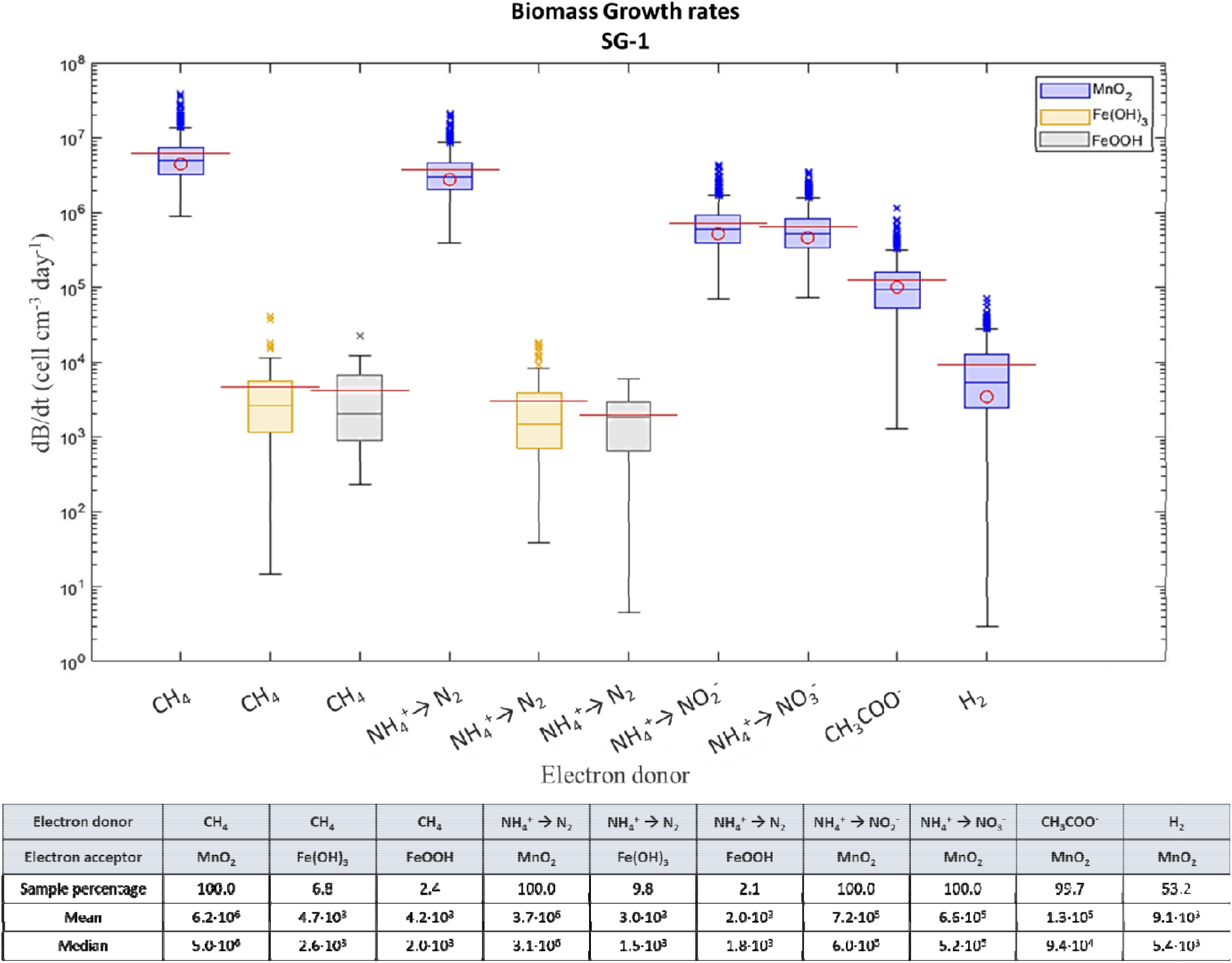
Monte Carlo simulation of biomass growth rates in SG-1. Whiskers cover 99.3% of all par ameter samplings, and the outliers (marked by ‘×’) represent the remaining 0.7% of the parameter sam plings. The boxes present the 25^th^ to the 75^th^ percentiles, and the bold lines in the box centers rep resent the median. Red lines represent the mean values, and red circles are the values calculated wit h the nominal model. The sample percentage in the table represents the proportion of times each out come occurred out of 1000 runs of the Monte Carlo algorithm.

**Figure 5:**
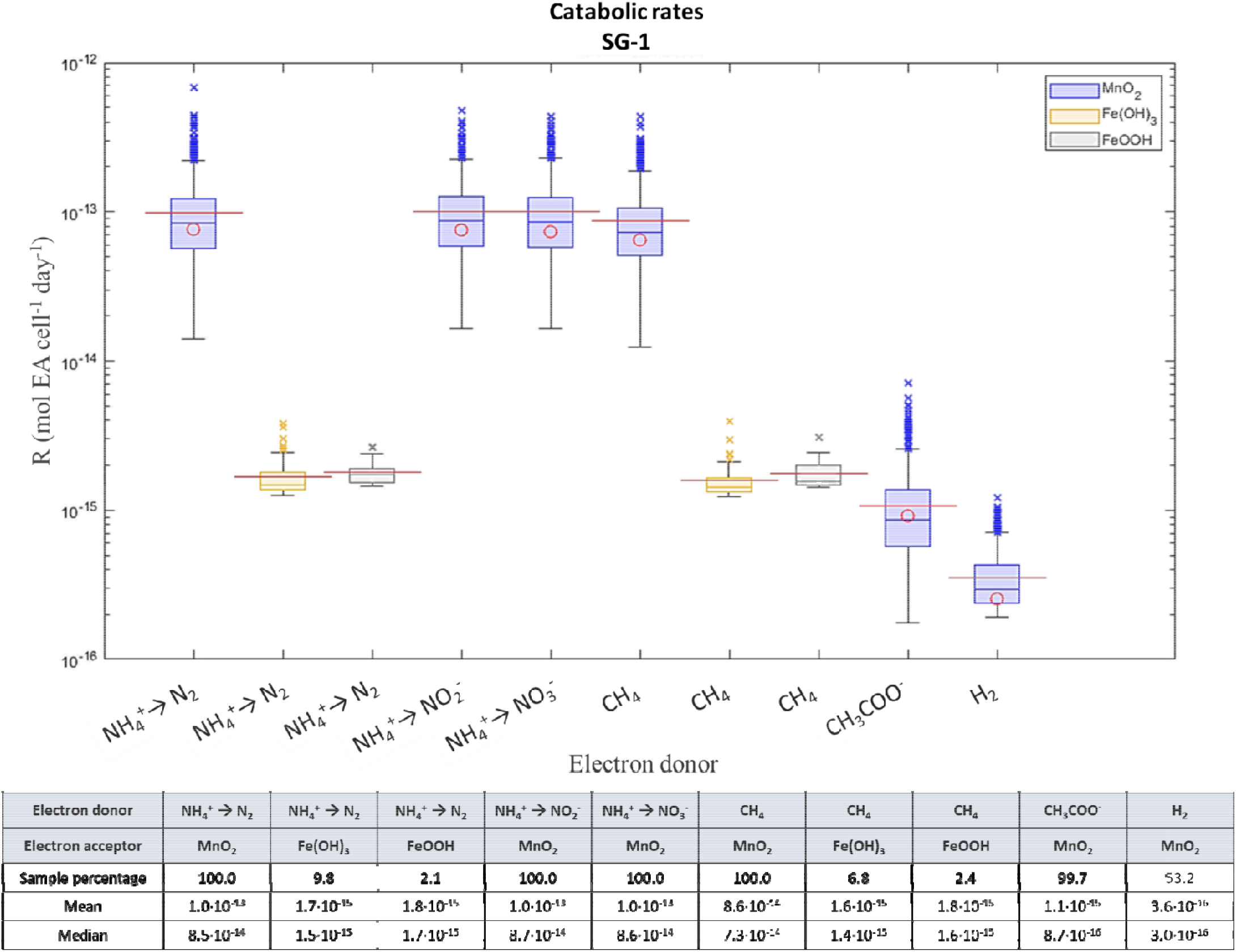
Monte Carlo simulation of catabolic rates in SG-1. Whiskers cover 99.3% of all parameter sam plings, and the outliers (marked by ‘×’) represent the remaining 0.7% of the parameter samplings. Th e boxes present the 25^th^ to the 75^th^ percentiles, and the bold lines in the box centers represent the me dian. Red lines represent the mean values, and red circles are the values calculated with the nominal model. The sample percentage in the table represents the proportion of times each outcome occ urred out of 1000 runs of the Monte Carlo algorithm.

**Figure 6:**
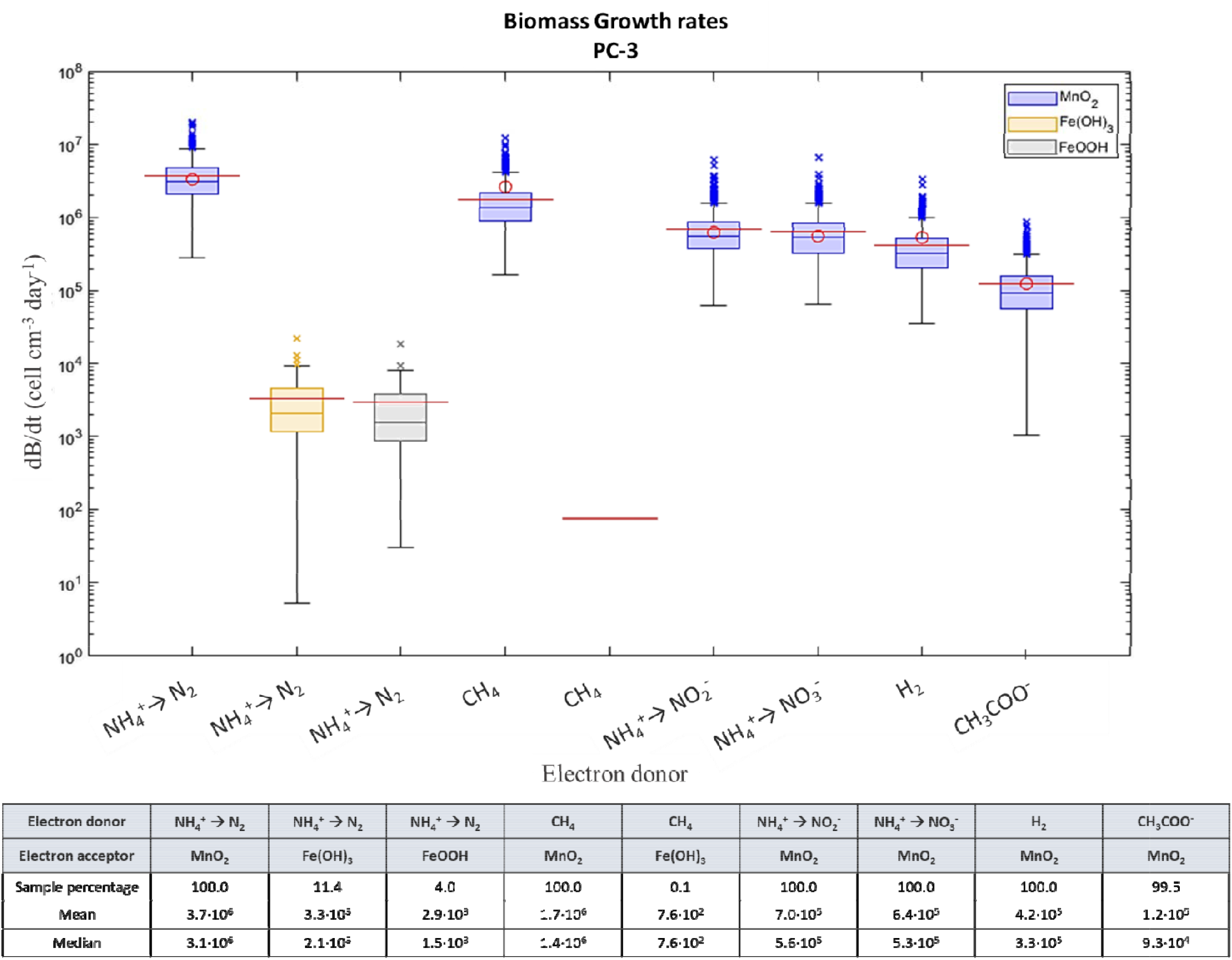
Monte Carlo simulation of biomass growth rates in PC-3. Whiskers cover 99.3% of all par ameter samplings, and the outliers (marked by ‘× ‘) represent the remaining 0.7% of the parameter sam plings. The boxes present the 25^th^ to the 75^th^ percentiles, and the bold lines in the box centers rep resent the median. Red lines represent the mean values, and red circles are the values calculated wit h the nominal model. The sample percentage in the table represents the proportion of times each out come occurred out of 1000 runs of the Monte Carlo algorithm.

**Figure 7:**
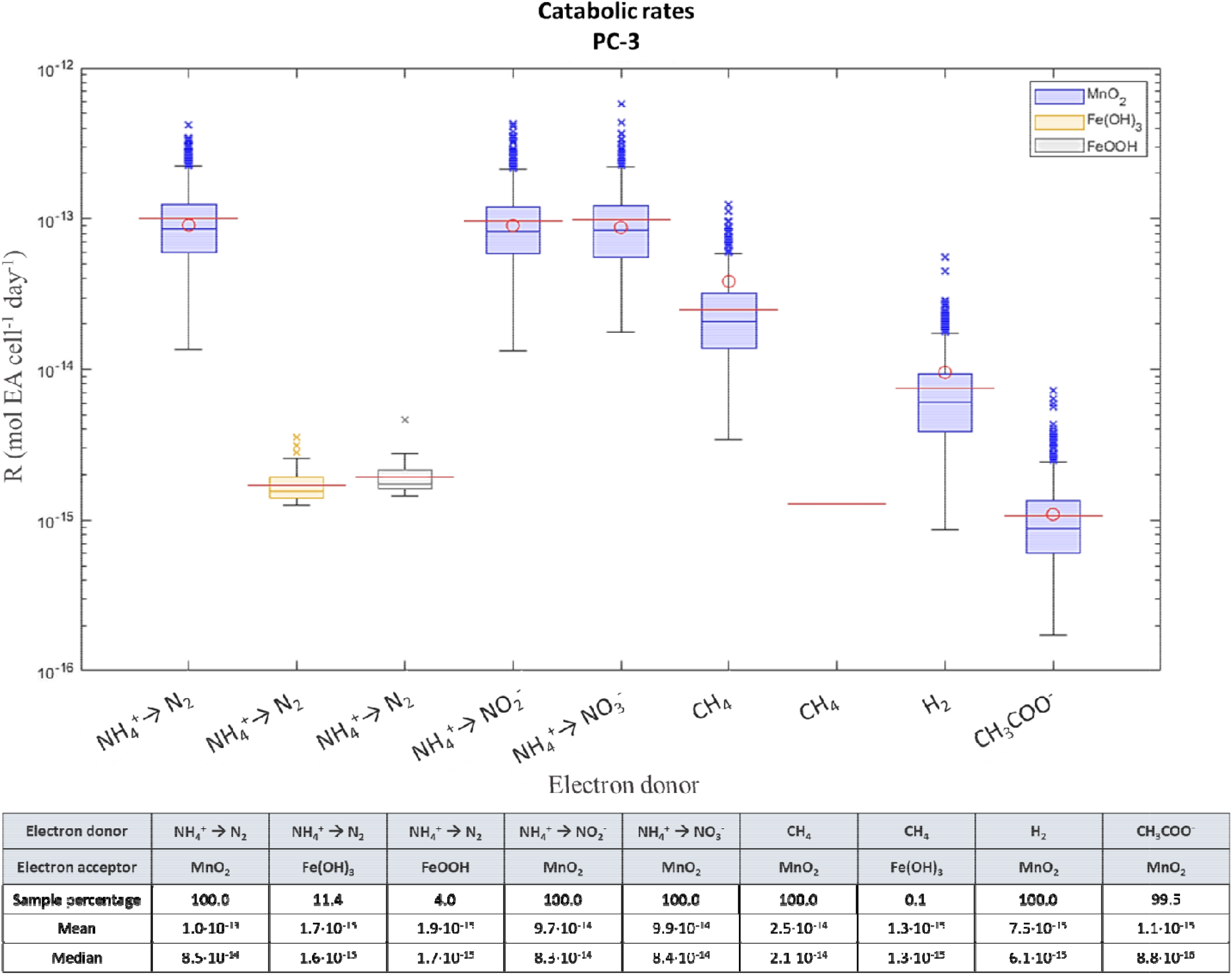
Monte Carlo simulation of catabolic rates in PC-3. Whiskers cover 99.3% of all parameter sam plings, and the outliers (marked by ‘× ‘) represent the remaining 0.7% of the parameter samplings. Th e boxes present the 25^th^ to the 75^th^ percentiles, and the bold lines in the box centers represent the me dian. Red lines represent the mean values, and red circles are the values calculated with the nom inal model. The sample percentage in the table represents the proportion of times each outcome occ urred out of 1000 runs of the Monte Carlo algorithm.

We present here only the catabolic rates of biomass generating metabolisms (see Section 3.3). In the nominal model, only manganese reduction with a variety of electron donors is energetic enough to generate biomass. The highest catabolic rates calculated for LK are for the oxidation of hydrogen, methane and ammonium, with acetate oxidation generating the lowest catabolic rates. In the MedS (PC-3 and SG-1), the ammonium oxidation rate is highest, followed by methane oxidation. This is followed by hydrogen and acetate oxidation in PC-3, whereas in SG-1 the catabolic rates calculated for acetate oxidation are higher than for hydrogen oxidation.

### 3.3. Biomass growth rates in the nominal model

Cell counts of archaea and bacteria in the methanic zone of SG-1 were obtained by qPCR analysis (Vigderovich et al., 2019) (Table S3). The cell counts were used in the biomass growth rate calculations in both the MedS and LK, in addition to archaea-bacteria ratios in LK, obtained from RNA sequencing (Table S4). The nominal model considers fixed cell counts, along with fixed probable values of kinetic parameters, parameters for biomass synthesis and measured concentrations. Thus, the nominal model does not show possible variations in the biomass growth rates, but rather the biomass growth rates in a single probable case (see Section 3.2).

Since ATP yields in low-energy environments are relatively low (see Section 1.1), it is reasonable to assume that *P*_*cs*_ values in such environments would be low. Thus, as this study focuses on anaerobic respiration in extremely low-energy environments, biomass growth rates for all modeled reactions (except iron sulfide oxidation) were calculated considering a low *P*_*cs*_ value of 3.6·10^−16^ W cell^−1^ (see Section 2.2.1; Tables 4-6; Figs. 2, 4 and 6). We assumed that only archaea can anaerobically oxidize methane (Orphan et al., 2002; Niemann et al., 2006; Milucka et al., 2012) and assigned the number of cells of methanotrophic archaea to equal the total number of archaeal cells (Tables S3-S4), with different dissipated power (*P*_*d*_) values (Eq. 12) resulting for methanogens and all other functional groups (Table S5). At both sites, Feammox (ammonium oxidation with Fe(III) oxyhydroxides) enabled biomass growth only with molecular nitrogen as the oxidation product (and not nitrite or nitrate), while all ammonium oxidation half-reactions produced biomass when coupled to manganese reduction.

**Table 4:** Nominal biomass growth rates (dB/dt) (cell cm^−3^ day^−1^) when consuming different electron donors (ED) and electron acceptors (EA) in LK. *P*_*cs*_ = 3.6·10^−16^ W/cell. Negative calculated growth rates are marked in red.

| EA | H<sub>2</sub> | CH<sub>4</sub> | NH<sub>4</sub><sup>+</sup> → N<sub>2</sub> | CH<sub>3</sub>COO<sup>-</sup> | NH<sub>4</sub><sup>+</sup> → NO<sub>2</sub><sup>-</sup> | NH<sub>4</sub><sup>+</sup> → NO<sub>3</sub><sup>-</sup> |
| --- | --- | --- | --- | --- | --- | --- |
| MnO<sub>2</sub> (amorphous) | 4.7·10<sup>6</sup> | 3.7·10<sup>6</sup> | 3.0·10<sup>6</sup> | 9.3·10<sup>5</sup> | 5.7·10<sup>5</sup> | 5.1·10<sup>5</sup> |
| Fe(OH)<sub>3</sub> (ferrihydrite) | 4.6·10<sup>3</sup> | 1.1·10<sup>3</sup> | -8.3·10<sup>2</sup> | -2.3·10<sup>4</sup> |  |  |
| FeOOH (amorphous) | 2.2·10<sup>2</sup> | -6.8·10<sup>3</sup> | -3.5·10<sup>3</sup> | -2.3·10<sup>4</sup> |  |  |
| γ-FeOOH (lepidocrocite) | -9.8·10<sup>3</sup> | -8.7·10<sup>3</sup> | -8.5·10<sup>3</sup> | -2.3·10<sup>4</sup> |  |  |
| α-FeOOH (goethite) | -1.1·10<sup>4</sup> |  |  | -2.3·10<sup>4</sup> |  |  |
| Fe<sub>2</sub>O<sub>3</sub> (hematite) | -1.0·10<sup>4</sup> |  |  | -2.8·10<sup>4</sup> |  |  |

**Table 5:** Nominal biomass growth rates (dB/dt) (cell cm^−3^ day^−1^) when consuming different electron donors (ED) and electron acceptors (EA) in PC-3. *P*_*cs*_ = 3.6·10^−16^ W/cell. Negative calculated growth rates are marked in red.

| EA | NH<sub>4</sub><sup>+</sup> → N<sub>2</sub> | CH<sub>4</sub> | NH<sub>4</sub><sup>+</sup> → NO<sub>2</sub><sup>-</sup> | NH<sub>4</sub><sup>+</sup> → NO<sub>3</sub><sup>-</sup> | H<sub>2</sub> | CH<sub>3</sub>COO<sup>-</sup> |
| --- | --- | --- | --- | --- | --- | --- |
| MnO<sub>2</sub> (amorphous) | 3.6·10<sup>6</sup> | 1.7·10<sup>6</sup> | 6.2·10<sup>5</sup> | 5.2·10<sup>5</sup> | 2.2·10<sup>5</sup> | 8.7·10<sup>4</sup> |
| Fe(OH)<sub>3</sub> (ferrihydrite) | 3.8·10<sup>3</sup> | -9.0·10<sup>3</sup> |  |  | -1.0·10<sup>4</sup> | -2.3·10<sup>4</sup> |
| FeOOH (amorphous) | 2.4·10<sup>2</sup> | -1.0·10<sup>4</sup> |  |  | -1.0·10<sup>4</sup> | -2.3·10<sup>4</sup> |
| γ-FeOOH (lepidocrocite) | -7.5·10<sup>3</sup> | -1.3·10<sup>4</sup> |  |  | -9.7·10<sup>3</sup> | -2.1·10<sup>4</sup> |
| α-FeOOH (goethite) |  |  |  |  | -9.7·10<sup>3</sup> | -2.1·10<sup>4</sup> |
| Fe<sub>2</sub>O<sub>3</sub> (hematite) |  |  |  |  | -1.2·10<sup>4</sup> | -2.0·10<sup>4</sup> |

**Table 6:** Nominal biomass growth rates (dB/dt) (cell cm^−3^ day^−1^) when consuming different electron donors (ED) and electron acceptors (EA) in SG-1. *P*_*cs*_ = 3.6·10^−16^ W/cell. Negative calculated growth rates are marked in red.

| EA | NH<sub>4</sub><sup>+</sup> → N<sub>2</sub> | CH<sub>4</sub> | NH<sub>4</sub><sup>+</sup> → NO<sub>2</sub><sup>-</sup> | NH<sub>4</sub><sup>+</sup> → NO<sub>3</sub><sup>-</sup> | CH<sub>3</sub>COO<sup>-</sup> | H<sub>2</sub> |
| --- | --- | --- | --- | --- | --- | --- |
| MnO<sub>2</sub> (amorphous) | 3.6·10<sup>6</sup> | 4.1·10<sup>6</sup> | 6.2·10<sup>5</sup> | 5.2·10<sup>5</sup> | 8.7·10<sup>4</sup> | 1.0·10<sup>3</sup> |
| Fe(OH)<sub>3</sub> (ferrihydrite) | 3.8·10<sup>3</sup> | -1.4·10<sup>3</sup> |  |  | -2.3·10<sup>4</sup> | -1.1·10<sup>4</sup> |
| FeOOH (amorphous) | 2.4·10<sup>2</sup> | -5.0·10<sup>3</sup> |  |  | -2.3·10<sup>4</sup> | -1.1·10<sup>4</sup> |
| γ-FeOOH (lepidocrocite) | -7.5·10<sup>3</sup> | -1.3·10<sup>4</sup> |  |  | -2.1·10<sup>4</sup> | -1.0·10<sup>4</sup> |
| α-FeOOH (goethite) |  |  |  |  | -2.0·10<sup>4</sup> | -9.7·10<sup>3</sup> |
| Fe<sub>2</sub>O<sub>3</sub> (hematite) |  |  |  |  | -2.5·10<sup>4</sup> | -9.7·10<sup>3</sup> |

In LK, hydrogen oxidation is the most biomass-generating oxidation half-reaction, followed by methane oxidation, ammonium oxidation to nitrogen, then acetate oxidation, and lastly ammonium oxidation to nitrite and nitrate. In the Mediterranean sediments at station PC-3, ammonium oxidation to molecular nitrogen is the most biomass-generating oxidation half-reaction followed by methane oxidation, ammonium oxidation to nitrite and nitrate, and lastly the oxidation of hydrogen and acetate. In SG-1, methane oxidation is the most biomass generating, then ammonium oxidation to molecular nitrogen, nitrite and nitrate, followed by the oxidation of acetate and hydrogen. The calculated rates of all iron reduction half-reactions in the nominal model are negative, which indicates insufficient energy for net forward reaction and growth in this scenario.

### 3.4. Sensitivity analysis

The model’s sensitivity to uncertain thermodynamic and kinetic parameters was tested (Tables S21-S33; Figs. S8-S14). In this sensitivity analysis, biomass growth rates for each modeled bioreaction were calculated for different parameter values. The parameters to which the model sensitivity was tested include all relevant electron donor and electron acceptor concentrations and half-saturation constants (*K*_*S*_), maximum rate capacities (*V*_*max*_), cell-specific maintenance power (*P*_*cs*_), the efficiency of energy transfer (□), the efficiency of catabolic energy capture (*K*) and the Gibbs free energy required to synthesize biomass(Δ*G*^*0’*^_*cells*_).

We find that the model is most sensitive to *V*_*max*_ and □, and shows some sensitivity to *K*_*S*_ values of both electron donors and electron acceptors, to Δ*G*^*0’*^_*cells*_, and to *K*. The results are insensitive to *P*_*cs*_. In addition, the biomass growth rates are directly proportional to the cell counts (*B*) (Eq. 18), and ρ_*cells*_ (Eqs. 13, 16 and 18), which are also uncertain model parameters. Nonetheless, since all mentioned parameters, other than *V*_*max*_ and *K*_*S*_ values, are the same for all modeled bioreactions (though with different cell counts for methane oxidation), these parameters do not affect the model trends in catabolic and growth rates.

Overall, the model demonstrates high sensitivity to reactants present in low concentrations and lower sensitivity to those present in high concentrations (Figs S8-S13; see relevant concentrations in Tables S18-S20). In most cases, the model is sensitive to electron donor concentrations, and in some cases, also to electron acceptor concentrations. For example, hydrogen concentrations in the MedS (SG-1 and PC-3), methane concentrations in PC-3, and acetate concentrations at all stations are relatively low, so in these instances the model is more sensitive to electron donor concentrations than to electron acceptor concentrations. On the other hand, hydrogen concentrations in LK, methane concentrations in LK and SG-1, and ammonium concentrations at all stations are relatively high, so in these instances the model is sensitive to the concentrations of both the electron donor and the electron acceptor.

### 3.5. The range of catabolic and biomass growth rates given parameter uncertainty

Given the inherent variability and uncertainty in the values of key parameters (such as kinetic constants, ΔGr, and biomass yield), as well as the limited statistical data available for these parameters, Monte Carlo simulations were employed to estimate the range of catabolic and biomass growth rates for iron and manganese reduction at all sites. The Monte Carlo algorithm was coded in MATLAB^®^. At each site, the algorithm calculated the catabolic and biomass growth rates for 1000 combinations of parameters randomly drawn from distributions that represent the uncertainty in their values (Figs. 2-7). We have compiled a statistical analysis of measurements of dissolved and solid chemical species in our study sites, and these distributions were incorporated into the Monte Carlo simulation (Tables S18-S20). All uncertain parameters (listed in Section 3.4) were randomly sampled in the algorithm (see Table S17 for mean kinetic parameters; see Table S34 for distribution and standard deviations of model parameters). A log-normal distribution was selected in cases where the standard deviation in parameter values is exceptionally high, or to avoid sampling negative values of concentrations and kinetic parameters.

In all cases, the mean model result (catabolic rate, biomass growth rate) is slightly higher than its median value. In most instances, the result obtained with the nominal model is close to both the mean and median results, with the exception of biomass growth rates and catabolic rates in LK, where the results for manganese reduction are widely dispersed.

While the nominal model predicts only manganese reduction, the Monte Carlo simulation predicts both manganese reduction and iron reduction (marginally) at all stations, with ferrihydrite and amorphous iron oxyhydroxide being the only iron oxides that generate positive biomass growth rates with some combinations of model parameters. Manganese reduction half-reactions are both the most favorable and the most probable reactions (i.e., they occur in almost all of the parameter combinations sampled by the Monte Carlo algorithm). Generally, the calculated catabolic rates for manganese reduction are about two orders of magnitude higher than for iron oxides reduction, and the biomass growth rates associated with manganese reduction are about three orders of magnitude higher than the calculated rates for iron oxides reduction.

In LK, only the oxidation of hydrogen, methane and ammonium oxidation to molecular nitrogen can be coupled to the reduction of iron oxides to generate biomass. This is because the calculated catabolic rates of iron reduction coupled to ammonium oxidation to nitrite and nitrate in LK are negative, which indicates insufficient energy for net forward reaction and growth. Thus, no modeled iron reduction half-reactions are capable of supporting cell growth in LK.

In PC-3, ferrihydrite and amorphous iron oxyhydroxide oxidation are plausible biomass-generating reactions only when coupled to ammonium oxidation to nitrogen. In SG-1, methane oxidation can also be coupled to iron reduction.

## 4. Discussion

### 4.1. The presence of highly reactive iron oxides in the methanic zone

According to both calculated Δ*G*^*0*^_*r*_ values, catabolic rates and biomass growth rates, amorphous manganese oxide (MnO_2_) is the most favorable electron acceptor, then ferrihydrite (Fe(OH)_3_) and then amorphous iron oxyhydroxides (FeOOH). This electron acceptor sequence is the same for all modeled redox reactions at all sites. On the other hand, all other iron oxides, which are more crystalline, generate negative biomass growth rates. This indicates that they cannot support cell growth, even when involved in exergonic reactions (where Δ*G*_*r*_ < 0, but where there is insufficient energy for anabolism).
Ferrihydrite and amorphous iron oxyhydroxides are the most reactive iron oxides (Canfield, 1989; Poulton et al., 2004b). Therefore, they are expected to be scavenged by the microbial communities in the “classical” iron reduction zone, above the SMTZ. Furthermore, reactive iron oxides that have persisted through the zone of iron reduction (Figs. S3 and S4) are expected to be scavenged by reaction with sulfide (sum of all S^2-^, mostly in the form of H_2_S and HS^−^) in the zone of sulfate reduction, above and in the SMTZ (Canfield, 1989b; Kostka and Luther, 1995; Krom et al., 2002; Poulton, 2003). Nonetheless, iron extractions from the methanic zones of both the MedS and LK show that reactive iron oxides have survived the iron reduction and persisted through the sulfidic zones, remaining present even below the SMTZ (Tables S18-S20).

There are three scenarios that can explain the presence of reactive iron oxides in this zone: (1) Non-steady state diagenesis and high sedimentation rates (Rooze et al., 2016); (2) Redox recycling of iron in the methanic zone and in-situ formation of iron oxides; (3 Higher burial of iron oxides compared to the rate of sulfate reduction, which enables survival of the iron in forms other than FeS and pyrite below the SMTZ.

In the sediments of both study sites, steady state digenesis is observed (Adler et al., 2011; Amiel et al., 2020), ruling out the first alternative. The second alternative, redox recycling of iron, appears possible. Nitrite or nitrate (produced during anammox) can oxidize ferrous iron to generate highly reactive iron oxides, either biotically or abiotically (Sørensen and Thorling, 1991; Straub et al., 1996; Hauck et al., 2001; Kappler et al., 2005; Melton et al., 2014; Schaedler et al., 2018). Our calculations suggest that oxidation of ammonium by MnO_2_ to generate both nitrate and nitrite can sustain microbial growth (Figs. 2-7). Reaction of the nitrate and/or nitrite with ferrous iron phases may regenerate fresh and reactive ferric iron (oxyhydr)oxides (Jørgensen and Nelson, 2004). Both of these explanations for the presence of reactive iron (oxyhydr)oxides in the methanic zone require the survival of manganese oxides below the SMTZ. Another potential pathway for iron recycling in the methanic zone is through the authigenic precipitation of magnetite (Amiel et al., 2020). The third alternative, iron oxide deposition fluxes that exceed the supply of sulfate, could also explain the presence of highly reactive iron in the methanic zone. To test this possibility, we compared the total sulfur and total iron fluxes reaching the methanic zones of SG-1 and LK (Fe:S ratio) (Table 7; see Table S35 for sediment properties and Table S36 for relevant concentrations). Total sulfur (S) fluxes were calculated as the sum of dissolved sulfate (SO_4_^2-^) diffusion across the SMTZ and the accumulation of solid acid-volatile sulfur species AVS (iron sulfide minerals such as FeS, Fe_3_S_4_, mackinawite and greigite) and solid chromium-reducible sulfur (CRS – sulfur-containing minerals such as FeS_2_, ZnS and PbS). The total iron (Fe) fluxes were calculated as the sum of AVS, CRS and amorphous and poorly crystalline iron oxides (termed “Oxides 1”). We deduced that Fe:S > 0.5 means that there is not enough S to lock al Fe in pyrite, and that Fe:S >1 means that ther is not enough S to lock all Fe in AVS as well. Similarly, manganese oxides can also persist through their zone of reduction if their fluxes are higher than those of sulfate. To calculate the diffusive supply of sulfate, we considered no sulfate below the SMTZ in both LK and MedS, 0.1 mM of sulfate above the SMTZ of LK (7 cm below the sediment-water interface) (Adler et al., 2011), and 7 mM of sulfate above the SMTZ of the MedS (150 cm below the sediment-water interface) (Vigderovich et al., 2019). Our calculations clearly indicate that in both LK and station SG-1 in the MedS, the fluxes of total highly reactive iron were substantially higher than those of total sulfur during sediment deposition and diagenesis. Thus, due to high ratios of Fe:S fluxes (7.1 in LK and 6.5 in SG-1), the availability of sulfur was insufficient to sequester all iron in AVS or pyrite, and iron oxides could persist through the sulfate reduction zone. This allowed reactive iron (oxyhydr)oxides to reach the methanic zone and participate in the microbial dissimilatory processes within this zone. The reactivation and reduction of iron can occur for several reasons: (1) A switch in methanogens (such as *Methanosarcinales*) from methanogenesis to iron reduction (Eliani-Russak et al., 2023), likely facilitated by electron shuttles like methanophenazines; (2) The availability of methane as a substrate for Fe-AOM; (3) Recycling of iron oxides (as described above).

The presence of highly reactive iron oxides in methanic sediments is not surprising or new (Sivan et al., 2011; Vigderovich et al., 2019; Aromokeye et al., 2021) when considering total sulfur and iron fluxes and recognizing the possibility of iron redox cycling within anoxic sediments. This notion contrasts the common assumption, that highly reactive iron oxides are absent from deep sediments (Shi et al., 2007; Eliani-Russak et al., 2023), which must be tested in each environment.

**Table 7:** Fluxes of sulfate, AVS, CRS and amorphous/poorly crystalline iron oxides (Oxides 1) above the methanogenic zones of LK and the MedS (SG-1), expressed in units of (mol (cm total sediment)^−2^ sec^−1^)·10^13^. The Fe:S fluxes ratio is calculated as (Oxides 1 +AVS + CRS)/(SO_4_^2-^ + AVS + CRS).

| Specie | Flux<br>LK | Flux<br>SG-1 |
| --- | --- | --- |
| Sulfate | $5.1 \cdot 10^{-1}$ | 1.5 |
| AVS | $3.7 \cdot 10^{-3}$ | 4.0 |
| CRS | 3.2 | 3.3 |
| Oxides 1 | 3.6 | 5.9 |
| **Fe:S** | **1.8** | **1.5** |

### 4.2. The main iron and manganese oxide reduction processes in the methanic zone

Our results emphasize that negative Δ*G*_*r*_, even though necessary for biomass generation, cannot alone guarantee cell growth. The cascade of electron donors from the most to the least energetically favorable (according to Δ*G*_*r*_) is consistent among all modeled redox reactions at all sites (Fig. 1). In contrast, there is little correlation between the electron donor cascade in terms of energy yield (Fig. 1) and the electron donor cascade in terms of the actual catabolic rates and biomass growth rates (Figs. 2-7). Acetate oxidation, which is the most energetically favorable process (Fig. 1), generates low catabolic and biomass growth rates relative to other metabolisms (Figs. 2-7). This can be explained by the low concentrations of acetate in both sites, which are considered in the catabolic rate calculations. In contrast, ammonium oxidation to nitrogen, which generates considerably less energy than acetate oxidation on a per mole basis (Fig. 1), is one of the main metabolisms at both sites, due to the relatively high concentration of ammonium.

Further differences between the order of catabolic rates and biomass growth rates with difference electron donors at a specific station (Figs. 2-7) are due to the dependence of the biomass growth rates not only on catabolic rates, but also on Δ*G*_*r*_ and the biomass yield (*Y*_*m*_^*ED*^; see Eq. 18). This is why, for example, the model shows that methane oxidation in the methanic zone of the MedS at both stations (SG-1 and PC-3) is one of the most favorable oxidation reactions in terms of biomass growth rates, but in terms of catabolic rates it is less favorable than all modeled ammonium oxidation reactions. Note that the dissipated power (*P*_*d*_) and the number of cells per volume of sediment (*B*_*j*_), which also contribute to the biomass growth rate, are assigned different values in cases of methane oxidation than in all other cases, as a consequence of the assumption that only archaea oxidize methane (Tables 3S-5S). This explains why the catabolic rates and growth rates for methane oxidation do not always follow the same trend as the other electron donors’ oxidation half-reactions; for example, the biomass growth rate for manganese oxide reduction coupled to methane oxidation in station SG-1 is higher than the biomass growth rate when coupled to ammonium oxidation to nitrogen.

The nominal model shows that iron reduction is thermodynamically favorable but results in no iron reduction. This suggests that more favorable kinetic parameters (*V*_*max*_ and *K*_*s*_) than the mean values used in the nominal model are needed for these bioreactions to occur at the concentrations of the electron donors and acceptors in the study sites. In the Monte Carlo simulation, where kinetic parameters are drawn from relatively wide distributions, iron reduction does occur, but at a low probability (a small fraction of random parameter combinations). The observation of iron reduction in methanic sediments, including in LK and MedS (Sivan et al. 2011; Riedinger et al., 2014; Treude et al., 2014; Bar-Or et al., 2015, 2017; Egger et al., 2017), suggests that more favorable kinetic parameters may better represent the microbial communities in these environments than the culture-constrained average values used in the nominal model.

The fact that the Δ*G*_*r*_ values at all sites are similar for each bioreaction (Fig. 1), while catabolic and biomass growth rates vary among sites (Figs. 2-7), suggests that the kinetics of the bioreactions, rather than their thermodynamic drive, create the main differences in the dominance of metabolisms among sites. The compounds that contribute to the kinetic factor (*F*_*K*_) are only the electron donor and electron acceptor concentrations, while all compounds contribute to ΔG_r_ (including hydrogen as a reactant and all products).

The Δ*G*_*r*_ values are significantly lower when reducing amorphous manganese oxide, followed by a cascade of less energetically favorable iron oxides. This suggests that the main differences between the electron acceptors in catabolic and biomass growth rate are caused by their Δ*G*^*0*^_*f*_; for instance, the Δ*G*^*0*^_*f*_ of amorphous manganese oxide is much higher than these of iron oxides (Table S1), and so it requires less energy when consumed. In addition, the high sensitivity of the model to the *V*_*max*_ values of the electron acceptors, suggests that they create further differences between catabolic rates, which directly controls the biomass growth rates. Therefore, the favorability of electron acceptors in terms of biomass growth is mainly determined by their identity (which determines their Δ*G*^*0*^_*f*_ and *V*_*max*_), and is less dependent on their concentrations.

Catabolic rates and biomass growth rates for iron sulfide (FeS) oxidation were not calculated due to the lack of kinetic parameters. In our model, we assume that the concentrations of electron donors and acceptors are in quasi-steady state, meaning there is no direct connection between different bioreactions, as concentrations do not change over time due to microbial consumption. Therefore, the absence of FeS oxidation does not influence the other modeled bioreactions. Nevertheless, the calculated Δ*G*_*r*_ values when coupled to manganese reduction half-reaction are negative enough (Table 2; Fig. 1) to infer that this reaction is bioenergetically plausible. In contrast, the Δ*G*_*r*_ values when iron sulfide oxidation is couple to amorphous iron oxyhydroxide or ferrihydrite reduction are close to equilibrium at both sites (Table 2; Fig. 1), suggesting that these redox reactions cannot generate enough energy to support cell growth. All other modeled electron acceptors (lepidocrocite, goethite, hematite and magnetite in both the MedS and LK, and amorphous iron oxyhydroxide in the MedS generate positive Δ*G*_*r*_ values when coupled to iron sulfide oxidation. Hence, these reactions are thermodynamically impossible in the bulk sediment.

### 4.3. Relative community sizes in the methanic zone

Ideally, the biomass growth rate calculations for each microbial functional group would consider only the number of active cells within the group. However, since the calculations considered the total amount of archaea and bacteria in the methanic zone (including dead cells, as qPCR measures the presence of genes, not the activity of the organisms; Eq. 18), with no separation between functional groups, all calculated growth rates are expected to be higher than the actual biomass growth rates.

Assuming that the calculated biomass growth rates of the different functional groups are proportional to their abundance, the relative community sizes were calculated (Fig. 8). This calculation does not represent all possible cases, but instead relies on the median biomass growth rates obtained from to the Monte Carlo simulation. The functional groups were classified according to the electron donor consumed – acetate, hydrogen, methane and ammonium oxidizers to molecular nitrogen and to nitrite and nitrate. The relative population sizes were calculated assuming that only archaea can anaerobically oxidize methane, although the presence of *methylococcales*, an aerobic methanotrophic bacteria, has been detected in the methanic sediments of LK (Bar-Or et al., 2015; 2017; Elul et al., 2021). Bar-Or et al. (2015; 2017) suggested that small amounts of remnant oxygen in the deep sediments are consumed by these aerobic methanotrophs. Still, we decided to not consider the methanotrophic bacteria in our model, since the concentrations of oxygen available for methanotrophy in this zone are below detection (1 ppb).

It is important to note that microbial community sizes may vary due to changes in the concentrations of electron donors and acceptors at different depths. These variations can lead to the formation of micro-environments with distinct microbial communities. However, the Monte Carlo simulation accounts for these variations, providing a statistical representation of microbial community dynamics and their reaction rates across different environmental conditions. By incorporating such variations into the simulation, we can capture the impact of changing geochemical conditions on community structure without needing to explicitly model each micro-environment.

**Figure 8:**
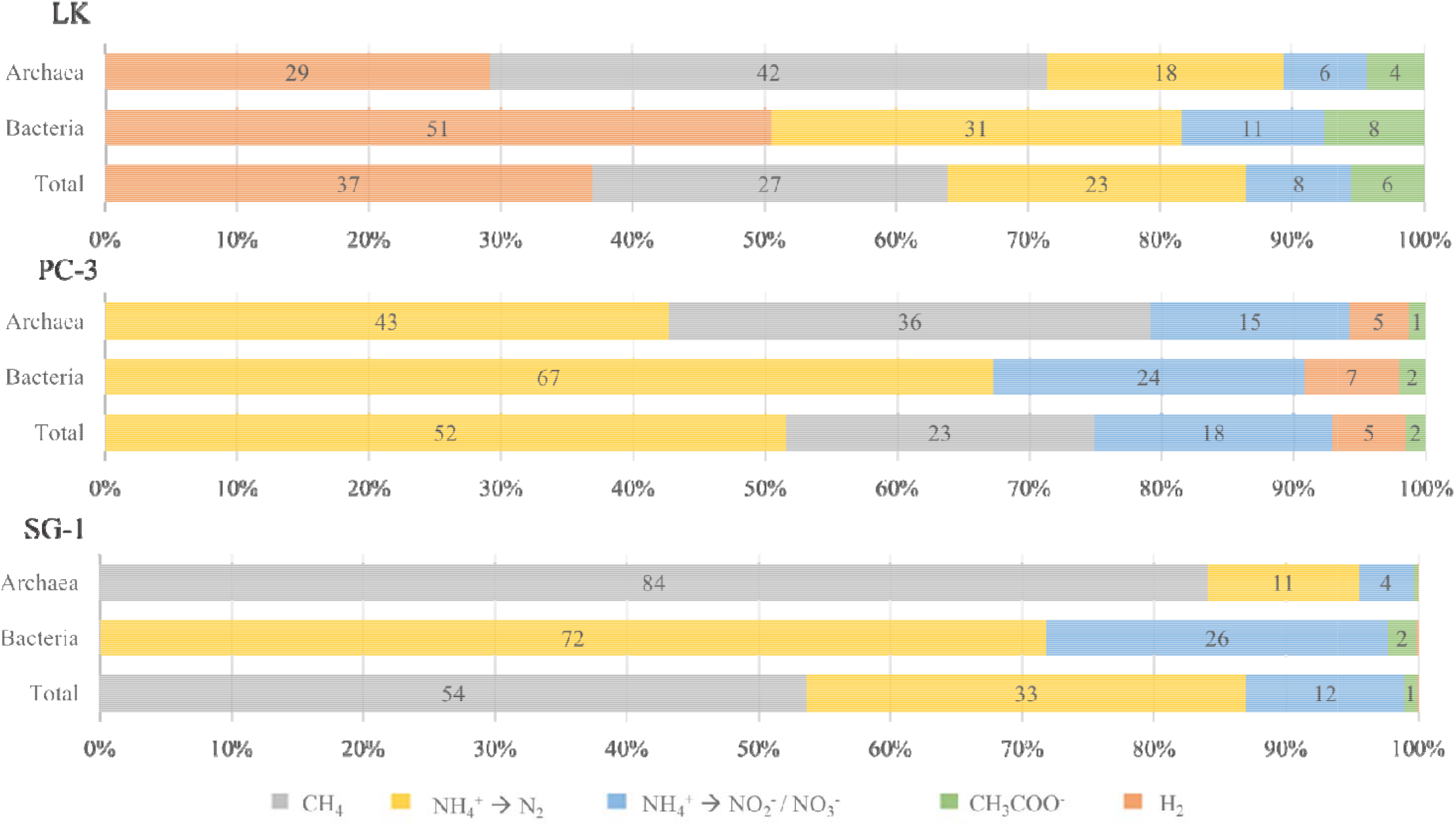
Relative functional group sizes (%) of both iron and manganese reducers in the methanogenic zones of LK and the MedS (PC-3 and SG-1), classified by electron donor, considering median biomass growth rates calculated by the Monte Carlo algorithm (Figs. 2, 4 and 6) and assuming a direct relation between population sizes and the calculated growth rates. At PC-3, only ammonium oxidizers to molecular nitrogen are present in the absence of manganese (Fig. 4; see Fig. S6 for relative abundances of iron reducers in the methanic zones of LK and SG-1). Iron sulfide (FeS) oxidizing functional groups are not included.

Table 8 presents the relative abundances of the modeled functional groups in LK and SG-1 according to metagenome sequencing (after Vigderovich et al., 2019; Elul et al., 2021). The microorganisms that were found in the natural environments were divided into functional groups considering their ability to oxidize organic matter, hydrogen, methane or ammonium, as described in the literature (Table S8). According to the table, organotrophs are the least abundant in both LK and the MedS. This aligns with the model results, which predict low actetotroph abundances (Fig. 8). In addition, the model anticipates a higher percentage of ammonium oxidizers at all sites, and also more methanotrophs in LK (we separate methanotrophs from other organotrophs because organic compounds that contain oxygen atoms have a very different ΔG_f_ than methane). There is a minor difference between observed and expected hydrogenotroph abundance in LK and the normalized abundance according to the metagenome sequencing. Another difference worth noting is the absence of methanotrophs in the MedS, and their low abundance in LK, while the model predicts high abundances. This is true not only for the median case, as the Monte Carlo simulations anticipate relatively high biomass growth rates when oxidizing methane while reducing amorphous manganese oxides at all stations. All differences between the model and the metagenome sequencing can be explained by the limited ability of metagenome sequencing to determine functional group abundances. This is because (1) certain microorganisms are capable of performing several metabolisms, (2) the ability of a microorganism to perform a certain metabolism does not guarantee its performance in a certain environment, (3) not all functional genes are known, which makes the sorting into functional groups less definitive, and (4) metagenome sequencing cannot identify all microorganisms present in a sample – about 30% of the metagenome in both LK and the MedS is unknown (Tables S6-S7).

**Table 8:** Functional group abundances determined by metagenome sequencing. The upper rows present relative abundances of possible iron/manganese reducers according to metagenome sequencing, sorted by electron donor consumption. Since certain microorganisms are capable of oxidizing several electron donors (Table S8), the sum of all functional groups’ relative abundances in both LK and SG-1 is higher than 100%. Non-shaded columns present relative abundances before normalizing to 100%, and gray-shaded columns present relative abundances normalized to 100%. The bottom rows present the proportion of archaea and bacteria among iron/manganese reducers.

| ED | LK, 25 cm<br>Iron reducers (%) | LK, normalized<br>Iron reducers (%) | SG-1, 400 cm<br>Iron reducers (%) | SG-1, normalized<br>Iron reducers (%) |
| --- | --- | --- | --- | --- |
| Ac/OM | 70.70 | 53.61 | 100.00 | 96.59 |
| H<sub>2</sub> | 41.39 | 31.39 | 1.03 | 1.00 |
| CH<sub>4</sub> | 6.59 | 5.00 |  |  |
| NH<sub>4</sub><sup>+</sup> | 13.19 | 10.00 | 2.50 | 2.41 |
| **Sum** | **131.87** | **100** | **103.53** | **100** |
| Archaea |  | 39.93 |  | 14.29 |
| Bacteria |  | 60.07 |  | 85.71 |

### 4.4. Total reaction rates and environmental significance

Total iron and manganese reduction rates in the methanic zones of the MedS and LK were calculated according to Eq. 1 for amorphous iron oxyhydroxide, ferrihydrite and amorphous manganese oxide reduction, assuming the metabolic processes in either iron or manganese reduction are similar (Fig. 9). The number of growing cells were calculated according to the cell counts (Table S3) and the median relative microbial functional group sizes obtained from the Monte Carlo simulation (Fig. 8). Similarly, the catabolic rates considered in this calculation are those calculated with the nominal model.

**Figure 9:**
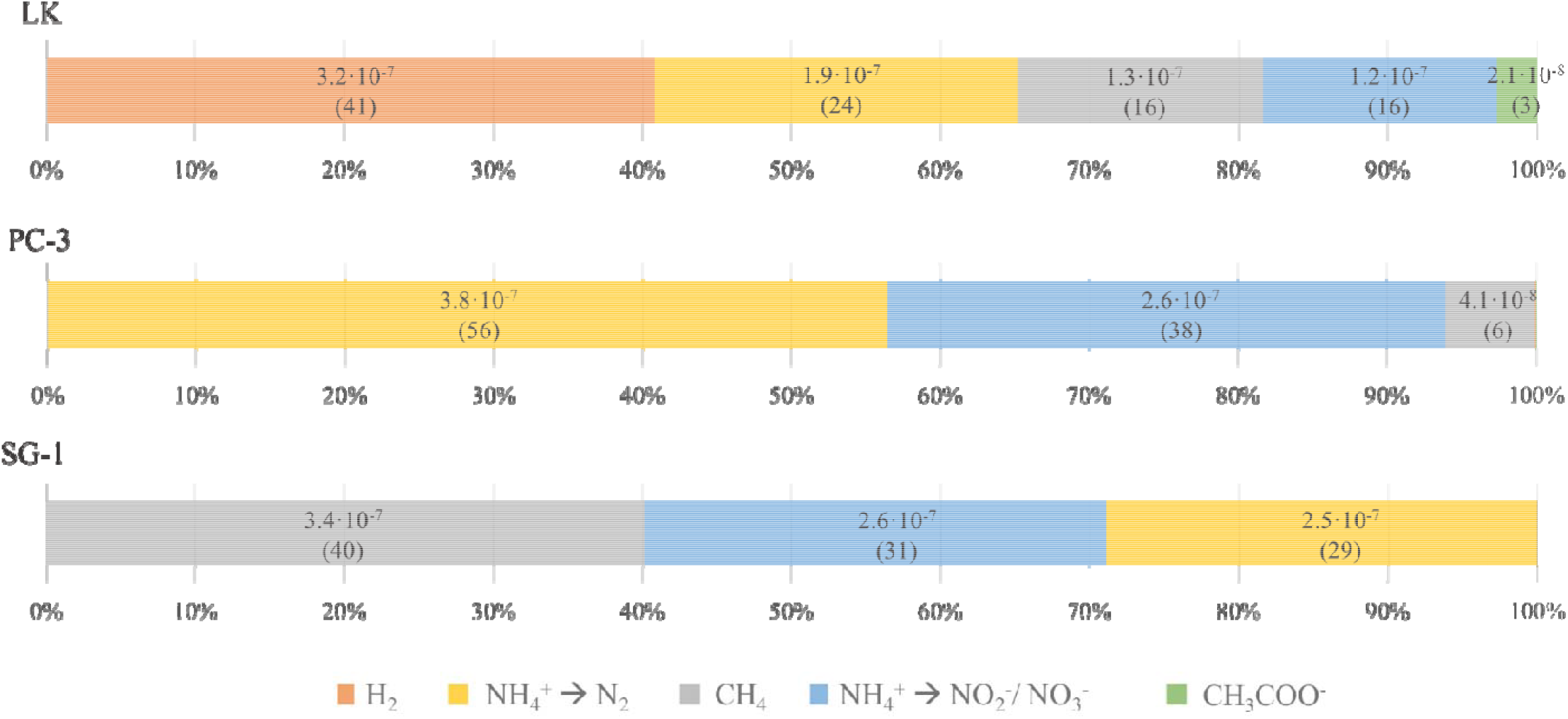
Iron and manganese oxide total reduction rates (mol oxide cm^−3^ day^−1^) in the methanic zones of LK and the MedS (PC-3 and SG-1), classified by electron donor, considering median biomass gro wth rates and catabolic rates calculated by the Monte Carlo algorithm (Figs. 2-3 and 6-7) and ass uming relative microbial population sizes that are directly proportional to the calculated growth rate s. At SG-1, the calculated total rates of acetate oxidation and hydrogen oxidation are 1.1·10^−10^ and 1.4 ·10^−12^ mol oxide cm^−3^ day^−1^, respectively. At PC-3, the calculated total rates of acetate oxidation and hyd rogen oxidation are 1.2·10^−10^ and 4.1·10^−10^ mol oxide cm^−3^ day^−1^, respectively. Ammonium oxi dation to nitrogen is the only plausible oxidation half-reaction at PC-3 in the absence of ma nganese, with a total rate of 2.8·10^−8^ mol oxide cm^−3^ day^−1^ (see Fig. S7 for total iron reduction rate in the methanic zones of LK and SG-1). In parentheses are the relative rates of each metabolic pat hway in percent (%). Iron sulfide (FeS) oxidizing functional groups are not included.

In LK, the biomass growth rates of all oxidation half-reactions coupled to manganese reduction are considerable, with acetate oxidation and ammonium oxidation to nitrate or nitrite being slightly slower than biomass synthesis based on hydrogen, methane or ammonium oxidation to nitrogen (Fig. 2). As shown in our model (Figs. 2-3), iron reduction in LK is most probably coupled to hydrogen or methane oxidation, while ammonium oxidation is the least probable. This agrees with previous observations of Fe^2+^ generation and deep AOM in the methanic zone of LK (Adler et al., 2011; Sivan et al., 2011; Bar-Or et al., 2015; Vigderovich et al., 2023).

In PC-3, on the other hand, ammonium oxidation to molecular nitrogen and methane oxidation, coupled to manganese reduction, are the most dominant metabolisms. The next dominant metabolism is ammonium oxidation to nitrite or nitrate. Hydrogen and acetate oxidation take place in the methanic zone of PC-3 according to the model, but their total rates are insignificant compared to the other metabolic pathways, as the catabolic rates of these metabolisms are considerably lower (Fig. 9). In SG-1, on the other hand, methane oxidation is the most dominant oxidation half-reaction, since the methane concentration in the methanic zone is high (Table S18). This is followed by ammonium oxidation to nitrate or nitrite and ammonium oxidation to molecular nitrogen, with hydrogen and acetate oxidation half-reactions being insignificant in this environment as well. An increase in nitrate and nitrite concentrations in the methanic zone of SG-1, and the presence of potential ammonium oxidazing microorganisms indicate ammonium oxidation in SG-1 were observed Our model shows that iron reduction in PC-3 coupled to ammonium oxidation to molecular nitrogen is the only plausible metabolic pathway that can explain the rise in Fe^2+^ concentrations (Figs. 6-7), while in SG-1 Fe-AOM is possible (but not certain) as well (Figs. 4-5). The model predicts Mn-AOM at both stations, assuming a similar ratio of manganese to total iron concertation to that of LK (manganese oxides were not measured in the MedS). Our findings suggest that manganese oxides play a more prominent role in the biogeochemistry of these environments compared to iron oxides. Thus, the fact that there are no indications for intensive AOM in the MedS methanogenic sediments (Yorshansky et al., 2022), as well as other marine sediments, such as in the South China Sea (Liang et al. 2022) and in the German Bight of the North Sea (Aromokeye et al., 2020), suggests lower manganese oxide concentrations in marine sediments than assumed here.

However, Yorshansky et al. (2022) did not detect AOM in natural samples or in samples with added amorphous iron. It is possible that if amorphous manganese oxides had been added to their experiments, AOM could have been detected, as manganese oxides might play a more significant role in AOM than previously considered in marine sediments.

It should be noted that even available iron extractions do not provide the exact concentrations of ferrihydrite and amorphous iron oxyhydroxide, as these phases are extracted together with lepidocrocite (Tables S18-S20). Other mechanisms that could decrease the availability of these reactive iron and manganese oxides to dissimilatory metabolisms are adsorption or incorporation of transition metals and heavy metals (e.g., Ni, Mo, Cu, Co, As, Zn and Pb), which stabilize the reactive iron and manganese oxides (Balistrieri and Murray, 1986; Burdige, 1993; Millward and Moore, 1982; Murray, 1975). Also, adsorption of phosphate to ferrihydrite (Wang et al., 2015, 2017) and manganese dioxide (Yao and Millero, 1996) may decrease the reactivity of these oxides. Hence, even bioenergetically plausible bioreactions might still not occur or become considerably less significant in natural environments.

In our model, we assume that the concentrations of electron donors and acceptors are in steady state, reflecting conditions that are already influenced by microbial consumption. This simplification means that we do not account for the temporal variation in these concentrations that would naturally occur as microbial reactions progress. In reality, concentrations would likely decrease over time, causing a corresponding drop in cell-specific rates according to Michaelis-Menten kinetics. While this assumption may lead to an overestimation of the total reaction rates and biomass, the Monte Carlo simulation accounts for variations in concentrations and kinetic parameters, providing statistical insights into the uncertainty and variability of the model’s predictions.

In our study, we calculate AOM rates of 3.4·10^−7^ mol cm^−3^ day^−1^ in SG-1 and 4.1·10^−8^ mol cm^−3^ day^−1^ in PC-3. These rates are significantly higher than the 0.65·10^−9^ mol cm^−3^ day^−1^ maximum AOM rate estimated by Dale et al. (2006) for coastal marine sediments. The higher rates in our study can be attributed to the site-specific nature of our model, which incorporates local geochemical and microbial data, along with the assumption of a quasi-steady state. While Dale et al. based their model on generalized conditions for coastal marine sediments, our approach factors in real methane gradients and electron acceptor availability, resulting in higher predicted AOM rates despite the lower methane concentrations in the MedS.

The methanotrophy rates by iron and manganese reduction in methanic sediments calculated in this study are compared to methanotrophy rates in marine and lake sediments calculated in previous studies in Table 9. Sivan et al. (2007) and Adler et al. (2011) calculated methanotrophy rates numerically according to sediment porewater methane profiles, as opposed to the bioenergetic approach taken here. The rates calculated for the MedS are about 4 orders of magnitude higher than the rates calculated numerically for the West African margin (Sivan et al., 2007). This may be explained by concurrent methanotrophy and methanogenesis in the studied sediments, such that high gross rates of methanotrophy and methanogenesis lead to the small net rates of methanotrophy calculated from the porewater profiles. In contrast with the large difference in the marine sites between bioenergetic and net measured methanotrophy rates, the median rates calculated for LK are only slightly higher than the median rate calculated numerically by Adler et al. (2011). The higher methanotrophy rates predicted by the bioenergetic model are expected to more accurately represent the actual bioreactions, as they focus specifically on the biogeochemical processes that drive methane oxidation.

**Table 9:** Typical methanotrophy rates (mol CH_4_ cm^−3^ day^−1^) in methanic sediments, coupled to the reduction of iron oxides (Fe-AOM) and manganese oxides (Mn-AOM). Total methanotrophy rates are also displayed. To obtain methanotrophy rates by iron and manganese oxide reduction, iron and manganese reduction rates calculated in this research were multiplied by 1/8 and 1/4, respectively (see balanced reactions in Table S11). Rates of iron and manganese oxide reduction reactions are calculated according to their ratio in the median case (Figs. 9 and S7). ODP: Ocean Drilling Program.

| Reference | Location | Fe-AOM rate | Mn-AOM rate | Total rate | Method |
| --- | --- | --- | --- | --- | --- |
| Sivan et al. (2007) | West African margin<br>ODP site 1081 | | | $1.38 \cdot 10^{-12}$ | Mass balance model |
| Adler et al. (2011) | Station A,<br>Lake Kinneret | | | $2.16 \cdot 10^{-8}$<br>$\pm 1.30 \cdot 10^{-8}$ | Mass balance model |
| Egger et al. (2015b) | Station US5B,<br>the Bothnian Sea | $3.62 \cdot 10^{-9}$<br>$\pm 2.47 \cdot 10^{-10}$ | | | Incubation experiments |
| Lenstra et al. (2023) | Station NB8,<br>the Bothnian Sea | $8.5 \cdot 10^{-9}$ | $1.5 \cdot 10^{-9}$ | $1.0 \cdot 10^{-8}$ | Bioenergetic model<br>(maximal nominal rates) |
| This study | Station PC-3, the<br>Mediterranean Sea | Unlikely<br>bioreaction | $1.0 \cdot 10^{-8}$ | $1.0 \cdot 10^{-8}$ | Bioenergetic model<br>(median rates) |
| This study | Station SG-1, the<br>Mediterranean Sea | $1.7 \cdot 10^{-9}$ | $8.3 \cdot 10^{-8}$ | $8.4 \cdot 10^{-8}$ | Bioenergetic model<br>(median rates) |
| This study | Station A,<br>Lake Kinneret | $6.5 \cdot 10^{-10}$ | $3.1 \cdot 10^{-8}$ | $3.2 \cdot 10^{-8}$ | Bioenergetic model<br>(median rates) |

Fe-AOM and Mn-AOM rates calculated in this study are also compared with those calculated for the coastal sediments of the Bothnian Sea (Egger et al., 2015b; Lenstra et al., 2023). It is important to note that both Lenstra et al. (2023) and this use bioenergetic methods, whereas Egger et al. (2015b) estimated Fe-AOM rates based on incubation experiments and not in in-situ conditions. Key differences between the bioenergetic models of Lenstra et al. (2023) and the model in this study include: (1) the model in this study accounts for a range of concentrations of relevant chemical species and a range of kinetic and thermodynamic parameters, implemented in a Monte Carlo simulation, while Lenstra et al. (2023) used nominal values for these parameters; and (2) in this model, the total reaction rates are derived from the median results of the Monte Carlo simulation, whereas Lenstra et al. (2023) reported the nominal values for the reaction rates calculated in the methanic zone of their study site. The cited rates in Table 9 are the maximal rates reported.

Overall, the Fe-AOM rates calculated for the methanic zone of the Bothnian Sea are somewhat higher than those for SG-1, which may be due to the higher methane concentrations at Station NB8 in the Bothnian Sea (approximately 6 mM CH_4_) compared to SG-1 (approximately 1.5 mM CH_4_). However, the Mn-AOM rates calculated in this study are significantly higher. This discrepancy may arise from differences in the manganese oxides considered in the two studies. Lenstra et al. (2023) calculated Mn-AOM rates assuming Mn(OH)_2_, while this model considered MnO_2_ as a manganese oxide. The manganese oxide used in this model has a much higher *V*_*max*_ value (1.30·10^−13^ mol cell^^−1^ day^−1^) compared to that in Lenstra et al. (2023) (1.62·10^−16^ mol cell^−1^ day^−1^).

Total iron and manganese reduction rates calculated in this study are compared to those reported by Lenstra et al. (2023) in Table 10. The iron oxides reduction rates calculated in this study are somewhat higher than those calculated for the Bothnian Sea, and the manganese reduction rates are higher by more than 2 orders of magnitude. In addition to the different manganese oxide considered in these two models, this discrepancy is likely due to the fact that the model in this study included more potential electron donors in the methanic zone – CH_4_, NH_4_^+^, H_2_, and CH_3_COO^−^, whereas Lenstra et al. (2023) considered O_2_, SO_4_^2-^, and CH_4_. Among these, only CH_4_ is present in significant concentrations in the methanic zone. We have added the table and this discussion to the manuscript.

**Table 10:**
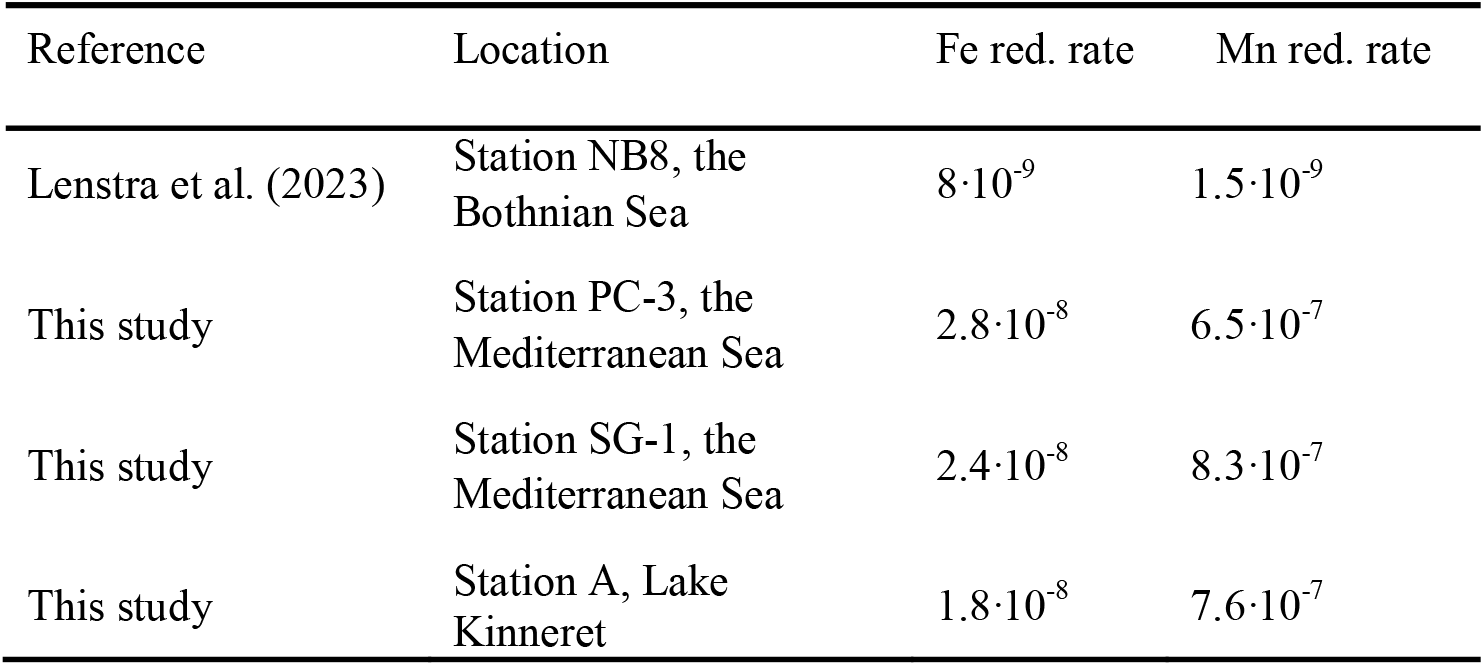
Calculated iron (Fe red.) and manganese (Mn red.) oxides reduction rates (mol oxide cm^−3^ day^−1^) in methanic sediments. Oxides reduction rates from this study are obtained from Figs 9 and S7.

## Conclusions

We used theoretical bioenergetic methods to develop a model to assess the dominance of microbial iron and manganese reduction processes and their effect on the geochemistry of methanic sediments in two lacustrine and marine natural study sites. This was done by calculating the microbial community sizes and total bioreaction rates, considering the expected catabolic and biomass growth rates with nominal parameter values constrained by field measurements and literature reports of laboratory experiments. While the geochemical conditions in these environments are more complex than considered in our model, as the diagenetic journey of electron donors and acceptors is not fully captured, we focused specifically on the biogeochemical reactions that change the concentrations of chemical species in the methanic zone. The Monte Carlo simulation accounts for variations in concentrations, as well as kinetic and thermodynamic parameters, and provides statistical insights into the variability and uncertainty of the model predictions. This approach helps to isolate the key biogeochemical processes within the methanic zone, despite the broader complexity of diagenetic processes throughout the sediment column.

Our model results show that amorphous manganese oxide (MnO_2_) is the most favorable electron acceptor in all cases examined, due to its very low Δ*G*^*0*^_*f*_ and high maximal rate capacities for its microbial reduction. The concentration of MnO_2_ was found to have a smaller effect on the catabolic and biomass growth rates. Thus, even when present at low concentrations compared to iron (oxyhydr)oxides, amorphous MnO_2_ is expected to play a significant role in the biogeochemistry of methanic sediments. This highlights an important point: while more iron oxide extractions are performed in most studies, manganese oxides are often overlooked, despite their potential significance. The higher availability of iron oxides in sediment samples may skew the focus toward Fe-based processes, potentially underestimating the role of manganese oxides in methane oxidation and other biogeochemical cycles in marine and lacustrine environments.

Ferrihydrite and amorphous iron oxyhydroxide are the only two iron oxides that were shown by our model to be able to support cell growth. Iron oxide reduction was found to be most probably coupled to hydrogen or methane oxidation in LK, and to ammonium oxidation to N_2_ in the MedS. The significance of Fe-AOM in the lake sediments, in contrast to its apparent absence in methane-depleted marine sediments, agrees with previous observations. However, Fe-AOM is possible (but not certain) in methane-enriched marine sediments. Our study reemphasizes that negative Δ*G*_*r*_ values are not a sufficient condition for net forward reaction and growth, and that reaction kinetics and microbial energetic needs must also be considered.

We found that considering both Mn and Fe bioenergetics and kinetics, ammonium oxidation is expected to be dominant at all sites, while hydrogen and acetate (or more generally – volatile organic acids) oxidation is expected to be significant only in LK, where their concentrations are higher. Methane oxidation coupled to manganese reduction is also possible at all sites.

In all cases, the Δ*G*_*r*_ showed low sensitivity to the compounds’ concentrations, which suggests that differences in similar microbial community sizes and the corresponding total bioreaction rates among different environments are predominantly the consequence of kinetic controls, affected by electron donor concentrations and bioreaction kinetic properties, rather than thermodynamic controls.

## Data availability

Data are available through Mendeley Data at https://doi.org/10.17632/9fdtnzgftr.1.

## CRediT authorship contribution statement

**Racheli Neumann Wallheimer:** Writing - Original Draft, Visualization, Data Curation, Investigation, Validation, Software, Methodology, Formal analysis. **Orit Sivan**: Writing - Review & Editing, Resources, Supervision, Project administration, Funding acquisition, Validation, Methodology, Conceptualization. **Itay Halevy:** Writing - Review & Editing, Supervision, Investigation, Validation, Methodology, Conceptualization.

## Declaration of competing interest

The authors declare that they have no known competing financial interests or personal relationships that could have appeared to influence the work reported in this paper.

## Acknowledgments

This project received funding from the European Research Council (ERC) under the European Union’s Horizon 2020 research and innovation program (Grant No. 818450 to OS). The authors thank Alexey Kamyshny, Almog Gafni, Valeria Boyko, Efrat Eliani Russak and Netta Gershon from Ben-Gurion University of the Negev, Hanni Vigderovich from the University of Bristol, Alice Bosco-Santos from the University of Lausanne and Yakar Zemach from the Hebrew University of Jerusalem, for the help in this research.

## Appendix A. Supplementary material

Accompanying information, including: 1) locations of study sites, 2) thermodynamic parameters, 3) standard Gibbs free energies for oxidation of carbonic electron donors, 4) microbiological data, 5) balanced redox reactions, 6) Gibbs free energies of net redox reactions in natural environments, 7) kinetic parameters, 8) model’s input for concentrations and activities in the methanic zone, 9) nominal catabolic rates, 10) nominal biomass growth rates, 11) relative functional group sizes of iron reducers and total iron reduction rates in Lake Kinneret, 12) sensitivity analysis, 13) standard deviations of normal distributed parameters, and 14) sediment properties, are available as a single file.

## Notes

https://doi.org/10.17632/9fdtnzgftr.1

### Competing Interest Statement

The authors have declared no competing interest.

