## Supplementary Material for "Modeling the Controls on Microbial Iron and Manganese Reduction in Methanic Sediments"

Number of pages: 42

Number of tables: 35
Number of figures: 14

1. **Study sites**

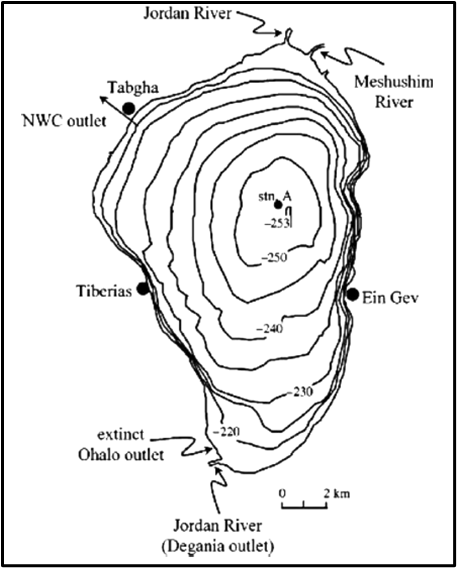

**Figure S1:** The location of the sampling station A in Lake Kinneret (Sea of Galilee). Geochemical and microbiological data collected from this station in previous studies are considered in the bioenergetic model developed in this study (map taken from Hambright et al., 2004).

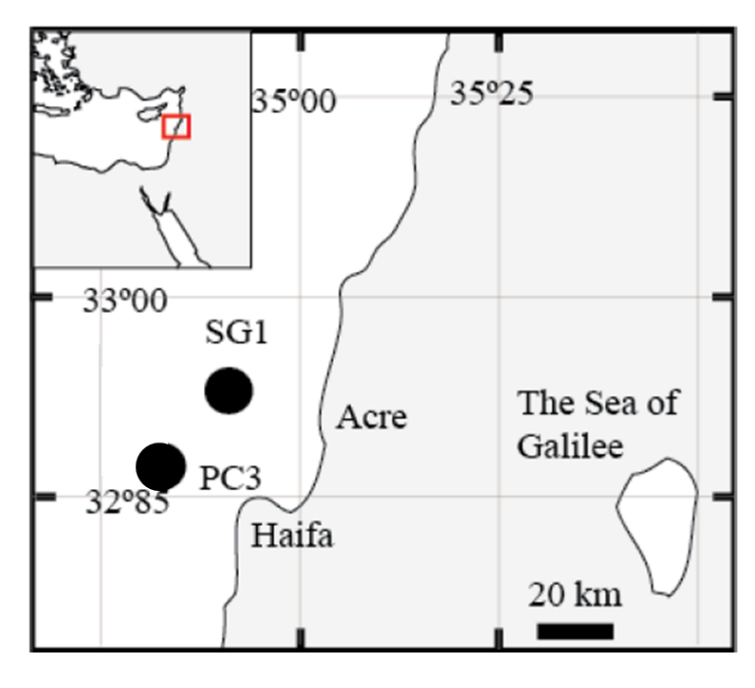

**Figure S2:** The location of the sampling stations SG-1 and PC-3, at the Eastern Mediterranean continental shelf. Geochemical and microbiological data collected from this station in previous studies are considered in the bioenergetic model developed in this study (map taken from Wurgaft et al., 2019).

2. **Thermodynamic parameters**

| **Reference** | **∆*H⁰_f_*** | **∆*G⁰_f_*** | **Specie** |
| --- | --- | --- | --- |
| Alberty (1998) | -486.83 | -247.82 | CH_3_COO^-^ (l) |
| Wagman et al. (1965) | -89.04 | -34.39 | CH_4_ (aq) |
| Wagman et al. (1982) | -823 | -696.5 | Fe(OH)_3_ (s) |
| Wagman et al. (1969) | -89.1 | -78.87 | Fe^2+^(aq) |
| Wagman et al. (1982) | -824.2 | -742.2 | Fe_2_O_3_ (s) |
| Wagman et al. (1982) | -1118.4 | -1015.4 | Fe_3_O_4_ (s) |
| Stumm and Morgan (1996) | -552 (assumed) | -462 | FeOOH (amorphous) (s) |
| Robie and Hemingway (1995) | -559.3 | -488.6 | α-FeOOH (goethite) (s) |
| Hiemstra (2015) | -552 | -482.7 | γ-FeOOH (lepidocorocite) (s) |
| Wagman et al. (1982) | -100 | -100.4 | FeS (s) |
| Wagman et al. (1982) | 0 | 0 | H^+^(aq) |
| Wagman et al. (1982) | -4.18 | +17.57 | H_2_ (aq) |
| Wagman et al. (1982) | -285.83 | -237.18 | H_2_O (l) |
| Wagman et al. (1982) | -20.6 | -27.83 | H_2_S (aq) |
| Wagman et al. (1982) | -691.11 | -586.77 | HCO_3_^-^(aq) |
| Wagman et al. (1982) | -220.8 | -228.1 | Mn^2+^ (aq) |
| Wagman et al. (1982) | -520 | -465.14 | MnO_2_ (s) |
| Calculated (*) | -10.29 | +10.85 | N_2_ (aq) |
| Wagman et al. (1982) | 0 | 0 | N_2_ (g) |
| Wagman et al. (1982) | -132.5 | -79.31 | NH_4_^+^ (aq) |
| Wagman et al. (1971) | -104.6 | -37.2 | NO_2_^-^ (aq) |
| Wagman et al. (1971) | -207.3 | -111.3 | NO_3_^-^(aq) |
| Wagman et al. (1982) | 0 | 0 | S^0^ (s) |

**Table S1:** Standard formation Gibbs free energies (∆*G⁰_f_*) (kJ mol^-1^) and enthalpies
(∆*H⁰_f_*) (kJ mol^-1^) of relevant compounds in 298 K and 1 atm.

**(*) Calculation of thermodynamic parameters of N_2_(aq):**

According to Broecker and Peng (1983):

$K_{H} \left( N_{2}\left( aq \right) \right), 273.15 K=8.19\cdot{10}^{-4} mol L^{-1} atm^{-1}$

$$K_{H} \left( N_{2}\left( aq \right) \right), 297.15 K=5.22\cdot{10}^{-4} mol L^{-1} atm^{-1}$$

Calculation of equilibrium constants:

$$K_{H}=\frac{K}{R\cdot T}, R=8.314472 J K^{-1}mol^{-1}= 0.082057 L atm K^{-1}mol^{-1}$$

$$K \left( N_{2}\left( aq \right) \right),273.15 K= 8.19\cdot{10}^{-4}\cdot0.082057\cdot273.15=1.84\cdot{10}^{-2}$$

$$K \left( N_{2}\left( aq \right) \right), 297.15 K=5.22\cdot{10}^{-4}\cdot0.082057\cdot297.15 =1.27\cdot{10}^{-2}$$

Calculation of Gibbs free energies in 273 K and 797 K:

$$\Delta G_{f}^{0}= -RTlnK$$

$$\Delta G_{f}^{0}\left( N_{2}\left( aq \right) \right), 273.15 K= -8.314472\cdot273.15\cdot\ln\left( 1.84\cdot{10}^{-2} \right)=9.08 kJ mol^{-1}$$

$$\Delta G_{f}^{0}\left( N_{2}\left( aq \right) \right), 297.15 K= -8.314472\cdot297.15\cdot\ln\left( 1.27\cdot{10}^{-2} \right)=10.78 kJ mol^{-1}$$

Calculation of Gibbs free energies in 298 K, assuming a linear proportion between Gibbs free energy and temperature:

$$\Delta G_{f}^{0}\left( N_{2}\left( aq \right) \right), 298 K=\Delta G_{f}^{0}\left( N_{2} \right), 273 K+ \frac{298-273}{297-273}\cdot\left( \Delta G_{f}^{0}\left( N_{2} \right),297 K-\Delta G_{f}^{0}\left( N_{2} \right), 273 K \right)=9.08+\frac{298-273}{297-273}\cdot\left( 10.78-9.08 \right)=10.85 kJ mol^{-1}$$

Calculation of the enthalpy (assuming constant enthalpy):

$$\frac{\Delta G\left( T_{2} \right)}{T_{2}} - \frac{\Delta G\left( T_{1} \right)}{T_{1}}= \Delta H\left( \frac{1}{T_{2}}-\frac{1}{T_{1}} \right) \to\Delta H= \frac{\frac{\Delta G\left( T_{2} \right)}{T_{2}} - \frac{\Delta G\left( T_{1} \right)}{T_{1}}}{\frac{1}{T_{2}}-\frac{1}{T_{1}}}$$

$$\Delta H\left( N_{2}\left( aq \right) \right)= \frac{\frac{10.78}{297.15}- \frac{9.08}{273.15}}{\frac{1}{297.15}-\frac{1}{273.15}}= -10.29 kJ mol^{-1}$$

3. **Standard Gibbs free energies for oxidation of carbonic electron donors**

| **Half-reaction** | **ΔG^0’^_ED_** | **Reference** |
| --- | --- | --- |
| $\frac{1}{8} HC{O_{3}^{-}}_{\left( aq \right)}+ \frac{9}{8}H_{\left( aq \right)}^{+}+e^{-}\to\frac{1}{8}C{H_{4}}_{\left( aq \right)}+\frac{3}{8}H_{2}O$ | +25.06 | Calculated (**) |
| $\frac{1}{4} HC{O_{3}^{-}}_{\left( aq \right)}+ \frac{9}{8}H_{\left( aq \right)}^{+}+e^{-}\to\frac{1}{8}CH_{3}COO_{\left( aq \right)}^{-}+\frac{1}{2}H_{2}O$ | +26.90 | Dale et al. (2006) |

**Table S2:** Standard Gibbs free energies (kJ mol^-1^) in biologically neutral pH conditions of the half-reactions for oxidation of various electron donors in 298 K and 1 atm.

**(**) Calculation of ΔG^0’^_ED_ of CH_4_(aq):**

Correction of standard Gibbs free energies to biologically neutral pH conditions:
 $\Delta G_{r}^{0^{'}}= \Delta G_{r}^{0} -RTln\left( Kw^{0.5} \right)$

where:

$$R=8.314472 J K^{-1}mol^{-1} , T=298.15 K, K_{w}={10}^{-14}$$

Calculation of standard Gibbs free energy of methane oxidation half-reaction:

$$\Delta G_{r}^{0}= \frac{1}{8}\cdot\Delta G_{f}^{0}\left( C{H_{4}}_{\left( aq \right)} \right)+\frac{3}{8}\cdot\Delta G_{f}^{0}\left( H_{2}O \right)-\frac{1}{8}\cdot\Delta G_{f}^{0}\left( {{HCO}_{3}^{-}}_{\left( aq \right)} \right)-\frac{9}{8}\cdot\Delta G_{f}^{0}\left( H_{\left( aq \right)}^{+} \right)=\frac{1}{8}\cdot\left( -34.39 \right)+\frac{3}{8}\cdot\left( -237.18 \right)-\frac{1}{8}\cdot\left( -586.77 \right)-\frac{9}{8}\cdot0= -19.895 kJ mol^{-1}$$

Correction:

$$\Delta G_{r}^{0^{'}}= -19.895-\frac{8.314472}{1000}\cdot298.15\cdot\ln\left( \left( {10}^{-14} \right)^{0.5} \right)=25.06 kJ mol^{-1}$$

4. **Microbiological data**

| **Microorganism** | **B (cell (gr tsed)^-1^)** | **B (cell (cm tsed)^-3^)** | **Reference** |
| --- | --- | --- | --- |
| Archaea | 4.81∙10^6^ | 5.38∙10^6^ | Vigderovich et al. (2019) |
| Bacteria | 2.72∙10^6^ | 3.05∙10^6^ | Vigderovich et al. (2019) |

**Table S3:** Cell counts in SG-1 from qPCR analysis. The example was taken from 400 cm below the sediment-water interface in station SG-1. ρ_tsed_ = 1.1192 gr tsed/cm^3^ tsed.

| **Microorganism** | **Relative ratio (%)** | **Reference** |
| --- | --- | --- |
| Archaea | 38.17 | Bar-Or et al. (2015) |
| Bacteria | 61.83 | Bar-Or et al. (2015) |

**Table S4:** Total archaea/bacteria ratio in station A in LK from SILVA ngs pipeline RNA sequence analysis. The example was taken from 29-32 cm below the sediment-water interface.

| **Environment** | **P_d_ (kJ cm^-3^ day^-1^)  archaea + bacteria** | **P_d_ (kJ cm^-3^ day^-1^)  archaea** |
| --- | --- | --- |
| MedS | 2.62∙10^-7^ | 1.67∙10^-7^ |
| LK | 2.95∙10^-7^ | 1.13∙10^-7^ |

**Table S5:** Dissipated power (P_d_) of the total microorganism community and archaea only, considering Tables 3S and 4S, and Eq. 14.

| **Taxa** | **Abundance (%)** |
| --- | --- |
| Acidobacteria Aminicenantia Aminicenantales | 2.6 |
| Acidobacteria Thermoanaerobaculia Thermoanaerobaculales | 1.0 |
| Euryarchaeota Methanomicrobia Methanomicrobiales | 5.4 |
| Euryarchaeota Methanomicrobia Methanosarcinales | 1.8 |
| Euryarchaeota Thermococci Methanofastidiosales | 2.3 |
| Euryarchaeota Thermoplasmata Methanomassiliicoccales | 1.3 |
| Firmicutes Clostridia Clostridiales | 0.4 |
| Nitrospirae Thermodesulfovibrionia uncultured | 3.7 |
| Proteobacteria Deltaproteobacteria Desulfarculales | 1.2 |
| Proteobacteria Deltaproteobacteria MBNT15 | 2.1 |
| Proteobacteria Deltaproteobacteria Myxococcales | 1.1 |
| Proteobacteria Deltaproteobacteria Sva0485 | 7.0 |
| Proteobacteria Deltaproteobacteria Syntrophobacterales | 2.2 |
| Non-iron reducers | 35.2 |
| Others | 32.7 |

**Table S6:** Relative abundance of possible iron reducers in the methanogenic sediments of LK. The example was taken from ̴ 25 cm below the sediment-water interface (Elul et al., 2021).

| **Taxa** | **Abundance (%)** |
| --- | --- |
| Actinobacteria; Acidomicrobia; Acidimicrobials; Acidimicrobiaceae; uncultured bacterium | 2.5 |
| Nitrospirae; Nitrospira; Nitrispirales; Nitrospiraceae; uncultured bacterium | 1.0 |
| Proteobacteria; Deltaproteobacteria | 6.2 |
| Non-iron reducers | 62.6 |
| Others | 27.9 |

**Table S7:** Relative abundance of possible iron reducers in the methanogenic sediments of the MedS. The example was taken from 400 cm below the sediment-water interface (Vigderovich et al., 2019).

| **Domain** | **Phylum** | **Class** | **Order** | **Genus** | **Specie** | **ED** | **Reference** |
| --- | --- | --- | --- | --- | --- | --- | --- |
| Archaea | Euryarchaeota | Methanomicrobia | Methanomicrobiales | Methanolinea |  | H_2_ | Borrel et al. (2011) |
|  |  |  |  | Methanoregula |  | H_2_ | Borrel et al. (2011) |
|  |  |  | Methanosarcinales |  |  | CH_4_ | Ettwig et al. (2016) |
|  |  |  |  | Methanosaeta |  | Ac  H_2_ | Jetten et al. (1990)  Yamada et al. (2014) |
|  | Nitrospirae | Thermodesulfovibrionia | Thermodesulfovibrionales | Thermodesulfovibrio |  | H_2_ OM | Bar-Or et al. (2015)  Sekiguchi et al. (2008) |
| Bacteria | Acidobacteria |  |  |  |  | OM NH_4_^+^ | Huang and Jaffé (2015) |
|  | Actinobacteria |  |  |  |  | OM, NH_4_^+^ | Huang and Jaffé (2015) |
|  | Proteobacteria (Pseudomonadota) | Deltaproteobacteria (Desulfuromonadota) | Geobacterales | Geobacter |  | Ac  H_2_ OM | Sung et al. (2006)  Speers and Reguera (2012)  Wagner et al. (2012)  Bar-Or et al. (2015)  Holmes et al. (2016) |
|  |  |  | Candidatus  Acidulodesulfobacterales (sva0485) |  |  | H_2_S  OM | Tan et al. (2019)  Vigderovich et al. (2019) Elul et al. (2021) |
|  |  |  | Syntrophobacterales |  |  | H_2_ | Elul et al. (2021) |
|  |  |  | MBNT15  (candidate phylum) |  |  | OM | Begmatov et al. (2022) |
|  |  |  | Myxococcales |  |  | OM | Sanford et al. (2016) |
|  | Firmicutes | Bacilli | Bacillales | Mesobacillus | Bacillus thioparans | S_2_O_3_^2-^ | Pérez-Ibarra et al. (2007) |
|  |  | Clostridia | Eubacteriales | Clostridium | Clostridium tunisiense | S^0^ | Thabet et al. (2004) |

**Table S8:** Possible iron reducers found in the MedS and LK and the possible electron donors oxidized by each taxon mentioned, as suggested in literature. Ac denotes acetate (CH_3_COO^-^) and OM denotes organic matter (other than methane and acetate). In the model developed in this research, acetate represents all organic matter (besides methane). A detailed description of electron donors consumed by various *Deltaproteobacteria* can be found in Table 11.1 in Sanford et al. (2016). More information about abundant microorganisms in LK can be found in the supplement of Bar-Or et al. (2015).

5. **Balanced redox reactions**

All redox reactions were normalized to the transfer of one electron.

| Electron Acceptor | Reaction | ΔG_r_°  @ 25°C | ΔG_r_°  @ 14°C | ΔH_r_° |
| --- | --- | --- | --- | --- |
| Amorphous manganese | $\frac{1}{2}MnO_{2}+\frac{1}{8}CH_{3}COO^{-}+\frac{7}{8}H^{+}\to\frac{1}{2}Mn^{2+}+\frac{1}{4}HC{O_{3}}^{-}+{\frac{1}{2}H}_{2}O$ | -115.79 | -115.40 | -105.46 |
| Magnetite | $\frac{1}{2}Fe_{3}O_{4}+\frac{1}{8}CH_{3}COO^{-}+\frac{23}{8}H^{+}\to\frac{3}{2}Fe^{2+}+\frac{1}{4}HC{O_{3}}^{-}+{\frac{3}{2}H}_{2}O$ | -109.45 | -111.09 | -153.79 |
| Ferrihydrite | $Fe\left( OH \right)_{3}+\frac{1}{8}CH_{3}COO^{-}+\frac{15}{8}H^{+} \to Fe^{2+}+\frac{1}{4}HC{O_{3}}^{-}+\frac{5}{2}H_{2}O$ | -82.09 | -83.32 | -115.34 |
| Amorphous iron | $FeOOH+\frac{1}{8}CH_{3}COO^{-}+\frac{15}{8}H^{+}\to$ $Fe^{2+}+\frac{1}{4}HC{O_{3}}^{-}+\frac{3}{2}H_{2}O$ | -88.36 | -87.97 | -77.99 |
| Lepidocrocite | $\gamma-FeOOH+\frac{1}{8}CH_{3}COO^{-}+\frac{15}{8}H^{+}\to Fe^{2+}+\frac{1}{4}HC{O_{3}}^{-}+\frac{3}{2}H_{2}O$ | -67.66 | -68.04 | -77.99 |
| Goethite | $\alpha-FeOOH+\frac{1}{8}CH_{3}COO^{-}+\frac{15}{8}H^{+}\to Fe^{2+}+\frac{1}{4}HC{O_{3}}^{-}+\frac{3}{2}H_{2}O$ | -61.66 | -61.98 | -70.59 |
| Hematite | $\frac{1}{2}Fe_{2}O_{3}+\frac{1}{8}CH_{3}COO^{-}+\frac{15}{8}H^{+}\to Fe^{2+}+\frac{1}{4}HC{O_{3}}^{-}+H_{2}O$ | -60.67 | -61.19 | -74.97 |

**Table S9:** Balanced acetate (CH_3_COO^-^) oxidation reactions, standard Gibbs free energies (kJ mol^-1^) and enthalpies (kJ mol^-1^) in 298 K and 287 K, considering a pressure of 1 atm.

| Electron Acceptor | Reaction | ΔG_r_°  @ 25°C | ΔG_r_°  @ 14°C | ΔH_r_° |
| --- | --- | --- | --- | --- |
| Amorphous manganese | $\frac{1}{2}MnO_{2}+\frac{1}{8}CH_{4}+\frac{7}{8}H^{+} \to$ $\frac{1}{2}Mn^{2+}+\frac{1}{8}HC{O_{3}}^{-}+{\frac{5}{8}H}_{2}O$ | -127.45 | -127.69 | -134.14 |
| Magnetite | $\frac{1}{2}Fe_{3}O_{4}+ \frac{1}{2}{H_{2}}/{CO_{2}+3H^{+} \to\frac{3}{2}Fe^{2+}+2H_{2}O}$ | -93.75 | -95.60 | -144.02 |
| Ferrihydrite | $Fe\left( OH \right)_{3}+ \frac{1}{2} {H_{2}}/{CO_{2}}+2H^{+} \to$  $Fe^{2+}+3H_{2}O$ | -102.70 | -103.39 | -121.5 |
| Amorphous iron | $FeOOH+\frac{\frac{1}{2}H_{2}}{CO_{2}}+2H^{+}\to Fe^{2+}+2H_{2}O$ | -100.02 | -100.26 | -106.67 |
| Lepidocrocite | $\gamma-FeOOH+\frac{\frac{1}{2}H_{2}}{CO_{2}}+2H^{+}\to Fe^{2+}+2H_{2}O$ | -79.32 | -80.32 | -106.67 |
| Goethite | $\alpha-FeOOH+\frac{\frac{1}{2}H_{2}}{CO_{2}}+2H^{+}\to Fe^{2+}+2H_{2}O$ | -73.32 | -74.27 | -99.27 |
| Hematite | $\frac{1}{2}Fe_{2}O_{3}+ \frac{1}{2}{H_{2}}/{CO_{2}+2H^{+} \to Fe^{2+}+ \frac{3}{2}H_{2}O}$ | -72.33 | -73.48 | -103.655 |

**Table S10:** Balanced hydrogen (H_2_) oxidation reactions, standard Gibbs free energies
(kJ mol^-1^) and enthalpies (kJ mol^-1^) in 298 K and 287 K, considering a pressure of 1 atm.

| Electron Acceptor | Reaction | ΔG_r_°  @ 25°C | ΔG_r_°  @ 14°C | ΔH_r_° |
| --- | --- | --- | --- | --- |
| Amorphous manganese | $\frac{1}{2}MnO_{2}+\frac{1}{8}CH_{4}+\frac{7}{8}H^{+} \to\frac{1}{2}Mn^{2+}+\frac{1}{8}HC{O_{3}}^{-}+{\frac{5}{8}H}_{2}O$ | -98.77 | -98.97 | -104.41 |
| Magnetite | $\frac{1}{2}Fe_{3}O_{4}+\frac{1}{8}CH_{4}+\frac{23}{8}H^{+} \to\frac{3}{2}Fe^{2+}+\frac{1}{8}HC{O_{3}}^{-}+\frac{13}{8}H_{2}O$ | -65.07 | -66.89 | -114.29 |
| Ferrihydrite | $Fe\left( OH \right)_{3}+\frac{1}{8}CH_{4}+\frac{15}{8}H^{+} \to\frac{1}{8}HC{O_{3}}^{-}+Fe^{2+}+\frac{21}{8}H_{2}O$ | -74.02 | -74.67 | -91.77 |
| Amorphous iron | $FeOOH+\frac{1}{8}CH_{4}+\frac{15}{8}H^{+}\to$ $Fe^{2+}+\frac{1}{8}HC{O_{3}}^{-}+\frac{13}{8}H_{2}O$ | -71.34 | -71.54 | -76.94 |
| Lepidocrocite | $\gamma-FeOOH+\frac{1}{8}CH_{4}+\frac{15}{8}H^{+}\to$ $Fe^{2+}+\frac{1}{8}HC{O_{3}}^{-}+\frac{13}{8}H_{2}O$ | -50.64 | -51.61 | -76.94 |
| Goethite | $\alpha-FeOOH+\frac{1}{8}CH_{4}+\frac{15}{8}H^{+}\to$ $Fe^{2+}+\frac{1}{8}HC{O_{3}}^{-}+\frac{13}{8}H_{2}O$ | -44.64 | -45.55 | -69.54 |
| Hematite | $\frac{1}{2}Fe_{2}O_{3}+\frac{1}{8}CH_{4}+\frac{15}{8}H^{+} \to$ $Fe^{2+}+\frac{1}{8}HC{O_{3}}^{-}+\frac{9}{8}H_{2}O$ | -43.65 | -44.76 | -73.93 |

**Table S11:** Balanced methane (CH_4_) oxidation reactions, standard Gibbs free energies
(kJ mol^-1^) and enthalpies (kJ mol^-1^) in 298 K and 287 K, considering a pressure of 1 atm.

| Electron Acceptor | Reaction | ΔG_r_°  @ 25°C | ΔG_r_°  @ 14°C | ΔH_r_° |
| --- | --- | --- | --- | --- |
| Amorphous manganese | $\frac{1}{2}MnO_{2}+\frac{1}{3}{{NH}_{4}}^{+}+\frac{2}{3}H^{+}\to\frac{1}{2}Mn^{2+}+\frac{1}{6}N_{2}+H_{2}O$ | -86.80 | -87.18 | -97.21 |
| Magnetite | $\frac{1}{2}Fe_{3}O_{4}+\frac{1}{3}NH_{4}^{+}+\frac{8}{3}H^{+} \to$ $\frac{3}{2}Fe^{2+}+\frac{1}{6}N_{2}+2H_{2}O$ | -56.72 | -58.45 | -103.66 |
| Ferrihydrite | $Fe\left( OH \right)_{3}+\frac{1}{3}NH_{4}^{+}+\frac{5}{3}H^{+} \to$ $Fe^{2+}+\frac{1}{6}N_{2}+3H_{2}O$ | -65.66 | -66.24 | -81.14 |
| Amorphous iron | $FeOOH+\frac{1}{3}{{NH}_{4}}^{+}+\frac{5}{3}H^{+}\to$ $Fe^{2}+ \frac{1}{6}N_{2}+2H_{2}O$ | -62.98 | -63.11 | -66.31 |
| Lepidocrocite | $\gamma-FeOOH+\frac{1}{3}{{NH}_{4}}^{+}+\frac{5}{3}H^{+}\to$ $Fe^{2+}+\frac{1}{6}N_{2}+2H_{2}O$ | -42.28 | -43.17 | -66.31 |
| Goethite | $\alpha-FeOOH+\frac{1}{3}{{NH}_{4}}^{+}+\frac{5}{3}H^{+}\to$ $Fe^{2+}+\frac{1}{6}N_{2}+2H_{2}O$ | -36.28 | -37.12 | -58.91 |
| Hematite | $\frac{1}{2}Fe_{2}O_{3}+\frac{1}{3}{NH}_{4}^{+}+\frac{5}{3} H^{+} \to$ $Fe^{2+}+ \frac{1}{6}N_{2}+\frac{3}{2}H_{2}O$ | -35.29 | -36.33 | -73.93 |

**Table S12:** Balanced ammonium (NH_4_^+^) oxidation to nitrogen (N_2_) reactions standard Gibbs free energies (kJ mol^-1^) and enthalpies (kJ mol^-1^) in 298 K and 287 K, considering a pressure of 1 atm.

| Electron Acceptor | Reaction | ΔG_r_°  @ 25°C | ΔG_r_°  @ 14°C | ΔH_r_° |
| --- | --- | --- | --- | --- |
| Amorphous manganese | $\frac{1}{2}MnO_{2}+\frac{1}{6}N{H_{4}}^{+}+\frac{2}{3}H^{+} \to$ $\frac{1}{2}Mn^{2+}+\frac{1}{6}{NO}_{2}^{-}+{\frac{2}{3}H}_{2}O$ | -32.57 | -32.71 | -36.30 |
| Magnetite | $\frac{1}{2}Fe_{3}O_{4}+\frac{1}{6}NH_{4}^{+}+\frac{8}{3}H^{+} \to$ $\frac{3}{2}Fe^{2+}+\frac{1}{6}{NO}_{2}^{-}+\frac{5}{3}H_{2}O$ | 1.12 | -0.62 | -46.18 |
| Ferrihydrite | $Fe\left( OH \right)_{3}+\frac{1}{6}NH_{4}^{+}+\frac{5}{3}H^{+} \to$ $Fe^{2+}+\frac{1}{6}NO_{2}^{-}+\frac{8}{3}H_{2}O$ | -7.82 | -8.41 | -23.66 |
| Amorphous iron | $FeOOH+\frac{1}{6}{{NH}_{4}}^{+}+\frac{5}{3}H^{+}\to$ $Fe^{2+}+ \frac{1}{6}NO_{2}^{-}+\frac{5}{3}H_{2}O$ | -5.14 | -5.28 | -8.83 |
| Lepidocrocite | $\gamma-FeOOH+\frac{1}{6}{{NH}_{4}}^{+}+\frac{5}{3}H^{+}\to$ $Fe^{2+}+\frac{1}{6}NO_{2}^{-}+\frac{5}{3}H_{2}O$ | +15.56 | +14.66 | -8.83 |
| Goethite | $FeO\left( OH \right)+\frac{1}{6}{{NH}_{4}}^{+}+\frac{5}{3}H^{+}\to$ $Fe^{2+}+\frac{1}{6}NO_{2}^{-}+\frac{5}{3}H_{2}O$ | +21.56 | +20.71 | -1.43 |
| Hematite | $\frac{1}{2}Fe_{2}O_{3}+\frac{1}{6}{NH}_{4}^{+}+\frac{5}{3} H^{+} \to$ $Fe^{2+}+ \frac{1}{6}NO_{2}^{-}+\frac{7}{6}H_{2}O$ | +22.55 | +21.50 | -5.82 |

**Table S13:** Balanced ammonium (NH_4_^+^) oxidation to nitrite (NO_2_^-^) reactions, standard Gibbs free energies (kJ mol^-1^) and enthalpies (kJ mol^-1^) in 298 K and 287 K, considering a pressure of 1 atm.

| Electron Acceptor | Reaction | ΔG_r_°  @ 25°C | ΔG_r_°  @ 14°C | ΔH_r_° |
| --- | --- | --- | --- | --- |
| Amorphous manganese | $\frac{1}{2}MnO_{2}+\frac{1}{8}{{NH}_{4}}^{+}+\frac{3}{4}H^{+}\to\frac{1}{2}Mn^{2+}+\frac{1}{8}NO_{3}^{-}+\frac{5}{8}H_{2}O$ | -33.72 | -33.89 | -38.39 |
| Magnetite | $\frac{1}{2}Fe_{3}O_{4}+\frac{1}{8}NH_{4}^{+}+\frac{11}{4}H^{+} \to$ $\frac{3}{2}Fe^{2+}+\frac{1}{8}NO_{3}^{-}+\frac{13}{8}H_{2}O$ | -0.02 | -1.80 | -48.27 |
| Ferrihydrite | $Fe\left( OH \right)_{3}+\frac{1}{8}NH_{4}^{+}+\frac{7}{4}H^{+} \to$ $Fe^{2+}+\frac{1}{8}NO_{3}^{-}+\frac{21}{8}H_{2}O$ | -8.97 | -9.59 | -25.75 |
| Amorphous iron | $FeOOH+\frac{1}{8}{{NH}_{4}}^{+}+\frac{7}{4}H^{+}\to$ $Fe^{2+}+\frac{1}{8}NO_{3}^{-}+\frac{13}{8}H_{2}O$ | -6.29 | -6.46 | -10.92 |
| Lepidocrocite | $\gamma-FeOOH++\frac{1}{8}{{NH}_{4}}^{+}+\frac{7}{4}H^{+}\to Fe^{2+}+\frac{1}{8}NO_{3}^{-}+\frac{13}{8}H_{2}O$ | +14.41 | +13.48 | -10.92 |
| Goethite | $\alpha-FeOOH+\frac{1}{8}{{NH}_{4}}^{+}+\frac{7}{4}H^{+}\to$ $Fe^{2+}+\frac{1}{8}NO_{3}^{-}+\frac{13}{8}H_{2}O$ | +20.41 | +19.53 | -3.52 |
| Hematite | $\frac{1}{2}Fe_{2}O_{3}+\frac{1}{8}{NH}_{4}^{+}+\frac{7}{4} H^{+} \to$ $Fe^{2+}+ \frac{1}{8}NO_{3}^{-}+\frac{9}{8}H_{2}O$ | +21.40 | +20.32 | -7.91 |

**Table S14:** Balanced ammonium (NH_4_^+^) oxidation to nitrate (NO_3_^-^) reactions standard Gibbs free energies (kJ mol^-1^) and enthalpies (kJ mol^-1^) in 298 K and 287 K, considering a pressure of 1 atm.

| Electron Acceptor | Reaction | ΔG_r_°  @ 25°C | ΔG_r_°  @ 14°C | ΔH_r_° |
| --- | --- | --- | --- | --- |
| Amorphous manganese | $\frac{1}{2}MnO_{2}+\frac{1}{2}FeS+2H^{+} \to$ $\frac{1}{2}Mn^{2+}+\frac{1}{2}Fe^{2+}+\frac{1}{2}S^{0}+H_{2}O$ | -107.90 | -108.74 | -130.78 |
| Magnetite | $\frac{1}{2}Fe_{3}O_{4}+ \frac{1}{2}FeS+4H^{+} \to2Fe^{2+}+ \frac{1}{2}S^{0}+2H_{2}O$ | -98.93 | -102.20 | -187.55 |
| Ferrihydrite | $Fe\left( OH \right)_{3}+ \frac{1}{2}FeS+3H^{+} \to$ $\frac{3}{2}Fe^{2+}+ \frac{1}{2}S^{0}+3H_{2}O$ | -83.15 | -84.44 | -118.14 |
| Amorphous iron | $FeOOH+\frac{1}{2}FeS+3H^{+}\to$ $\frac{3}{2}Fe^{2+}+\frac{1}{2}S^{0}+2H_{2}O$ | -80.46 | -81.31 | -103.31 |
| Lepidocrocite | $\gamma-FeOOH+\frac{1}{2}FeS+3H^{+}\to$ $\frac{3}{2}Fe^{2+}+\frac{1}{2}S^{0}+2H_{2}O$ | -74.20 | -76.65 | -140.66 |
| Goethite | $\alpha-FeOOH+\frac{1}{2}FeS+3H^{+}\to$ $\frac{3}{2}Fe^{2+}+\frac{1}{2}S^{0}+2H_{2}O$ | -53.77 | -55.32 | -95.91 |
| Hematite | $\frac{1}{2}Fe_{2}O_{3}+ \frac{1}{2}FeS+3H^{+} \to$ $\frac{3}{2}Fe^{2+}+ \frac{1}{2}S^{0}+\frac{3}{2}H_{2}O$ | -52.78 | -54.53 | -100.30 |

**Table S15:** Balanced iron sulfide (FeS) oxidation reactions, standard Gibbs free energies (kJ mol^-1^) and enthalpies (kJ mol^-1^) in 298 K and 287 K, considering a pressure of 1 atm.

6. **Gibbs free energies of net redox reactions in natural environments**

| **Environment** | **EA** | **CH_3_COO^-^** | **H_2_** | **CH_4_** | **NH_4_^+^→ N2** | **FeS** | **NH_4_^+^→ NO_2_^-^** | **NH_4_^+^→ NO_3_^-^** |
| --- | --- | --- | --- | --- | --- | --- | --- | --- |
| LK | MnO_2_ (amorphous) | -97.0 | -90.8 | -78.8 | -72.9 | -58.6 | -24.1 | -20.9 |
|  | Fe(OH)_3_ (ferrihydrite) | -44.8 | -40.2 | -28.2 | -25.7 | -8.1 | +26.5 | +29.6 |
|  | FeOOH (amorphous) | -41.7 | -37.1 | -25.1 | -22.6 | -5.0 | +29.6 | +32.7 |
|  | γ-FeOOH (lepidocrocite) | -21.8 | -17.2 | -5.2 | -2.7 | +15.0 | +48.4 | +52.7 |
|  | α-FeOOH (goethite) | -15.8 | -11.2 | +0.8 | +3.3 | +20.9 | +55.5 | +58.6 |
|  | Fe_2_O_3_ (hematite) | -14.9 | -10.3 | +1.7 | +4.2 | +21.8 | +56.4 | +59.5 |
|  | Fe_3_O_4_ (magnetite) | -11.6 | -7.0 | +5.0 | +7.5 | +25.2 | +59.7 | +62.8 |
| PC-3 | MnO_2_ (amorphous) | -92.9 | -82.1 | -74.2 | -71.0 | -54.3 | -21.6 | -18.1 |
|  | Fe(OH)_3_ (ferrihydrite) | -45.1 | -36.9 | -28.9 | -29.2 | -9.1 | +23.7 | +27.2 |
|  | FeOOH (amorphous) | -41.9 | -33.7 | -25.7 | -26.0 | -5.9 | +26.9 | +30.4 |
|  | γ-FeOOH (lepidocrocite) | -22.0 | -13.8 | -5.8 | -6.0 | +14.1 | +45.6 | +50.3 |
|  | α-FeOOH (goethite) | -16.0 | -7.8 | +0.1 | -0.1 | +20.0 | +52.7 | +56.3 |
|  | Fe_2_O_3_ (hematite) | -15.1 | -6.9 | +1.1 | +0.8 | +20.9 | +53.7 | +57.2 |
|  | Fe_3_O_4_ (magnetite) | -12.1 | -3.9 | +4.1 | +3.9 | +24.0 | +56.7 | +60.2 |
| SG-1 | MnO_2_ (amorphous) | -92.9 | -78.6 | -74.6 | -71.0 | -54.3 | -21.6 | -18.1 |
|  | Fe(OH)_3_ (ferrihydrite) | -45.1 | -33.3 | -29.2 | -29.2 | -9.1 | +23.7 | +27.2 |
|  | FeOOH (amorphous) | -41.9 | -30.1 | -26.1 | -26.0 | -5.9 | +26.9 | +30.4 |
|  | γ-FeOOH (lepidocrocite) | -22.0 | -10.2 | -6.2 | -6.0 | +14.1 | +45.6 | +50.3 |
|  | α-FeOOH (goethite) | -16.0 | -4.3 | -0.2 | -0.1 | +20.0 | +52.7 | +56.3 |
|  | Fe_2_O_3_ (hematite) | -15.1 | -3.3 | +0.7 | +0.8 | +20.9 | +53.7 | +57.2 |
|  | Fe_3_O_4_ (magnetite) | -12.1 | -0.3 | +3.7 | +3.9 | +24.0 | +56.7 | +60.2 |

**Table S16:** Gibbs free energies (ΔG_r_) (kJ (mol e^-^)^-1^) of net redox reactions in LK, PC-3 and SG-1, considering a temperature of 287 K and a pressure of 1 atm, normalized to the transfer of one electron. Positive ΔG_r_ values, for which the reactions are not thermodynamically favorable, are shown in red.

7. **Kinetic parameters**

| **Specie** | **K_s_  (mol L^-1^)** | **Reference** | **V_max_  (mol cell^-1^ day^-1^)** | **Reference** |
| --- | --- | --- | --- | --- |
| Fe(OH)_3_ (ferrihydrite) | 7.00∙10^-4^ – 3.00∙10^-3^ | Bonneville et al. (2004) | 1.94∙10^-15^ | Bonneville et al. (2004) |
| Fe_2_O_3_ (hematite) | 6.00∙10^-4^– 1.80∙10^-3^ | Bonneville et al. (2004) | 5.76∙10^-16^ | Bonneville et al. (2004) |
| Fe_3_O_4_ (magnetite) |  | NOT AVAILABLE |  | NOT AVAILABLE |
| α-FeOOH (goethite) | 1.45∙10^-2^– 2.79∙10^-2^ | Bonneville et al. (2004) | 4.80∙10^-16^ | Bonneville et al. (2004) |
| FeOOH (amorphous) | 6.00∙10^-4^– 1.00∙10^-3^ | Bonneville et al. (2004) | 1.56∙10^-15^ | Bonneville et al. (2004) |
| γ-FeOOH (lepidocrocite) | 1.45∙10^-2^– 2.79∙10^-2^ | Bonneville et al. (2004) | 4.80∙10^-16^ | Bonneville et al. (2004) |
| MnO_2_ (amorphous) | 8.88∙10^-3^ | Lovley and Phillips (1988b) | 1.30∙10^-13^ | Lovley and Phillips (1988b) |
| H_2_ | 5.80∙10^-5^ | Lovley et al. (1982) |  |  |
| CH_3_COO^-^ | 6.00∙10^-4^– 1.00∙10^-3^ | Oude Elferink et al. (1998) |  |  |
| CH_4_ | 1.50∙10^-3^ | Dale et al. (2006) |  |  |
| NH_4_^+^ | 1.50∙10^-3^ | Chen et al. (2011) |  |  |
| FeS |  | NOT AVAILABLE |  |  |
| H^+^ | 0.00 |  |  |  |
| Mn_2_^+^ |  |  |  |  |

**Table S17:** Kinetic parameters for microbial respiration. Estimated values based on the mentioned references are marked in red.

8. **Model’s input for concentrations and activities** **in the methanic zone**

| **Specie** |  | **Concentration (mM)** | **Activity (M)** | **Concentration range (mM)** |
| --- | --- | --- | --- | --- |
| pH |  |  | 8.1 | 8.0-8.1 (June 2015) |
| DIC |  | 30 |  | 10-20 (Sep 2015) 10 – 80 (Nov 2017) |
| HCO_3_^-^ |  |  | 1.5⸱10^-2^ |  |
| Ca^2+^ |  | 4.0 |  | 3.6-3.8 (Sep 2015) 4.1-4.2 (June 2015) |
| Cl^-^ |  | 620 |  | 615-620 (Sivan et al., 2004) |
| K^+^ |  | 11.3 |  | 11.5-12 (Sep 2015) 11.3-11.6 (June 2015) |
| Mg^2+^ |  | 44 |  | 49.0-51.8 (Sep 2015) 41-44 (June 2015) |
| Na^+^ |  | 560 |  | 511.1-528.5 (Sep 2015) 478-561 (June 2015) |
| Mn^2+^ |  | 0.004 | 2⸱10^-6^ | 0.004-0.020 |
| Fe^2+^ |  | 0.005 | 1∙10^-7^ | 0.001-0.017 (June 2015) 0.001-0.035 (Sep 2017) |
| H_2_ |  | 0.0001 | 6∙10^-8^ | 0.0001 (Sep 2017) |
|  |  | 0.004 | 2.6∙10^-6^ | 0.002-0.006 |
| CH_4_ |  | 1.5 | 1.5∙10^-3^ | 1.1-2.0 (June 2015) 1-3 (Nov 2017) |
| NH_4_^+^ |  | 5 | 3∙10^-3^ | 4-6 (Dec 2022) |
| NO_2_^-^ |  | 0.002 | 7∙10^-7^ | 0.0018-0.0021 (Dec 2022) |
| NO_3_^-^ |  | 0.002 | 1∙10^-6^ | 0.002 (Dec 2022) |
| Oxides 1 |  | 57 | 1 | 12-203 (Dec 2022) |
| Fe(OH)_3_ (Ferrihydrite) |  | 57 | 1 |  |
| FeOOH (Amorphous) |  | 57 | 1 |  |
| γ-FeOOH (Lepidocrocite) |  | 57 |  |  |
| Oxides 2 |  | 387 | 1 | 52-387 (Dec 2022) |
| Fe_2_O_3_ (Hematite) |  | 387 | 1 |  |
| α-FeOOH (Goethite) |  | 387 | 1 |  |
| Fe_3_O_4_ (Magnetite) |  | 7 | 1 | 7-86 (Dec 2022) |

**Table S18:** Model’s input for concentrations and activities in the methanic zone of the MedS, at a depth of 400 cm below sediment-water interface at station SG-1 from September 2015, June 2015, January 2017, September 2017, November 2017 and December 2022 (after Vigderovich et al. (2019); Wurgaft et al. (2019); Yorshansky et al. (2022) and unpublished data). Acetate concentrations were measured by Zhuang et al. (2018) in station GeoB17306 in the Western Mediterranean. Amorphous manganese oxide concentrations were measured by Gershon and Boyko (unpublished results) in station A in LK (Table S20). Measured Concentration ranges are mentioned. Selected concentrations for the model are average concentrations. Activities were calculated using the PHREEQC program. As ferrihydrite, amorphous iron oxyhydroxide and lepidocrocite extract as the same phase (oxi.1), their concentrations were assumed to equal the total concertation of this phase.

| **Specie** | **Concentration (mM)** | **Activity (M)** | **Concentration Range (mM)** |
| --- | --- | --- | --- |
| H_2_ | 0.003 | 1∙10^-6^ | 0.002-0.006 (January 2015) |
| CH_4_ | 0.4 | 4∙10^-4^ | 0.3-0.7 (August 2013) 0.1-0.4 (January 2015) |
| Fe^2+^ | 0.01 |  | 0.001-0.062 (August 2013) 0.001 (January 2015) |
| Oxides 1 | 233 |  | 58 (January 2015) |
| Fe(OH)_3_ (Ferrihydrite) | 233 | 1 |  |
| FeOOH (Amorphous) | 233 | 1 |  |
| γ-FeOOH (Lepidocrocite) | 233 | 1 |  |
| Oxides 2 | 151 |  | 45 (January 2015) |
| Fe_2_O_3_ (Hematite) | 151 | 1 |  |
| α-FeOOH (Goethite) | 151 | 1 |  |
| Fe_3_O_4_ (Magnetite) | 64 | 1 | (January 2015) |

**Table S19:** Model’s input for concentrations and activities in the methanic zone of the MedS, at a depth of 400 cm below sediment-water interface at station PC-3 from August 2013 and January 2015 (after Vigderovich et al. (2019)). Iron oxides were extracted as amorphous/poorly poorly crystalline iron oxides (Oxides 1), crystalline iron oxides (Oxides 2) and magnetite. Amorphous manganese oxide concentrations were measured by Gershon and Boyko (unpublished results) in station A in LK (Table S20). Concentrations of other relevant species are from SG-1 (Table S18). Measured Concentration ranges are mentioned. Selected concentrations for the model are average concentrations. Activities were calculated using the PHREEQC program. As ferrihydrite, amorphous iron oxyhydroxide and lepidocrocite extract as the same phase (oxi.1), their concentrations were assumed to equal the total concertation of this phase.

| **Specie** | **Concentration (mM)** | **Activity (M)** | **Concentration Range (mM)** |
| --- | --- | --- | --- |
| pH |  | 7.2 |  |
| DIC | 10 |  | 9-10 (March 2020) 11-12 (Sep 2020) |
| HCO_3_^-^ |  | 5.7∙10^-3^ |  |
| Ca^2+^ | 3 |  | 2.9-3.1 (March 2020) |
| K^+^ | 0.2 |  | 0.2 (March 2020) |
| Mg^2+^ | 2 |  | 2 (March 2020) |
| Na^+^ | 7 |  | 7.4-7.9 (March 2020) |
| Mn^2+^ | 0.03 | 2∙10^-5^ | 0.02-0.03 (March 2020) |
| Fe^2+^ | 0.1 | 3.8∙10^-5^ | 0.08-0.1 (March 2020) 0.1-0.2 (Sep 2020) |
| H_2_ | 0.1 | 3.5∙10^-5^ | 0.07-0.33 (June 2022) |
| CH_3_COO^-^ | 0.2 | 2.6∙10^-5^ | 0.03-0.50 (March, 2020) |
| CH_4_ | 3 | 3∙10^-3^ | 3 (March 2020) 2-3 (Sep 2020) |
| NH_4_^+^ | 2 | 1.7∙10^-3^ | 1-2 (March 2020) |
| NO_2_^-^ | 0.0003 | 2.6∙10^-7^ | 0.0001-0.0005 (March 2020) |
| NO_3_^-^ | 0.0008 | 7∙10^-7^ | 0.0002-0.0015 (March 2020) |
| Oxides 1 | 27 |  | (2015) |
| Fe(OH)_3_ (Ferrihydrite) | 27 | 1 |  |
| FeOOH (Amorphous) | 27 | 1 |  |
| γ-FeOOH (Lepidocrocite) | 27 | 1 |  |
| Oxides 2 | 34 |  | (2015) |
| Fe_2_O_3_ (Hematite) | 34 | 1 |  |
| α-FeOOH (Goethite) | 34 | 1 |  |
| Fe3O4 (Magnetite) | 6 | 1 | (2015) |
| MnO_2_ (Amorphous) | 2 | 1 | (2019) |

**Table S20:** Model’s input for concentrations and activities of relevant aqueous substances in the methanic zone of LK, at a depth of 22 cm below sediment-water interface from September 2007, July 2008, August 2018 and October 2019 (Adler et al. (2011), Elul et al. (2020) and unpublished data). Iron oxides were extracted as amorphous/poorly poorly crystalline iron oxides (Oxides 1), crystalline iron oxides (Oxides 2) and magnetite. Measured Concentration ranges are mentioned. Activities were calculated using the PHREEQC program. As ferrihydrite, amorphous iron oxyhydroxide and lepidocrocite extract as the same phase (oxi.1), their concentrations were assumed to equal the total concertation of this phase.

9.
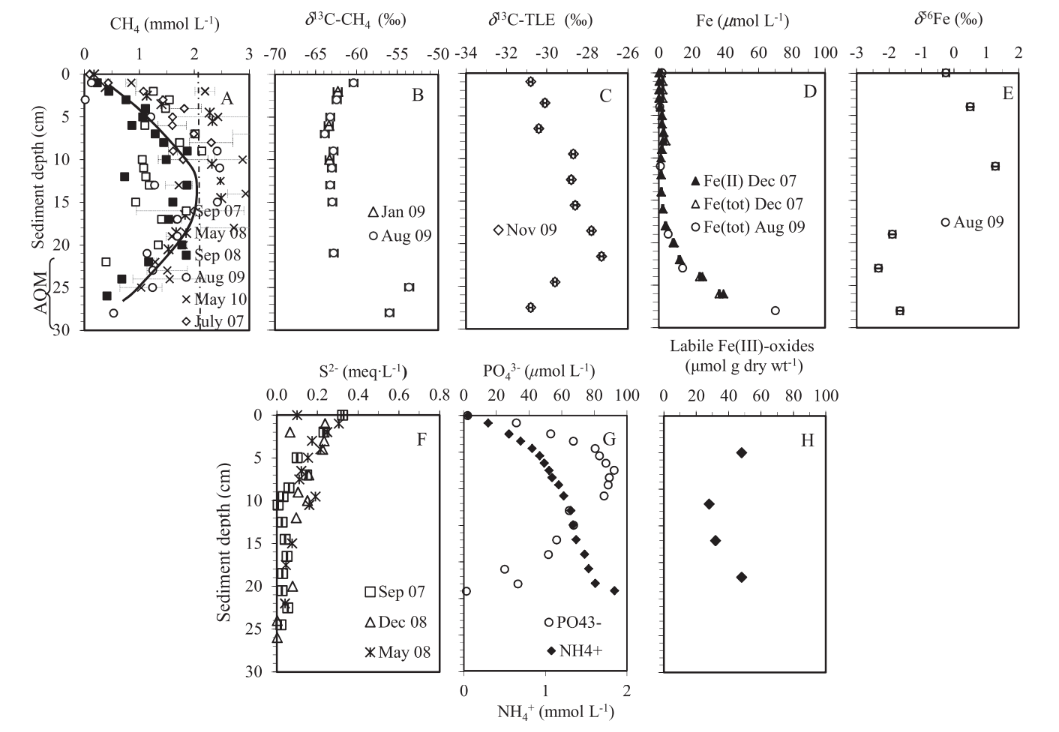
**Geochemical profiles**

**Figure S3:** Profiles in LK sediments. (A) Dissolved methane in pore water (error bar is marked when duplicates were measured). (B) δ^13^C of dissolved methane in pore water. (C) δ^13^C of TLE from the sediment. (D) Typical dissolved Fe(II) and Fe(total (tot)) in the porewater. (E) d56Fe in pore water. (F) Sulfide in pore water. (G) Phosphate and ammonium in pore water. (H) Highly accessible Fe(III)-oxides in LK sediments extracted by diluted ascorbic acid. The depth range of the sampling is 0.5–2 cm, and the error bar is smaller thanthe symbol, unless marked (after Sivan et al., 2011). Acetate concentrations of 0.03-0.50 mM are from Elul et al. (2020) and H_2_ concentrations of 0.07-0.33 mM are from unpublished data (Table S20).

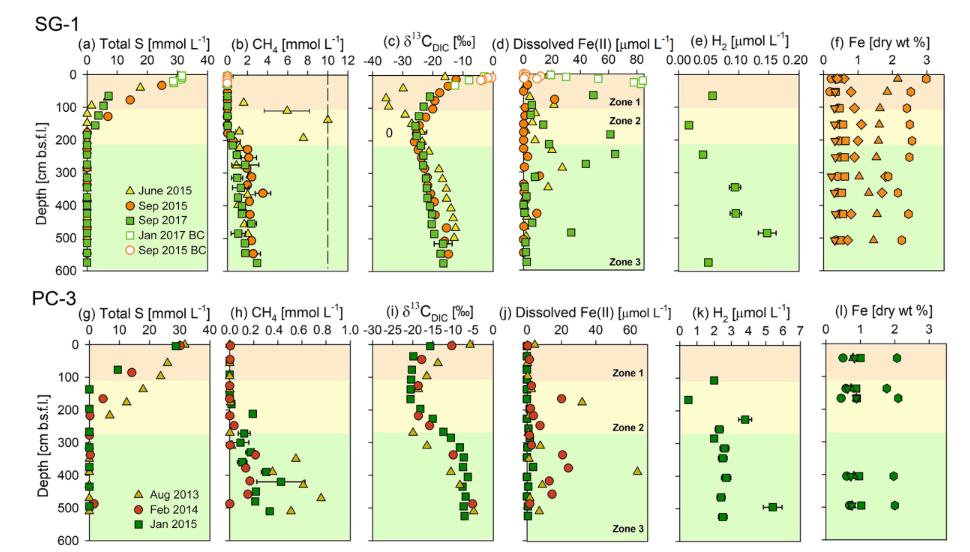

**Figure S4:** Geochemical porewater profiles of total S, CH_4_, δ^13^C_DIC_, dissolved Fe(II), H_2_ and extractable Fe fractions from sediment cores collected at the two stations: SG-1 (a–f) and PC-3 (g–l) in the SE Mediterranean. The profiles are divided roughly into three zones according to the dominant processes: upper microbial iron and sulfate reduction, sulfate-methane transition zone (SMTZ), and the methanic zone at the deep part. The dashed line in the CH_4_ graph at SG-1 station represents the CH_4_ saturation value in the porewater (Sela-Adler et al., 2015). The following extractable Fe fraction profiles of stations SG-1 (f) and PC-3 (l) are from the September and January 2015 cruise (respectively): Fe_carb_ (circle), Fe_ox1_ (square) Fe_ox2_ (triangle), Fe_mag_ (inverted triangle), Fe_py_ (diamond) (Wurgaft et al., 2019) and total reactive iron (hexagon). The error bars for CH_4_ are presented where duplicate sediment samples were collected. The error bars for Fe(II), δ^13^C_DIC_ and H_2_ are presented where measurements from the same sample were repeated at least twice. The analytical errors were too small to be displayed (after Vigderovich et al., 2019).

10. **Microbiological profiles**

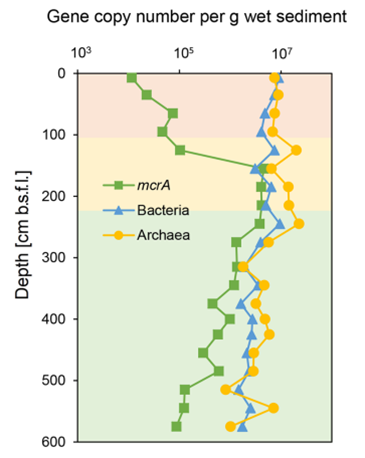

**Figure S5:** Sedimentary depth profiles of bacterial and archaeal 16S rRNA and *mrcA* functional genes of station SG-1 from January 2017 (after Vigderovich et al. (2019)).

11. **Relative functional group sizes of iron reducers** **and total iron reduction rates in Lake Kinneret**

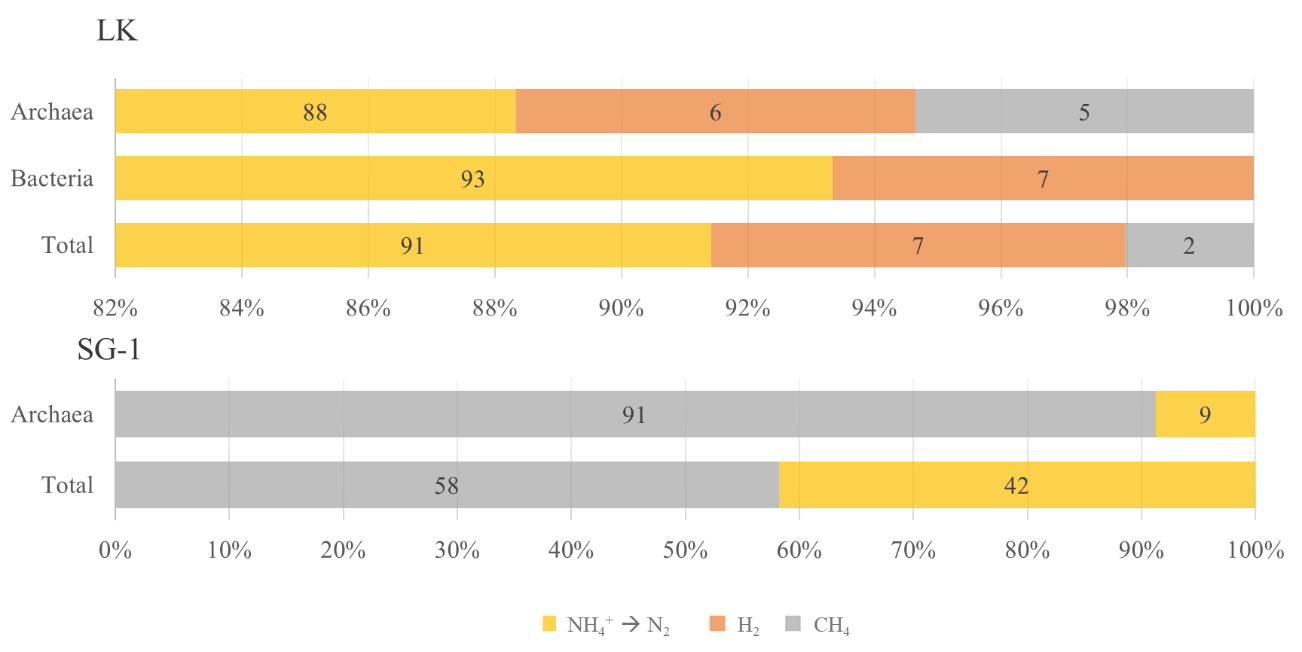

**Figure S6:** Relative functional group sizes of iron reducers only in the methanogenic zones of LK an SG-1, divided by electron donor, according to the nominal model and assuming direct relation to the calculated growth rates. Iron sulfide (FeS) oxidizing functional groups are not included.

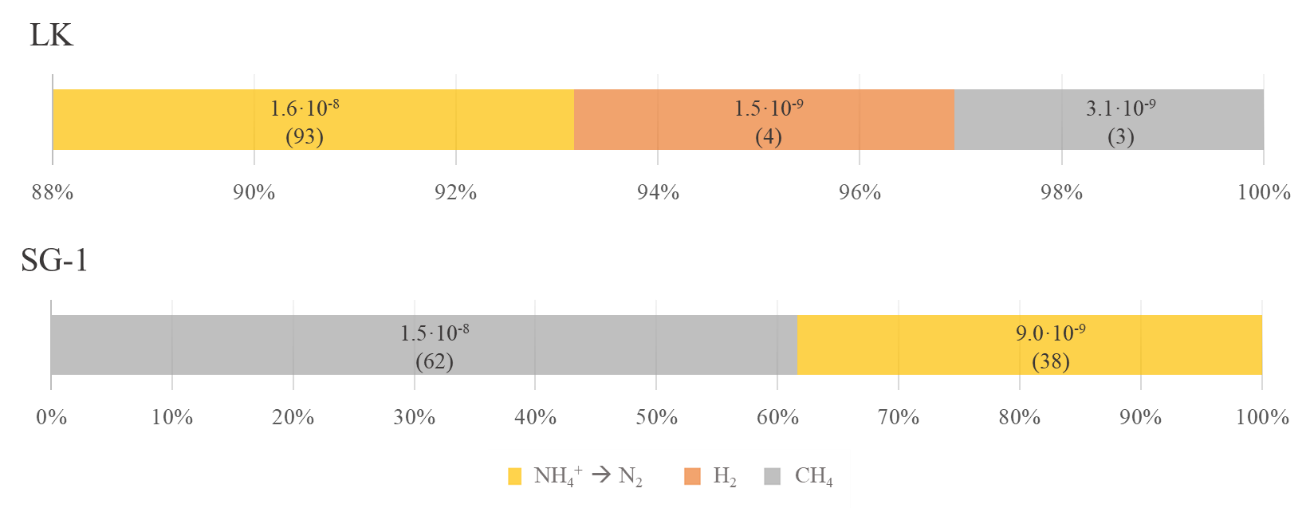
 **Figure S7:** Iron reduction rates (mol oxide cm^-3^ day^-1^) in the methanogenic zones of LK and SG-1, divided by electron donor, according to the nominal model and assuming direct relation to the calculated growth rates. In parentheses are the relative rates of each metabolic pathway in percent (%). Iron sulfide (FeS) oxidizing functional groups are not included.

12. **Sensitivity analysis**

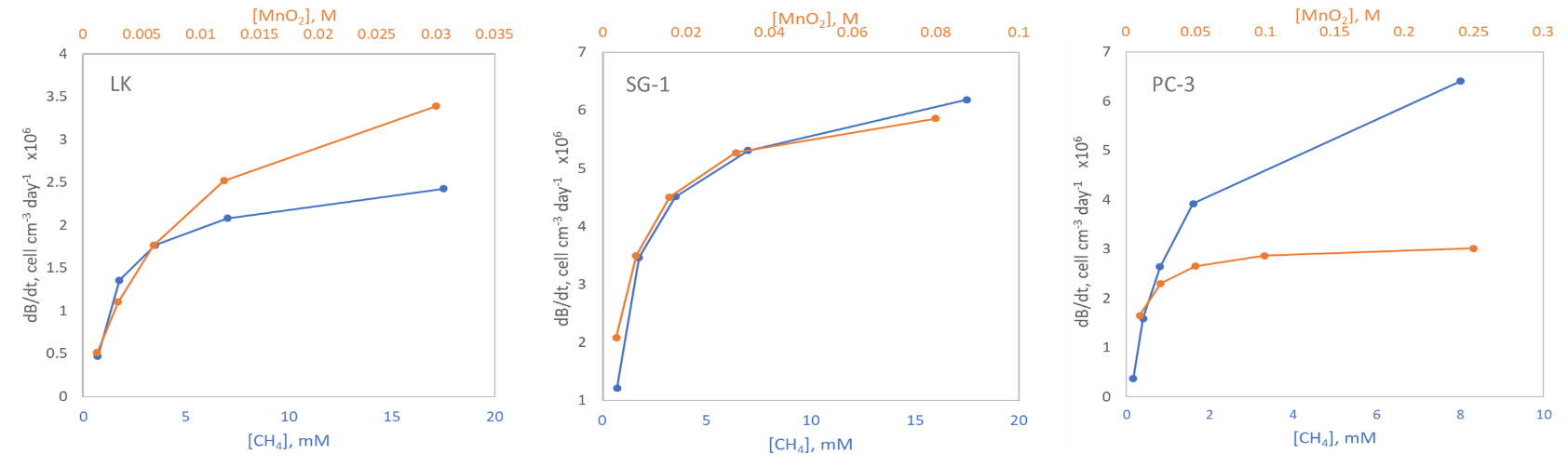
 **Figure S8:** Biomass growth rates for methane oxidation in LK, SG-1 and PC-3, versus methane concentrations of 0.2, 0.5, 1, 2 and 5 times the nominal value. Modeled bioreactions that do not appear in this figure generate negative biomass growth rates in all scenarios.

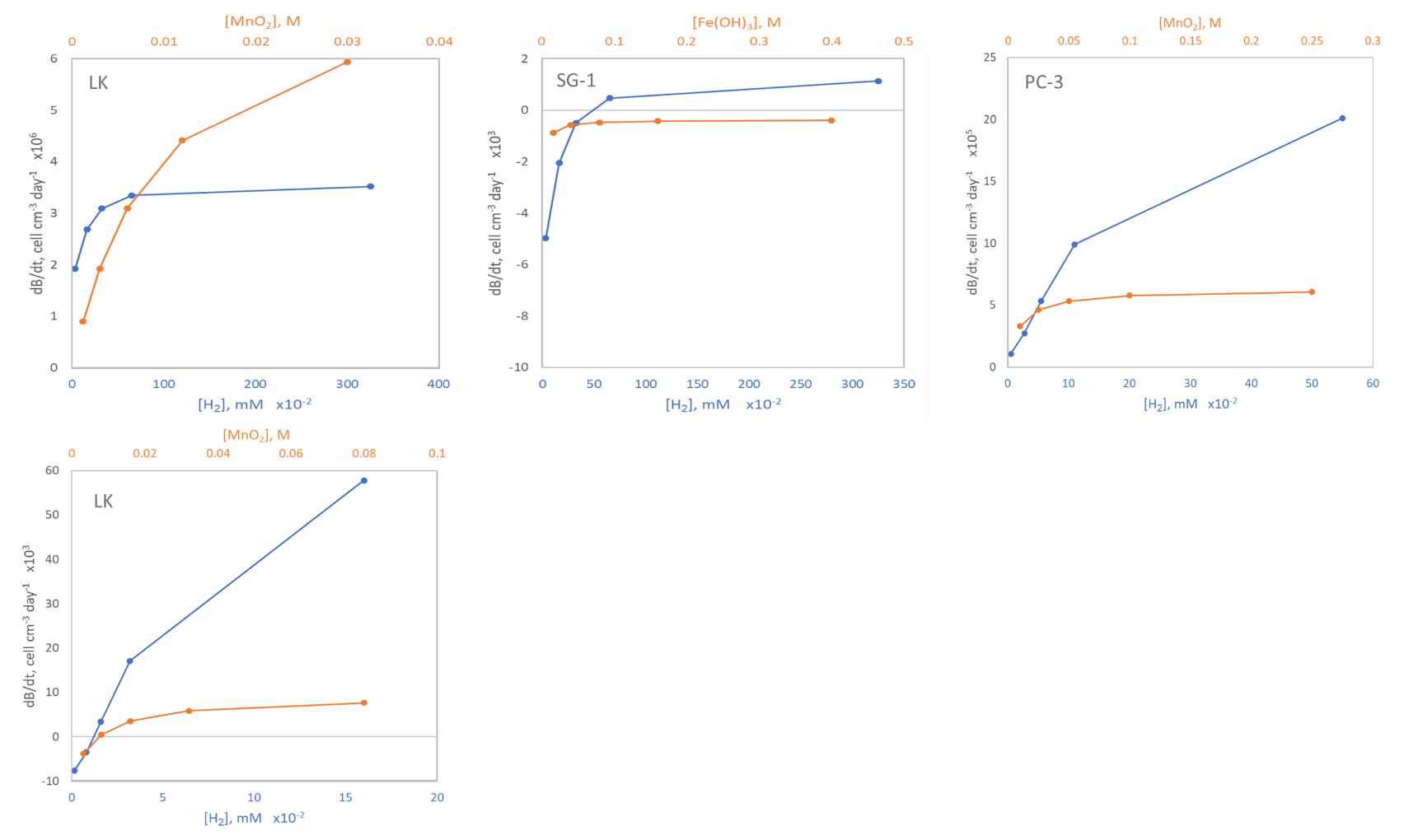

**Figure S9:** Biomass growth rates for hydrogen oxidation in LK, SG-1 and PC-3, versus hydrogen concentrations of 0.2, 0.5, 1, 2 and 5 times the nominal value. Modeled bioreactions that do not appear in this figure generate negative biomass growth rates in all scenarios.

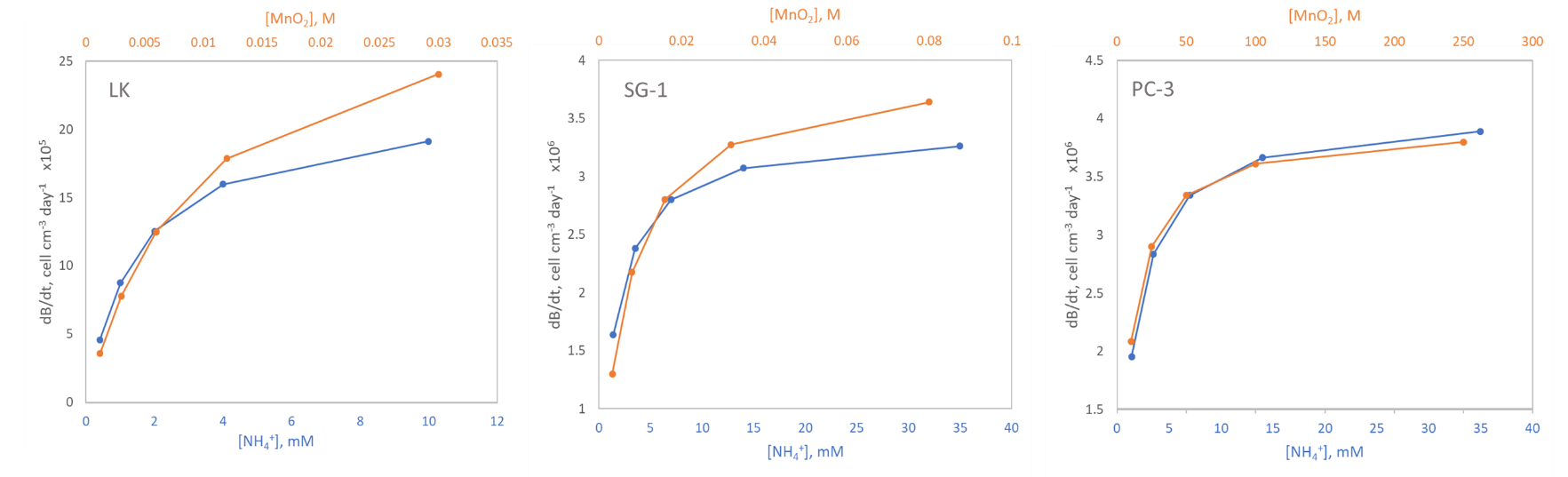
 **Figure S10:** Biomass growth rates for ammonium oxidation to molecular nitrogen in LK and the MedS (SG-1 and PC-3), versus ammonium concentrations of 0.2, 0.5, 1, 2 and 5 times the nominal value. 1. Modeled bioreactions that do not appear in this figure generate negative biomass growth rates in all scenarios.

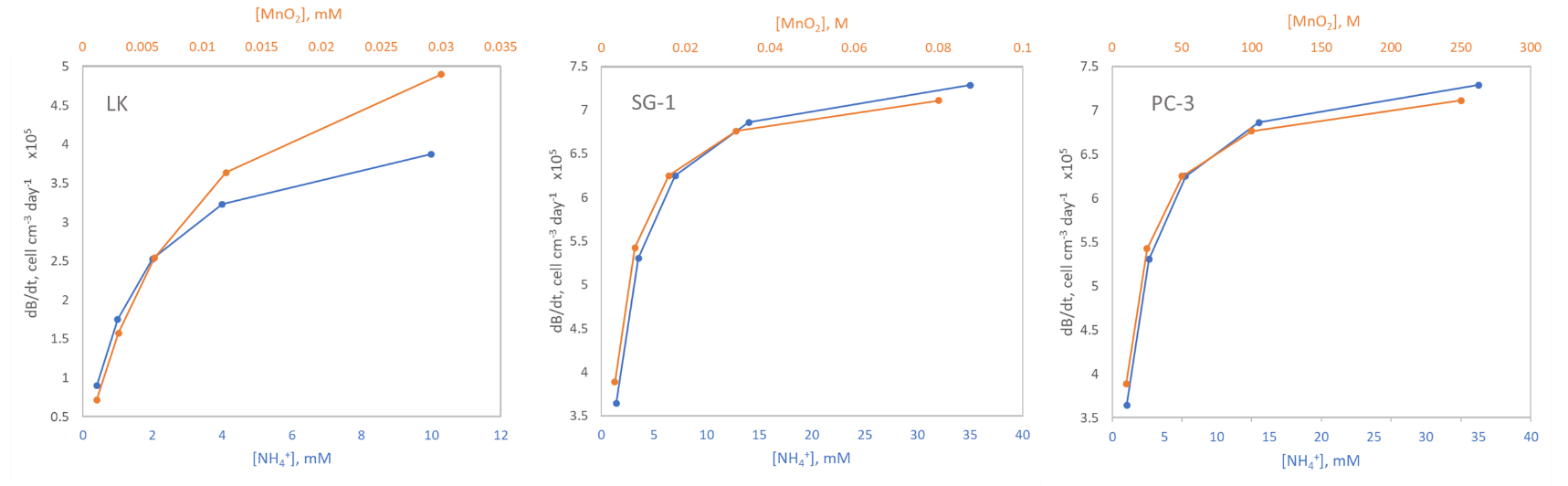
 **Figure S11:** Biomass growth rates for ammonium oxidation to nitrite in LK and the MedS (SG-1 and PC-3), versus ammonium concentrations of 0.2, 0.5, 1, 2 and 5 times the nominal value. 1. Modeled bioreactions that do not appear in this figure generate negative biomass growth rates in all scenarios.

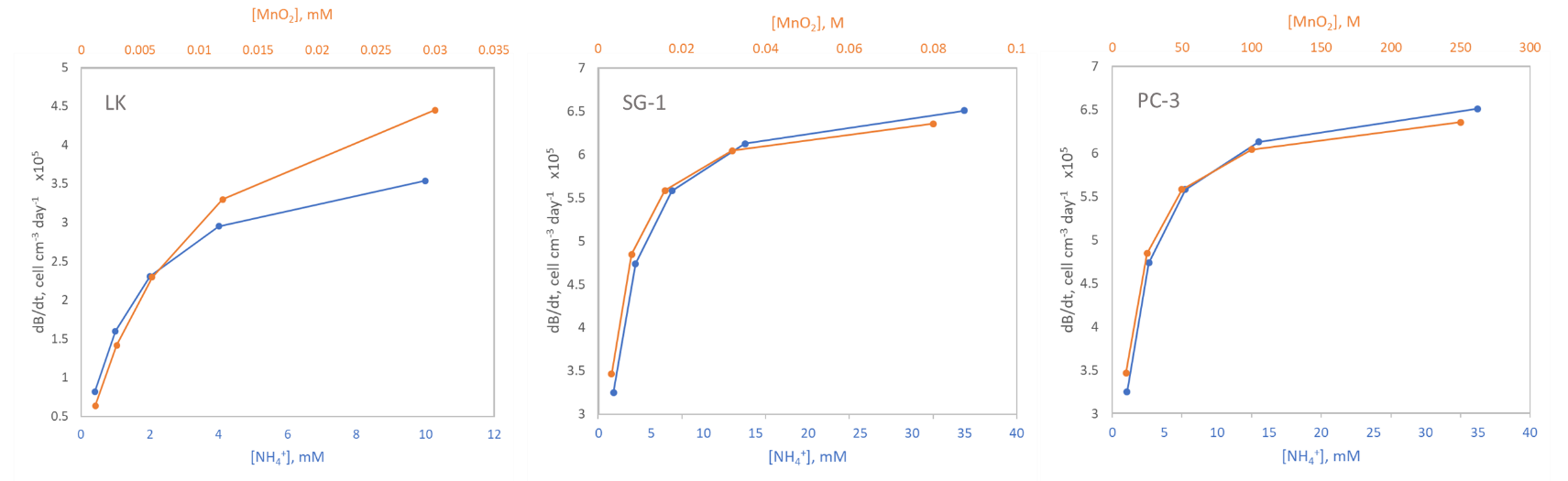
**Figure S12:** Biomass growth rates for ammonium oxidation to nitrate in LK and the MedS (SG-1 and PC-3), versus ammonium concentrations of 0.2, 0.5, 1, 2 and 5 times the nominal value. 1. Modeled bioreactions that do not appear in this figure generate negative biomass growth rates in all scenarios.

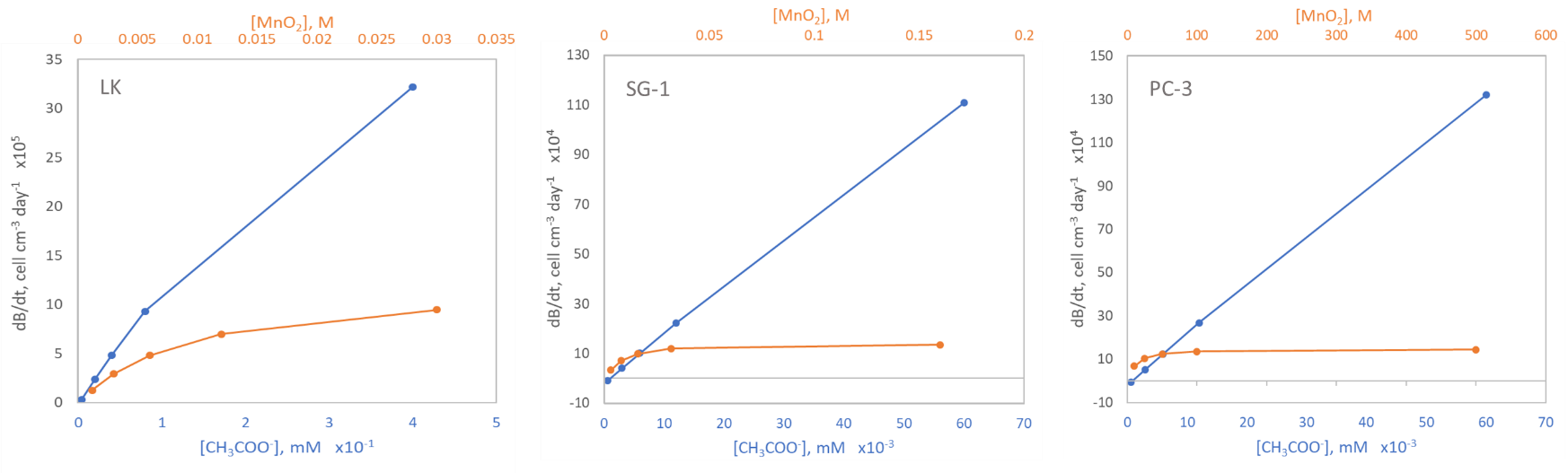

**Figure S13:** Biomass growth rates for acetate oxidation in LK and the MedS (SG-1 and PC-3), versus ammonium concentrations of 0.2, 0.5, 1, 2 and 5 times the nominal value. Modeled bioreactions that do not appear in this figure generate negative biomass growth rates in all scenarios.

| **Electron donor** | **Electron acceptor** | **Product** | **dB/dt for 2∙K_s_^ED^** | **Change in dB/dt (%)** | **dB/dt for 0.5∙K_s_^ED^** | **Change in dB/dt (%)** |
| --- | --- | --- | --- | --- | --- | --- |
| CH_4_ | MnO_2_ | HCO_3_^-^ | 2.70∙10^6^ | -2.69∙10 | 4.52∙10^6^ | 2.25∙10 |
| CH_4_ | Fe(OH)_3_ | HCO_3_^-^ | -1.79∙10^3^ |  | 3.53∙10^3^ | 2.19∙10^2^ |
| CH_4_ | FeOOH | HCO_3_^-^ | -4.03∙10^3^ |  | -2.88∙10^2^ |  |
| H_2_ | MnO_2_ | H_2_O | 3.25∙10^6^ | -3.13∙10 | 6.11∙10^6^ | 2.94∙10 |
| H_2_ | Fe(OH)_3_ | H_2_O | -6.66∙10^2^ |  | 9.62∙10^3^ | 1.07∙10^2^ |
| H_2_ | FeOOH | H_2_O | -3.67∙10^3^ |  | 3.86∙10^3^ | 1.71∙10^3^ |
| CH_3_COO^-^ | MnO_2_ | HCO_3_^-^ | 4.67∙10^5^ | -5.00∙10 | 1.80∙10^6^ | 9.30∙10 |
| CH_3_COO^-^ | Fe(OH)_3_ | HCO_3_^-^ | -2.42∙10^4^ |  | -1.92∙10^4^ |  |
| CH_3_COO^-^ | FeOOH | HCO_3_^-^ | -2.44∙10^4^ |  | -2.07∙10^4^ |  |
| NH_4_^+^ | MnO_2_ | NO_3_^-^ | 3.08∙10^5^ | -3.37∙10 | 6.21∙10^5^ | 3.37∙10 |
| NH_4_^+^ | MnO_2_ | NO_2_^-^ | 1.74∙10^6^ | -3.34∙10 | 3.48∙10^6^ | 3.34∙10 |
| NH_4_^+^ | MnO_2_ | N_2_ | -4.64∙10^3^ |  | 5.73∙10^2^ | -1.28∙10^2^ |
| NH_4_^+^ | Fe(OH)_3_ | N_2_ | -6.14∙10^3^ |  | -2.56∙10^3^ |  |
| NH_4_^+^ | FeOOH | N_2_ | 3.37∙10^5^ | -3.37∙10 | 6.79∙10^5^ | 3.37∙10 |

**Table S21:** Biomass growth rates (cell cm^-3^ day^-1^) in LK and change in biomass growth rates relatively to the nominal values, in scenarios where the half-saturation constants of the electron donors (*K_s_^ED^*) are 2 times the nominal values (blue-shaded columns) and 0.5 times the nominal values (orange-shaded columns). Negative biomass growth values are marked in red.

| **Electron donor** | **Electron acceptor** | **Product** | **dB/dt for 2∙K_s_^ED^** | **Change in dB/dt (%)** | **dB/dt for 0.5∙K_s_^ED^** | **Change in dB/dt (%)** |
| --- | --- | --- | --- | --- | --- | --- |
| CH_4_ | MnO_2_ | HCO_3_^-^ | 2.72∙10^6^ | -3.34∙10 | 5.45∙10^6^ | 3.34∙10 |
| CH_4_ | Fe(OH)_3_ | HCO_3_^-^ | -6.00∙10^3^ |  | 2.37∙10^3^ | -2.30∙10^2^ |
| CH_4_ | FeOOH | HCO_3_^-^ | -8.30∙10^3^ |  | -2.38∙10^3^ |  |
| H_2_ | MnO_2_ | H_2_O | -4.66∙10^3^ |  | 1.24E+04 | 1.11∙10^3^ |
| H_2_ | Fe(OH)_3_ | H_2_O | -1.10∙10^4^ |  | -1.09∙10^4^ |  |
| H_2_ | FeOOH | H_2_O | -1.09∙10^4^ |  | -1.08∙10^4^ |  |
| CH_3_COO^-^ | MnO_2_ | HCO_3_^-^ | 3.32∙10^4^ | -6.22∙10 | 1.96∙10^5^ | 1.23∙10^2^ |
| CH_3_COO^-^ | Fe(OH)_3_ | HCO_3_^-^ | -2.29∙10^4^ |  | -2.23∙10^4^ |  |
| CH_3_COO^-^ | FeOOH | HCO_3_^-^ | -2.28∙10^4^ |  | -2.23∙10^4^ |  |
| NH_4_^+^ | MnO_2_ | NO_3_ | 4.56∙10^5^ | -1.51∙10 | 5.89∙10^5^ | 9.70 |
| NH_4_^+^ | MnO_2_ | NO_2_^-^ | 5.51∙10^5^ | -1.50∙10 | 7.11∙10^5^ | 9.68 |
| NH_4_^+^ | MnO_2_ | N_2_ | 3.16∙10^6^ | -1.50∙10 | 4.08∙10^6^ | 9.63 |
| NH_4_^+^ | Fe(OH)_3_ | N_2_ | 1.88∙10^3^ | -4.98∙10 | 4.95∙10^3^ | 3.21∙10 |
| NH_4_^+^ | FeOOH | N_2_ | -1.18∙10^3^ |  | 9.61∙10^2^ | 6.99∙10^2^ |

**Table S22:** Biomass growth rates (cell cm^-3^ day^-1^) in SG-1 and change in biomass growth rates relatively to the nominal values, in scenarios where the half-saturation constants of the electron donors (*K_s_^ED^*) are 2 times the nominal values (blue-shaded columns) and 0.5 times the nominal values (orange-shaded columns). Negative biomass growth values are marked in red.

| **Electron donor** | **Electron acceptor** | **Product** | **dB/dt for 2∙K_s_^ED^** | **Change in dB/dt (%)** | **dB/dt for 0.5∙K_s_^ED^** | **Change in dB/dt (%)** |
| --- | --- | --- | --- | --- | --- | --- |
| CH_4_ | MnO_2_ | HCO_3_^-^ | 3.59∙10^6^ | -2.76∙10 | 3.75∙10^6^ | 1.44∙10 |
| CH_4_ | Fe(OH)_3_ | HCO_3_^-^ | 1.11∙10^3^ | -1.97∙10^2^ | 1.11∙10^3^ | 9.87∙10^3^ |
| CH_4_ | FeOOH | HCO_3_^-^ | -1.99∙10^3^ |  | -1.99∙10^3^ |  |
| H_2_ | MnO_2_ | H_2_O | 4.59∙10^6^ | -2.76∙10 | 4.79∙10^6^ | 1.44∙10 |
| H_2_ | Fe(OH)_3_ | H_2_O | 4.64∙10^3^ | -7.43∙10^3^ | 4.64∙10^3^ | 3.72∙10^3^ |
| H_2_ | FeOOH | H_2_O | 2.14∙10^2^ | -1.01∙10^-1^ | 2.14∙10^2^ | 5.06∙10^-2^ |
| CH_3_COO^-^ | MnO_2_ | HCO_3_^-^ | 9.08∙10^5^ | -2.82∙10 | 9.48∙10^5^ | 1.47∙10 |
| CH_3_COO^-^ | Fe(OH)_3_ | HCO_3_^-^ | -2.25∙10^4^ |  | -2.25∙10^4^ |  |
| CH_3_COO^-^ | FeOOH | HCO_3_^-^ | -2.31∙10^4^ |  | -2.31∙10^4^ |  |
| NH_4_^+^ | MnO_2_ | NO_3_^-^ | 4.51∙10^5^ | -2.78∙10 | 4.71∙10^5^ | 1.45∙10 |
| NH_4_^+^ | MnO_2_ | NO_2_^-^ | 4.94∙10^5^ | -2.78∙10 | 5.15∙10^5^ | 1.45∙10 |
| NH_4_^+^ | MnO_2_ | N_2_ | 2.54∙10^6^ | -2.76∙10 | 2.65∙10^6^ | 1.44∙10 |
| NH_4_^+^ | Fe(OH)_3_ | N_2_ | -2.04∙10^3^ |  | -2.03∙10^3^ |  |
| NH_4_^+^ | FeOOH | N_2_ | -4.35∙10^3^ |  | -4.35∙10^3^ |  |

**Table S23:** Biomass growth rates (cell cm^-3^ day^-1^) in LK and change in biomass growth rates relatively to the nominal values, in scenarios where the half-saturation constants of the electron acceptors (*K_s_^EA^*) are 2 times the nominal values (blue-shaded columns) and 0.5 times the nominal values (orange-shaded columns). Negative biomass growth values are marked in red.

| **Electron donor** | **Electron acceptor** | **Product** | **dB/dt for 2∙K_s_^ED^** | **Change in dB/dt (%)** | **dB/dt for 0.5∙K_s_^ED^** | **Change in dB/dt (%)** |
| --- | --- | --- | --- | --- | --- | --- |
| CH_4_ | MnO_2_ | HCO_3_^-^ | 3.97∙10^6^ | -2.76 | 4.15∙10^6^ | 1.44 |
| CH_4_ | Fe(OH)_3_ | HCO_3_^-^ | -1.97∙10^3^ |  | -1.74∙10^3^ |  |
| CH_4_ | FeOOH | HCO_3_^-^ | -5.43∙10^3^ |  | -5.29∙10^3^ |  |
| H_2_ | MnO_2_ | H_2_O | 7.06∙10^2^ | -3.07∙10 | 1.18∙10^3^ | 1.60∙10 |
| H_2_ | Fe(OH)_3_ | H_2_O | -1.09∙10^4^ |  | -1.09∙10^4^ |  |
| H_2_ | FeOOH | H_2_O | -1.09∙10^4^ |  | -1.09∙10^4^ |  |
| CH_3_COO^-^ | MnO_2_ | HCO_3_^-^ | 8.49∙10^4^ | -3.43 | 8.95∙10^4^ | 1.79 |
| CH_3_COO^-^ | Fe(OH)_3_ | HCO_3_^-^ | -2.27∙10^4^ |  | -2.27∙10^4^ |  |
| CH_3_COO^-^ | FeOOH | HCO_3_^-^ | -2.26∙10^4^ |  | -2.26∙10^4^ |  |
| NH_4_^+^ | MnO_2_ | NO_3_^-^ | 5.22∙10^5^ | -2.78 | 5.45∙10^5^ | 1.45 |
| NH_4_^+^ | MnO_2_ | NO_2_^-^ | 6.30∙10^5^ | -2.77 | 6.57∙10^5^ | 1.44 |
| NH_4_^+^ | MnO_2_ | N_2_ | 3.62∙10^6^ | -2.76 | 3.77∙10^6^ | 4.82∙10^2^ |
| NH_4_^+^ | Fe(OH)_3_ | N_2_ | 3.60∙10^3^ | -3.98 | 3.83∙10^3^ | -9.99∙10 |
| NH_4_^+^ | FeOOH | N_2_ | 3.05∙10 | -7.46∙10 | 1.66∙10^2^ | -9.56∙10 |

**Table S24:** Biomass growth rates (cell cm^-3^ day^-1^) in SG-1 and change in biomass growth rates relatively to the nominal values, in scenarios where the half-saturation constants of the electron acceptors (*K_s_^EA^*) are 2 times the nominal values (blue-shaded columns) and 0.5 times the nominal values (orange-shaded columns). Negative biomass growth values are marked in red.

| **Electron donor** | **Electron acceptor** | **Product** | **dB/dt for 2∙V_max_** | **Change in dB/dt (%)** | **dB/dt for 0.5∙V_max_** | **Change in dB/dt (%)** |
| --- | --- | --- | --- | --- | --- | --- |
| CH_4_ | MnO_2_ | HCO_3_^-^ | 7.39∙10^6^ | 1.00∙10^2^ | 1.84∙10^6^ | -5.01∙10 |
| CH_4_ | Fe(OH)_3_ | HCO_3_^-^ | 1.19∙10^4^ | 9.74∙10^2^ | -4.28∙10^3^ |  |
| CH_4_ | FeOOH | HCO_3_^-^ | 5.59∙10^3^ | -3.81∙10^2^ | -5.78∙10^3^ |  |
| H_2_ | MnO_2_ | H_2_O | 9.46∙10^6^ | 1.00∙10^2^ | 2.36∙10^6^ | -5.01∙10 |
| H_2_ | Fe(OH)_3_ | H_2_O | 2.16∙10^4^ | 3.67∙10^2^ | -3.87∙10^3^ |  |
| H_2_ | FeOOH | H_2_O | 1.27∙10^4^ | 5.83∙10^3^ | -6.02∙10^3^ |  |
| CH_3_COO^-^ | MnO_2_ | HCO_3_^-^ | 1.89∙10^6^ | 1.03∙10^2^ | 4.55∙10^5^ | -5.13∙10 |
| CH_3_COO^-^ | Fe(OH)_3_ | HCO_3_^-^ | -1.89∙10^4^ |  | -2.42∙10^4^ |  |
| CH_3_COO^-^ | FeOOH | HCO_3_^-^ | -2.05∙10^4^ |  | -2.45∙10^4^ |  |
| NH_4_^+^ | MnO_2_ | NO_3_^-^ | 9.34∙10^5^ | 3.59∙10^20^ | 2.29∙10^5^ | 8.82∙10^19^ |
| NH_4_^+^ | MnO_2_ | NO_2_^-^ | 1.02∙10^6^ | 1.01∙10^2^ | 2.51∙10^5^ | -5.05∙10 |
| NH_4_^+^ | MnO_2_ | N_2_ | 5.23∙10^6^ | 1.00∙10^2^ | 1.30∙10^6^ | -5.02∙10 |
| NH_4_^+^ | Fe(OH)_3_ | N_2_ | 5.79∙10^3^ | -3.84∙10^2^ | -5.95∙10^3^ |  |
| NH_4_^+^ | FeOOH | N_2_ | 1.01∙10^3^ | -1.23∙10^2^ | -7.03∙10^3^ |  |

**Table S25:** Biomass growth rates (cell cm^-3^ day^-1^) in LK and change in biomass growth rates relatively to the nominal values, in scenarios where the *V_max_* values are 2 times the nominal values (blue-shaded columns) and 0.5 times the nominal values (orange-shaded columns). Negative biomass growth values are marked in red.

| **Electron donor** | **Electron acceptor** | **Product** | **dB/dt for 2∙V_max_** | **Change in dB/dt (%)** | **dB/dt for 0.5∙V_max_** | **Change in dB/dt (%)** |
| --- | --- | --- | --- | --- | --- | --- |
| CH_4_ | MnO_2_ | HCO_3_^-^ | 8.19∙10^6^ | 1.00∙10^2^ | 2.04∙10^6^ | -5.02E∙10 |
| CH_4_ | Fe(OH)_3_ | HCO_3_^-^ | 1.07∙10^4^ | -6.91∙10^2^ | -8.10∙10^3^ |  |
| CH_4_ | FeOOH | HCO_3_^-^ | 3.54∙10^3^ | -1.66∙10^2^ | -9.78∙10^3^ |  |
| H_2_ | MnO_2_ | H_2_O | 1.24∙10^4^ | 1.12∙10^3^ | -4.67∙10^3^ |  |
| H_2_ | Fe(OH)_3_ | H_2_O | -1.09∙10^4^ |  | -1.10∙10^4^ |  |
| H_2_ | FeOOH | H_2_O | -1.08∙10^4^ |  | -1.09∙10^4^ |  |
| CH_3_COO^-^ | MnO_2_ | HCO_3_^-^ | 1.98∙10^5^ | 1.25∙10^2^ | 3.30∙10^4^ | -6.24E∙10 |
| CH_3_COO^-^ | Fe(OH)_3_ | HCO_3_^-^ | -2.23∙10^4^ |  | -2.29∙10^4^ |  |
| CH_3_COO^-^ | FeOOH | HCO_3_^-^ | -2.23∙10^4^ |  | -2.28∙10^4^ |  |
| NH_4_^+^ | MnO_2_ | NO_3_^-^ | 1.08∙10^6^ | 1.01∙10^2^ | 2.66∙10^5^ | -5.05∙10 |
| NH_4_^+^ | MnO_2_ | NO_2_^-^ | 1.30∙10^6^ | 1.01∙10^2^ | 3.22∙10^5^ | -5.04∙10 |
| NH_4_^+^ | MnO_2_ | N_2_ | 7.44∙10^6^ | 1.00∙10^2^ | 1.86∙10^6^ | -5.01∙10 |
| NH_4_^+^ | Fe(OH)_3_ | N_2_ | 1.63∙10^4^ | 3.34∙10^2^ | -2.50∙10^3^ |  |
| NH_4_^+^ | FeOOH | N_2_ | 8.87∙10^3^ | 7.27∙10^3^ | -4.25∙10^3^ |  |

**Table S26:** Biomass growth rates (cell cm^-3^ day^-1^) in SG-1 and change in biomass growth rates relatively to the nominal values, in scenarios where the *V_max_* values are 2 times the nominal values (blue-shaded columns) and 0.5 times the nominal values (orange-shaded columns). Negative biomass growth values are marked in red.

| **Electron donor** | **Electron acceptor** | **Product** | **dB/dt LK** | **dB/dt SG-1** |
| --- | --- | --- | --- | --- |
| CH_4_ | MnO_2_ | HCO_3_^-^ | 3.49∙10^6^ | 3.78∙10^6^ |
| CH_4_ | Fe(OH)_3_ | HCO_3_^-^ | -5.95∙10^5^ | -3.31∙10^5^ |
| CH_4_ | FeOOH | HCO_3_^-^ | -5.92∙10^5^ | -3.31∙10^5^ |
| H_2_ | MnO_2_ | H_2_O | 4.46∙10^6^ | -2.36∙10^5^ |
| H_2_ | Fe(OH)_3_ | H_2_O | -2.78∙10^5^ | -2.62∙10^5^ |
| H_2_ | FeOOH | H_2_O | -2.80∙10^5^ | -2.60∙10^5^ |
| CH_3_COO^-^ | MnO_2_ | HCO_3_^-^ | 3.71∙10^5^ | -4.13∙10^5^ |
| CH_3_COO^-^ | Fe(OH)_3_ | HCO_3_^-^ | -6.18∙10^5^ | -5.52∙10^5^ |
| CH_3_COO^-^ | FeOOH | HCO_3_^-^ | -6.14∙10^5^ | -5.47∙10^5^ |
| NH_4_^+^ | MnO_2_ | NO_3_^-^ | 3.38∙10^5^ | 4.25∙10^5^ |
| NH_4_^+^ | MnO_2_ | NO_2_^-^ | 3.90∙10^5^ | 5.43∙10^5^ |
| NH_4_^+^ | MnO_2_ | N_2_ | 2.41∙10^6^ | 3.54∙10^6^ |
| NH_4_^+^ | Fe(OH)_3_ | N_2_ | -2.28∙10^5^ | -1.97∙10^5^ |
| NH_4_^+^ | FeOOH | N_2_ | -2.27∙10^5^ | -1.97∙10^5^ |

**Table S27:** Biomass growth rates (cell cm^-3^ day^-1^) in LK and SG-1, for *P_cs_* values of
8.6∙10^-15^ W cell^-1^. *P_cs_* values of 3.6∙10^-16^ W cell^-1^ were used in the nominal model. Negative biomass growth values are marked in red.

| **Electron donor** | **Electron acceptor** | **Product** | **dB/dt for 2∙ΔG^0’^_cells_** | **Change in dB/dt (%)** | **dB/dt for 0.5∙G^0’^_cells_** | **Change in dB/dt (%)** |
| --- | --- | --- | --- | --- | --- | --- |
| CH_4_ | MnO_2_ | HCO_3_^-^ | 2.60∙10^6^ | -2.96∙10 | 4.53∙10^6^ | 2.26∙10 |
| CH_4_ | Fe(OH)_3_ | HCO_3_^-^ | 7.83∙10^2^ | -2.93∙10 | 1.35∙10^3^ | 2.22∙10 |
| CH_4_ | FeOOH | HCO_3_^-^ | -1.41∙10^3^ |  | -2.43∙10^3^ |  |
| H_2_ | MnO_2_ | H_2_O | 3.77∙10^6^ | -2.02∙10 | 5.34∙10^6^ | 1.31∙10 |
| H_2_ | Fe(OH)_3_ | H_2_O | 3.71∙10^3^ | -2.00∙10 | 5.24∙10^3^ | 1.30∙10 |
| H_2_ | FeOOH | H_2_O | 1.71∙10^2^ | -2.00∙10 | 2.42∙10^2^ | 1.30∙10 |
| CH_3_COO^-^ | MnO_2_ | HCO_3_^-^ | 6.65∙10^5^ | -2.88∙10 | 1.14∙10^6^ | 2.16∙10 |
| CH_3_COO^-^ | Fe(OH)_3_ | HCO_3_^-^ | -1.61∙10^4^ |  | -2.72∙10^4^ |  |
| CH_3_COO^-^ | FeOOH | HCO_3_^-^ | -1.65∙10^4^ |  | -2.81∙10^4^ |  |
| NH_4_^+^ | MnO_2_ | NO_3_^-^ | 3.63∙10^5^ | -2.19∙10 | 5.32∙10^5^ | 1.46∙10 |
| NH_4_^+^ | MnO_2_ | NO_2_^-^ | 3.97∙10^5^ | -2.19∙10 | 5.82∙10^5^ | 1.46∙10 |
| NH_4_^+^ | MnO_2_ | N_2_ | 2.06∙10^6^ | -2.10∙10 | 2.97∙10^6^ | 1.38∙10 |
| NH_4_^+^ | Fe(OH)_3_ | N_2_ | -1.61∙10^3^ |  | -2.31∙10^3^ |  |
| NH_4_^+^ | FeOOH | N_2_ | -3.45∙10^3^ |  | -4.94∙10^3^ |  |

**Table S28:** Biomass growth rates (cell cm^-3^ day^-1^) in LK and change in biomass growth rates relatively to the nominal values, in scenarios where the *ΔG^0’^_cells_* values are 2 times the nominal values (blue-shaded columns) and 0.5 times the nominal values (orange-shaded columns). Negative biomass growth values are marked in red.

| **Electron donor** | **Electron acceptor** | **Product** | **dB/dt for 2∙ΔG^0’^_cells_** | **Change in dB/dt (%)** | **dB/dt for 0.5∙ΔG^0’^_cells_** | **Change in dB/dt (%)** |
| --- | --- | --- | --- | --- | --- | --- |
| CH_4_ | MnO_2_ | HCO_3_^-^ | 2.87∙10^6^ | -2.99∙10 | 5.01∙10^6^ | 2.26∙10 |
| CH_4_ | Fe(OH)_3_ | HCO_3_^-^ | -1.15∙10^4^ |  | -2.22∙10^3^ |  |
| CH_4_ | FeOOH | HCO_3_^-^ | -1.38∙10^4^ |  | -6.53∙10^3^ |  |
| H_2_ | MnO_2_ | H_2_O | -7.45∙10^3^ |  | 1.15∙10^3^ | 1.31∙10 |
| H_2_ | Fe(OH)_3_ | H_2_O | -1.75∙10^4^ |  | -1.24∙10^4^ |  |
| H_2_ | FeOOH | H_2_O | -1.74∙10^4^ |  | -1.23∙10^4^ |  |
| CH_3_COO^-^ | MnO_2_ | HCO_3_^-^ | 4.70∙10^4^ | -4.65∙10 | 1.07∙10^5^ | 2.16∙10 |
| CH_3_COO^-^ | Fe(OH)_3_ | HCO_3_^-^ | -3.28∙10^4^ |  | -2.75∙10^4^ |  |
| CH_3_COO^-^ | FeOOH | HCO_3_^-^ | -3.25∙10^4^ |  | -2.74∙10^4^ |  |
| NH_4_^+^ | MnO_2_ | NO_3_^-^ | 4.16∙10^5^ | -2.26∙10 | 6.15∙10^5^ | 1.46∙10 |
| NH_4_^+^ | MnO_2_ | NO_2_^-^ | 2.93∙10^6^ | -2.11∙10 | 4.23∙10^6^ | 1.38∙10 |
| NH_4_^+^ | MnO_2_ | N_2_ | -3.97∙10^3^ |  | 4.26∙10^3^ | 1.36∙10 |
| NH_4_^+^ | Fe(OH)_3_ | N_2_ | -6.74∙10^3^ |  | 1.37∙10^2^ | 1.36∙10 |
| NH_4_^+^ | FeOOH | N_2_ | -6.74∙10^3^ |  | 1.37∙10^2^ | 1.36∙10 |

**Table S29:** Biomass growth rates (cell cm^-3^ day^-1^) in SG-1 and change in biomass growth rates relatively to the nominal values, in scenarios where the *ΔG^0’^_cells_* values are 2 times the nominal values (blue-shaded columns) and 0.5 times the nominal values (orange-shaded columns). Negative biomass growth values are marked in red.

| **Electron donor** | **Electron acceptor** | **Product** | **dB/dt for 2∙ϵ** | **Change in dB/dt (%)** | **dB/dt for 0.5∙ϵ** | **Change in dB/dt (%)** |
| --- | --- | --- | --- | --- | --- | --- |
| CH_4_ | MnO_2_ | HCO_3_^-^ | 1.07∙10^7^ | 1.90∙10^2^ | 1.14∙10^6^ | -6.92∙10 |
| CH_4_ | Fe(OH)_3_ | HCO_3_^-^ | 3.15∙10^3^ | 1.85∙10^2^ | 3.47∙10^2^ | -6.87∙10 |
| CH_4_ | FeOOH | HCO_3_^-^ | -5.69∙10^3^ |  | -6.22∙10^2^ |  |
| H_2_ | MnO_2_ | H_2_O | 1.44∙10^7^ | 2.05∙10^2^ | 1.40∙10^6^ | -7.04∙10 |
| H_2_ | Fe(OH)_3_ | H_2_O | 1.39∙10^4^ | 2.00∙10^2^ | 1.39∙10^3^ | -7.00∙10 |
| H_2_ | FeOOH | H_2_O | 6.44∙10^2^ | 2.01∙10^2^ | 6.39∙10 | -7.01∙10 |
| CH_3_COO^-^ | MnO_2_ | HCO_3_^-^ | 2.67∙10^6^ | 1.85∙10^2^ | 2.92∙10^5^ | -6.87∙10 |
| CH_3_COO^-^ | Fe(OH)_3_ | HCO_3_^-^ | -6.30∙10^4^ |  | -7.13∙10^3^ |  |
| CH_3_COO^-^ | FeOOH | HCO_3_^-^ | -6.52∙10^4^ |  | -7.33∙10^3^ |  |
| NH_4_^+^ | MnO_2_ | NO_3_^-^ | 1.62∙10^6^ | 2.49∙10^2^ | 1.25∙10^5^ | -7.30∙10 |
| NH_4_^+^ | MnO_2_ | NO_2_^-^ | 1.79∙10^6^ | 2.51∙10^2^ | 1.36∙10^5^ | -7.31∙10 |
| NH_4_^+^ | MnO_2_ | N_2_ | 8.45∙10^6^ | 2.24∙10^2^ | 7.39∙10^5^ | -7.17∙10 |
| NH_4_^+^ | Fe(OH)_3_ | N_2_ | -6.44∙10^3^ |  | -5.86∙10^2^ |  |
| NH_4_^+^ | FeOOH | N_2_ | -1.38∙10^4^ |  | -1.25∙10^3^ |  |

**Table S30:** Biomass growth rates (cell cm^-3^ day^-1^) in LK and change in biomass growth rates relatively to the nominal values, in scenarios where the *ϵ* values are 2 times the nominal values (blue-shaded columns) and 0.5 times the nominal values (orange-shaded columns). Negative biomass growth values are marked in red.

| **Electron donor** | **Electron acceptor** | **Product** | **dB/dt for 2∙ϵ** | **Change in dB/dt (%)** | **dB/dt for 0.5∙ϵ** | **Change in dB/dt (%)** |
| --- | --- | --- | --- | --- | --- | --- |
| CH_4_ | MnO_2_ | HCO_3_^-^ | 1.19∙10^7^ | 1.90∙10^2^ | 1.26∙10^6^ | -6.92∙10 |
| CH_4_ | Fe(OH)_3_ | HCO_3_^-^ | -5.18∙10^3^ |  | -5.70∙10^2^ |  |
| CH_4_ | FeOOH | HCO_3_^-^ | -1.53∙10^4^ |  | -1.67∙10^3^ |  |
| H_2_ | MnO_2_ | H_2_O | 3.11∙10^3^ | 2.05∙10^2^ | 3.02∙10^2^ | -7.04∙10 |
| H_2_ | Fe(OH)_3_ | H_2_O | -3.29∙10^4^ |  | -3.28∙10^3^ |  |
| H_2_ | FeOOH | H_2_O | -3.27∙10^4^ |  | -3.25∙10^3^ |  |
| CH_3_COO^-^ | MnO_2_ | HCO_3_^-^ | 2.51∙10^5^ | 1.85∙10^2^ | 2.75∙10^4^ | -6.87∙10 |
| CH_3_COO^-^ | Fe(OH)_3_ | HCO_3_^-^ | -6.37∙10^4^ |  | -7.21∙10^3^ |  |
| CH_3_COO^-^ | FeOOH | HCO_3_^-^ | -6.36∙10^4^ |  | -7.16∙10^3^ |  |
| NH_4_^+^ | MnO_2_ | NO_3_^-^ | 1.87∙10^6^ | 2.49∙10^2^ | 1.45∙10^5^ | -7.30∙10 |
| NH_4_^+^ | MnO_2_ | NO_2_^-^ | 2.28∙10^6^ | 2.51∙10^2^ | 1.74∙10^5^ | -7.31∙10 |
| NH_4_^+^ | MnO_2_ | N_2_ | 1.20∙10^7^ | 2.24∙10^2^ | 1.05∙10^6^ | -7.17∙10 |
| NH_4_^+^ | Fe(OH)_3_ | N_2_ | 1.19∙10^4^ | 2.16∙10^2^ | 1.08∙10^3^ | -7.12∙10 |
| NH_4_^+^ | FeOOH | N_2_ | 3.82∙10^2^ | 2.17∙10^2^ | 3.46∙10 | -7.13∙10 |

**Table S31:** Biomass growth rates (cell cm^-3^ day^-1^) in SG-1 and change in biomass growth rates relatively to the nominal values, in scenarios where the *ϵ* values are 2 times the nominal values (blue-shaded columns) and 0.5 times the nominal values (orange-shaded columns). Negative biomass growth values are marked in red.

| **Electron donor** | **Electron acceptor** | **Product** | **dB/dt for 2∙K** | **Change in dB/dt (%)** | **dB/dt for 0.5∙K** | **Change in dB/dt (%)** |
| --- | --- | --- | --- | --- | --- | --- |
| CH_4_ | MnO_2_ | HCO_3_^-^ | 5.36∙10^6^ | 4.51∙10 | 2.28∙10^6^ | -3.83∙10 |
| CH_4_ | Fe(OH)_3_ | HCO_3_^-^ | 1.58∙10^3^ | 4.24∙10 | 6.94∙10^2^ | -3.73∙10 |
| CH_4_ | FeOOH | HCO_3_^-^ | -2.84∙10^3^ |  | -1.24∙10^3^ |  |
| H_2_ | MnO_2_ | H_2_O | 7.20∙10^6^ | 5.24∙10 | 2.80∙10^6^ | -4.07∙10 |
| H_2_ | Fe(OH)_3_ | H_2_O | 6.97∙10^3^ | 5.02∙10 | 2.78∙10^3^ | -4.01∙10 |
| H_2_ | FeOOH | H_2_O | 3.22∙10^2^ | 5.06∙10 | 1.28∙10^2^ | -4.02∙10 |
| CH_3_COO^-^ | MnO_2_ | HCO_3_^-^ | 1.33∙10^6^ | 4.27∙10 | 5.84∙10^5^ | -3.74∙10 |
| CH_3_COO^-^ | Fe(OH)_3_ | HCO_3_^-^ | -3.15∙10^4^ |  | -1.43∙10^4^ |  |
| CH_3_COO^-^ | FeOOH | HCO_3_^-^ | -3.26∙10^4^ |  | -1.47∙10^4^ |  |
| NH_4_^+^ | MnO_2_ | NO_3_^-^ | 8.09∙10^5^ | 7.43∙10 | 2.51∙10^5^ | -4.60∙10 |
| NH_4_^+^ | MnO_2_ | NO_2_^-^ | 8.93∙10^5^ | 7.57∙10 | 2.73∙10^5^ | -4.63∙10 |
| NH_4_^+^ | MnO_2_ | N_2_ | 4.22∙10^6^ | 6.19∙10 | 1.48∙10^6^ | -4.33∙10 |
| NH_4_^+^ | Fe(OH)_3_ | N_2_ | -3.22∙10^3^ |  | -1.17∙10^3^ |  |
| NH_4_^+^ | FeOOH | N_2_ | -6.90∙10^3^ |  | -2.50∙10^3^ |  |

**Table S32:** Biomass growth rates (cell cm^-3^ day^-1^) in LK and change in biomass growth rates relatively to the nominal values, in scenarios where the *K* values are 2 times the nominal values (blue-shaded columns) and 0.5 times the nominal values (orange-shaded columns). Negative biomass growth values are marked in red.

| **Electron donor** | **Electron acceptor** | **Product** | **dB/dt for 2∙K** | **Change in dB/dt (%)** | **dB/dt for 0.5∙K** | **Change in dB/dt (%)** |
| --- | --- | --- | --- | --- | --- | --- |
| CH_4_ | MnO_2_ | HCO_3_^-^ | 5.93∙10^6^ | 4.51∙10 | 2.52∙10^6^ | -3.83∙10 |
| CH_4_ | Fe(OH)_3_ | HCO_3_^-^ | -2.59∙10^3^ |  | -1.14∙10^3^ |  |
| CH_4_ | FeOOH | HCO_3_^-^ | -7.63∙10^3^ |  | -3.34∙10^3^ |  |
| H_2_ | MnO_2_ | H_2_O | 1.55∙10^3^ | 5.24∙10 | 6.04∙10^2^ | -4.07∙10 |
| H_2_ | Fe(OH)_3_ | H_2_O | -1.64∙10^4^ |  | -6.56∙10^3^ |  |
| H_2_ | FeOOH | H_2_O | -1.63∙10^4^ |  | -6.49∙10^3^ |  |
| CH_3_COO^-^ | MnO_2_ | HCO_3_^-^ | 1.25∙10^5^ | 4.27∙10 | 5.50∙10^4^ | -3.74∙10 |
| CH_3_COO^-^ | Fe(OH)_3_ | HCO_3_^-^ | -3.19∙10^4^ |  | -1.44∙10^4^ |  |
| CH_3_COO^-^ | FeOOH | HCO_3_^-^ | -3.18∙10^4^ |  | -1.43∙10^4^ |  |
| NH_4_^+^ | MnO_2_ | NO_3_^-^ | 9.36∙10^5^ | 7.43∙10 | 2.90∙10^5^ | -4.60∙10 |
| NH_4_^+^ | MnO_2_ | NO_2_^-^ | 1.14∙10^6^ | 7.57∙10 | 3.48∙10^5^ | -4.63∙10 |
| NH_4_^+^ | MnO_2_ | N_2_ | 6.02∙10^6^ | 6.19∙10 | 2.11∙10^6^ | -4.33∙10 |
| NH_4_^+^ | Fe(OH)_3_ | N_2_ | 5.93∙10^3^ | 5.82∙10 | 2.16∙10^3^ | -4.24∙10 |
| NH_4_^+^ | FeOOH | N_2_ | 1.91∙10^2^ | 5.87∙10 | 6.92∙10 | -4.25∙10 |

**Table S33:** Biomass growth rates (cell cm^-3^ day^-1^) in SG-1 and change in biomass growth rates relatively to the nominal values, in scenarios where the *K* values are 2 times the nominal values (blue-shaded columns) and 0.5 times the nominal values (orange-shaded columns). Negative biomass growth values are marked in red.

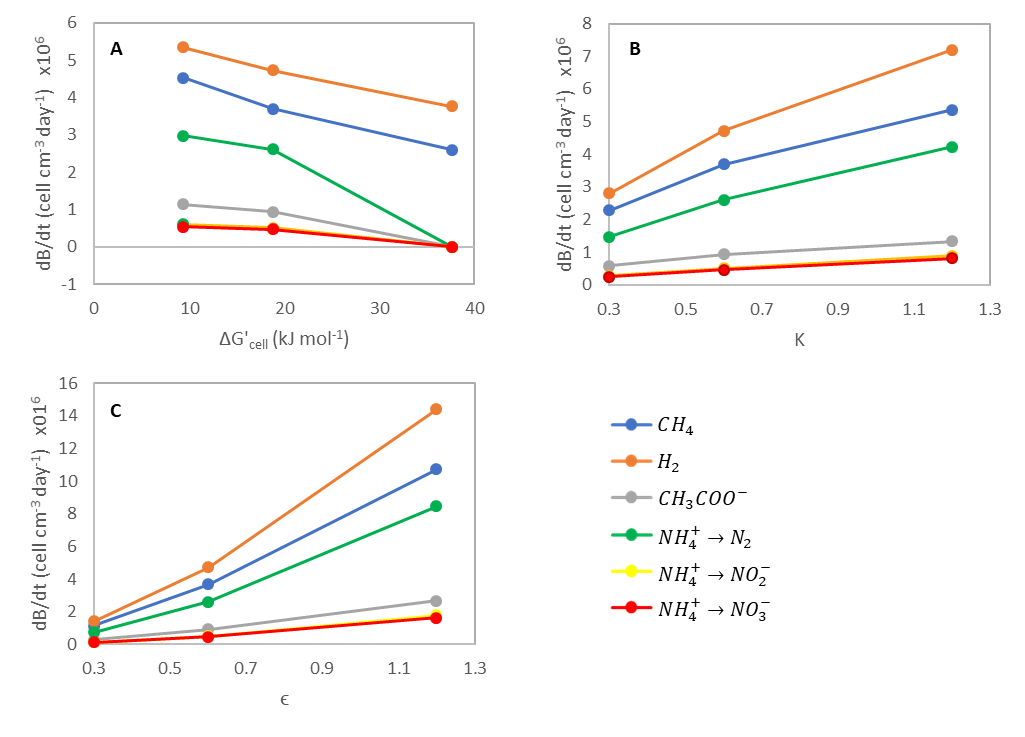
**Figure S14:** Biomass growth rates versus *ΔG’_cells_*, *K*, and *ϵ*, in LK for nominal parameter values, and for parameter values of 2 times and 0.5 times the nominal values. The electron acceptor in all bioreactions is MnO_2_ and the electron donors are listed in the legend.

13. **Distributions and standard deviations of model parameters**

| **Parameter** | **Distribution** | **Standard Deviation** | **Notes** |
| --- | --- | --- | --- |
| ΔG'_cells_ | Normal | 2.17·10^-1^ | All reactions |
| K | Normal | 5.20·10^-2^ | All reactions |
| E | Normal | 5.20·10^-2^ | All reactions |
| P_cs_ | Normal | 2.69·10^-15^ | All reactions |
| B(archaea) | Normal | 3.14·10^5^ | All reactions |
| B(bacteria) | Normal | 5.08·10^-5^ | All reactions |
| r_cell_ | Log-normal | 2.20·10^-1^ | All reactions |
| V_max_ | Log-normal | 2.20·10^-1^ | All reactions |
| K_S_^ED^ | Log-normal | 2.20·10^-1^ | All reactions |
| K_S_^EA^ | Log-normal | 2.20·10^-1^ | All reactions |
| [CH_4_] | Log-normal | 5.50·10^-1^ | Only in LK |
| [H_2_] | Log-normal | 5.50·10^-1^ | Only in LK |
| [CH_4_] | Log-normal | 2.20·10^-1^ | Only in the MedS |
| [H_2_] | Log-normal | 2.20·10^-1^ | Only in the MedS |
| [CH_3_COO^-^] | Log-normal | 2.20·10^-1^ | In Lk and the MedS |
| [NH_4_^+^] | Log-normal | 2.20·10^-1^ | In Lk and the MedS |

**Table S34:** Distribution and standard deviations of model parameters chosen for the Monte Carlo Simulation.

14. **Sediment properties**

| **Property** | **LK** 9.5 cm BSWI | **Reference** | **MedS** 167 cm BSWI | **Reference** |
| --- | --- | --- | --- | --- |
| Φ – porosity | 0.8 | Vigderovich et al. (unpublished results) | 0.48 | Vigderovich et al. (unpublished results) |
| ω – sedimentation rate (cm/sec) | 1.27∙10^-8^ | Adler et al. (2011) | 9.51∙10^-9^ | Zemach et al. (unpublished results) |
| ρ_ds_ – density of dry sediment (gr/cm^3^) | 1.5 | Assumption | 1.5 | Assumption |
| D_0,SO4_ – diffusion coefficient of sulfate in water (cm^2^/sec) | 7.90∙10^-6^ | After Lerman (1979) | 9.92∙10^-6^ | Berner (1980) |

**Table S35:** Sediment properties above the methanogenic zones of LK and the MedS.

| **Environment** | **Depth** cm | **[SO_4_^2-^]** | **[AVS]** | **[CRS]** | **[Oxides 1]** | **Reference** |
| --- | --- | --- | --- | --- | --- | --- |
| LK | 7 | 5.00∙10^-8^ |  |  |  | Adler et al. (2011) |
|  | 9 |  | 3.00∙10^-8^ | 4.76∙10^-13^ |  | Kamyshny et al. (unpublished results) |
|  | 9.5 |  |  |  | 2.82∙10^-5^ | Kamyshny et al. (unpublished results) |
|  | 11 | 0 |  |  |  | Adler et al. (2011) |
| SG-1 | 50 |  | 4.19∙10^-5^ | 3.47∙10^-5^ |  | Bosco-Santos et al. (unpublished results) |
|  | 145 |  |  |  | 6.22∙10^-5^ | Vigderovich et al. (2019) |
|  | 150 | 7 |  |  |  | Vigderovich et al. (2019) |
|  | 200 | 0 |  |  |  | Vigderovich et al. (2019) |

**Table S36:** Concentrations of sulfate (mol cm^-3^ porewater), AVS, CRS and amorphous/poorly crystalline iron oxides (Oxides 1) (mol/cm^3^ total sediment) above the methanogenic zones of LK and the MedS (SG-1).
